# Transcriptional patterns of sexual dimorphism and in host developmental programs in the model parasitic nematode *Heligmosomoides bakeri*

**DOI:** 10.1101/2022.09.14.508015

**Authors:** Stephen M. J. Pollo, Aralia Leon-Coria, Hongrui Liu, Davide Cruces Gonzalez, Constance A. M. Finney, James D. Wasmuth

**Affiliations:** Faculty of Veterinary Medicine, University of Calgary, Calgary, Alberta, Canada; Host-Parasite Interactions Research Training Network, University of Calgary, Calgary, Alberta, Canada; Department of Biological Sciences, Faculty of Science, University of Calgary, Calgary, Alberta, Canada

## Abstract

*Heligmosomoides bakeri* (often mistaken for *Heligmosomoides polygyrus*) is a promising model for parasitic nematodes with the key advantage of being amenable to study and manipulation within a controlled laboratory environment. While draft genome sequences are available for this worm, which allow for comparative genomic analyses between nematodes, there is a notable lack of information on its gene expression. Here, we have generated biologically replicated RNA-seq datasets from samples taken throughout the parasitic life of *H. bakeri*. We find extensive transcriptional sexual dimorphism throughout the fourth larval and adult stages of this parasite and identify alternative splicing, glycosylation, and ubiquitination as particularly important processes for establishing and/or maintaining sex-specific gene expression in this species. Further, we find sex-linked differences in transcription related to aging and oxidative and osmotic stress responses. Additionally, we observe a starvation-like signature among transcripts whose expression is consistently up-regulated in males, which may reflect a higher energy expenditure by male worms. We detect evidence of increased importance for anaerobic respiration among the adult worms, which coincides with the parasite’s migration into the physiologically hypoxic environment of the intestinal lumen. Further, we hypothesize that oxygen concentration may be an important driver of the worms encysting in the intestinal mucosa as larvae, which not only fully exposes the worms to their host’s immune system, but also shapes many of the interactions between the host and parasite. We find stage- and sex-specific variation in the expression of immunomodulatory genes and in anthelmintic targets. In addition to generating new hypotheses for follow-up experiments into the worm’s behaviour, physiology, and metabolism, our datasets enable future more in-depth comparisons between nematodes to better define the utility of *H. bakeri* as a model for parasitic nematodes in general.

**Author Summary:** Parasitic nematodes (roundworms) that infect humans and livestock are a major health and economic burden but are challenging to study in a laboratory environment because of their required hosts. One strategy to get around this difficulty is to first study a rodent model to guide targeted experiments in the more difficult study system. *Heligmosomoides bakeri* is closely related to the nematode parasites of humans and livestock and naturally parasitizes mice. We have generated information on the expression of all the genes in this worm throughout the stages of its life when it is parasitic. This information allows us to examine how different the male and female worms are at the molecular level. We also describe major developmental events that occur in the worm, which extend our understanding of the interactions between this parasite and its host. We analyse the expression of genes known to be involved in interfering with host immune responses and others known to be targeted by drugs designed to kill worms. This new information will allow for better comparisons among nematodes to assess how well this rodent model system works for studying parasitic nematodes in general.

## Introduction

Parasitic nematodes (roundworms) infect over 1 billion people, causing significant morbidity [1]. They also pose a significant threat to livestock, costing billions of dollars annually in production losses and treatment costs [2,3]. Yet, nematodes that parasitize humans or livestock are challenging to study in a controlled laboratory environment because the host is required to complete the parasite lifecycle. Studying the *in vivo* behaviour of the parasite during its infection requires samples to be taken from an infected host, not just obtaining eggs from feces. These samples can be technically challenging and expensive to obtain from large animals and not possible to obtain from humans. *Heligmosomoides bakeri* is well-suited to being a laboratory model because it is a natural parasite of mice, which are easily maintained in a laboratory environment, and there are protocols for maintaining *H. bakeri* in a laboratory environment. It is closely related to economically important parasites of livestock and the hookworm parasites of humans [4,5]. Furthermore, it can establish chronic infections in its mouse host, coarsely mimicking many other parasitic infections in a variety of hosts. Unravelling the responses to infection, adaptations to parasitism, and immunomodulatory strategies of *H. bakeri* will enable further development of this worm as a model for parasitic nematodes. Increased understanding of the biology of *H. bakeri* and what can be generalized to other related nematodes will then also enable preliminary experiments in *H. bakeri* to inform targeted experiments in harder to study parasites of large animals.

There has been an ongoing discussion as to the species designation for the lab isolates. Much of the literature on *H. bakeri* refers to the species as *H. polygyrus*, and a third name, *Nematospiroides dubius* has also been used. However, molecular studies demonstrate that the isolates widely used in laboratories are sufficiently genetically different to justify the original two species designation [6–8]. We confirm the species designation of the isolate used in this work (see Results). We, therefore, refer to the laboratory strain of this parasite as *H. bakeri*.

*H. bakeri* has a direct lifecycle with free-living and parasitic stages (Fig 1). Eggs present in feces hatch after 36-37 hours, releasing the L1 larvae which molt to L2 larvae 28-29 hours after hatching. After another 17-20 hours, the larvae partially molt resulting in ensheathed L3 worms that are the infective stage of the parasite. Once eaten by a rodent host, the L3 larvae exsheath and within 24 hours have penetrated the intestinal mucosa. The L3 larvae molt 90-96 hours (∼4 days) after infection into L4 larvae which continue to develop in a granuloma formed by the host. A final molt 144-166 hours (∼7 days) after infection results in adults which migrate out to the intestinal lumen [9]. The adult worms coil around the villi where they feed, mate, and lay the eggs which get passed with the feces [10]. The adults reside in the small intestine until they are expelled by the mouse, which can take anywhere from four to greater than 20 weeks depending on the strain of mouse [11].

**Fig 1.**
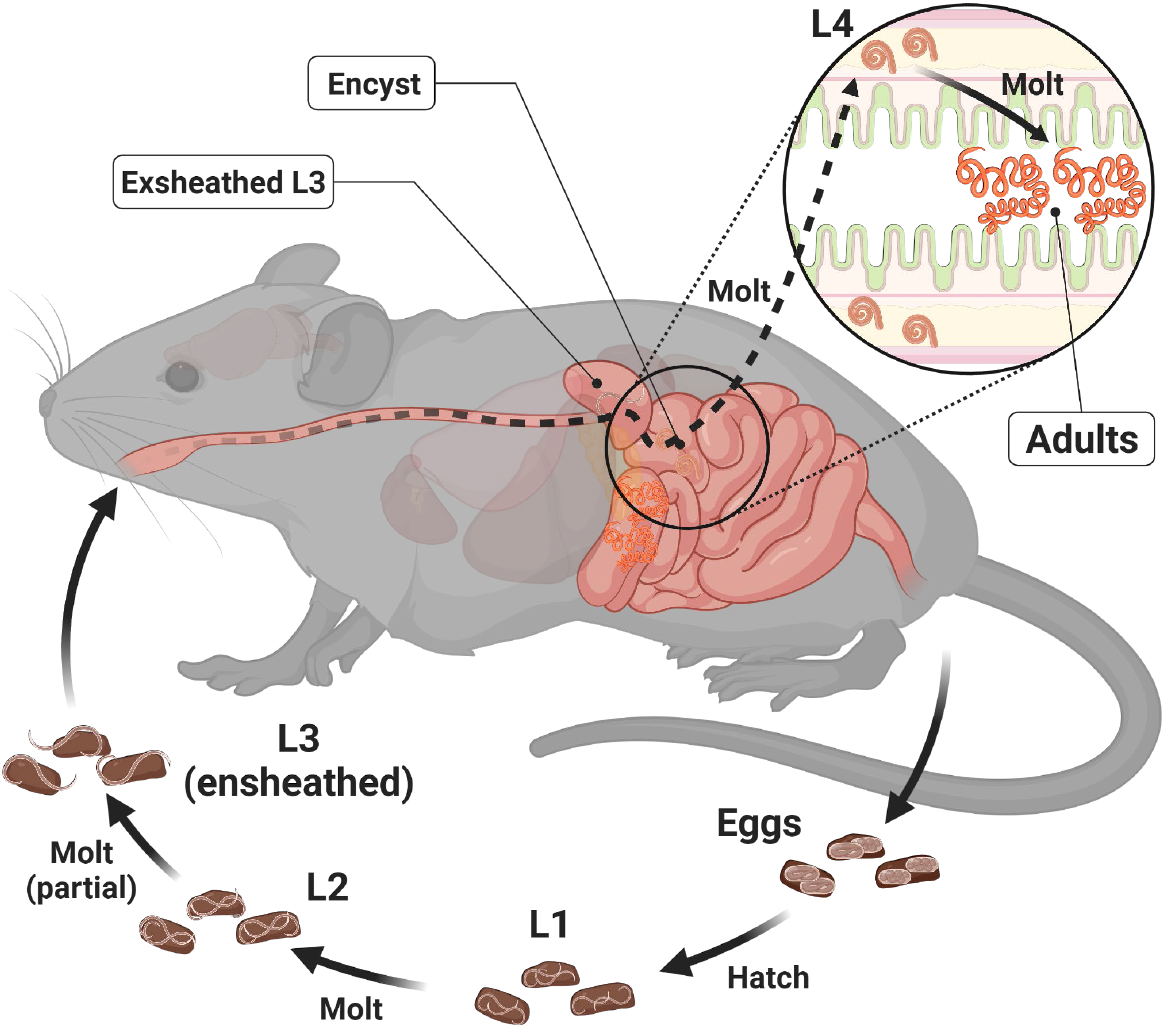
Life cycle of *H. bakeri*. Eggs present in mouse feces develop through two larval stages and arrest during their infective L3 stage. Upon being eaten by a mouse host the L3s exsheath in the stomach and upper duodenum and burrow into the duodenum tissue to form a granuloma. Development continues until the final molt to the adult form when the worms migrate to the lumen of the duodenum where they mate and lay their eggs that get passed with the feces.

Most studies on *H. bakeri* are motivated to understand its effects on its host’s immune system; comparatively little is known about the worm itself. Two draft genome assemblies [12,13] are available for *H. bakeri*, which not only enable comparative genomic studies, but also allow for explorations of gene expression. A small set of 52 genes in *H. bakeri* (including chitinase, lysozyme, and glutathione S-transferases) was found to be differentially expressed between germ-free and pathogen-free mice using bulk RNA-seq of mixed-sex cultures, suggesting sensing of the microbial environment by the adult worms [14]. However, no transcriptomic information exists for other stages throughout the lifecycle or for sex-separated adults. Finally, attempts have been made to identify excreted/secreted products from the worm that may have immunomodulatory potential, such as miRNAs within exosomes [15] and proteins from separated and mixed-sex cultures [16].

Here, we have generated biologically replicated mRNA transcriptomes for both male and female *H. bakeri* at four timepoints throughout their infection using RNA-sequencing (RNA-seq). The samples span the fourth larval and adult stages of the worm (5, 7, 10, and 21 days post-infection). Our dataset allows us to examine, for the first time, sexual dimorphism in the L4s and adults of this species at the level of gene transcription. It also enables us to describe the major developmental changes and processes that occur in the worms during their parasitic phase. Moreover, this dataset provides the first publicly available resource to query the overall level of transcription of any gene of interest in this species throughout the parasitic stages.

## Results and Discussion

### Confirming source species as *Heligmosomoides bakeri*

We mapped a randomly selected ten million RNA reads to the genome assemblies of *H. bakeri* and *H. polygyrus*, each derived from single worms. We found that 93% of the reads mapped to *H. bakeri* with 0.5% mismatch rate per base and 89% of the reads mapped to *H. polygyrus* with 2% mismatch rate per base. When considering the alignment scores, 78% of reads mapped to better to *H. bakeri*, 8% mapped better to *H. polygyrus*, and 14% mapped equally well to both. Collectively these demonstrate that our nematode lab isolate is *H. bakeri*.

### Mapping of bulk RNA-seq data and Differential Gene Expression (DGE)

Using the splice-aware aligner STAR, we mapped the RNA-seq reads to the *H. bakeri* genome assembly obtained from WormBase ParaSite (PRJEB15396). Among all the datasets, 93.26 – 95.62% of the reads uniquely mapped to the reference genome (Table 1), reflecting the high quality of the RNA datasets (See S1 Text for additional quality control discussion). Moreover, the high mapping rate (which does not include chimeric alignments) also indicates that the genome, while highly fragmented in 23647 scaffolds, contains the vast majority of the transcribed polyadenylated RNAs.

**Table 1.** Metadata of the RNA-seq samples in this study.

| Sample Name | File Name | # worms extracted | Days Post Infection | RIN | Reads Uniquely Mapped / Total Reads (%) | Included in Pairwise Comparisons? |
| --- | --- | --- | --- | --- | --- | --- |
| D5 F2 | D5-F2_Li29408_S5 | 174 | 5 | 9.6 | 112169129 / 118520168 (94.64) | Yes |
| D5 F3 | D5-F3_Li29409_S6 | 297 | 5 | 9.4 | 129241490 / 136943707 (94.38) | Yes |
| D5 F4 | D5-F4_Li29410_S7 | 173 | 5 | 9.3 | 107651456 / 113590373 (94.77) | Yes |
| D5 F5 | D5-F5_Li29411_S8 | 232 | 5 | 9.5 | 118950175 / 125198338 (95.01) | Yes |
| D5 M1 | D5-M1_Li29404_S1 | 132 | 5 | 9.3 | 128308873 / 135898387 (94.42) | Yes |
| D5 M2 | D5-M2_Li29405_S2 | 100 | 5 | 9.5 | 124417881 / 131400648 (94.69) | Yes |
| D5 M4 | D5-M4_Li29406_S3 | 95 | 5 | 9.6 | 138697190 / 146832868 (94.46) | Yes |
| D5 M5 | D5-M5_Li29407_S4 | 179 | 5 | 9.6 | 134892659 / 142089783 (94.93) | Yes |
| D7 F1 | D7-F1_Li29416_S13 | 150 | 7 | 9.4 | 108668904 / 114523342 (94.89) | Yes |
| D7 F4 | D7-F4_Li29417_S14 | 211 | 7 | 9.2 | 141571706 / 149554981 (94.66) | Yes |
| D7 F5 | D7-F5_Li29418_S15 | 120 | 7 | 9.4 | 131197468 / 138924573 (94.44) | Yes |
| D7 F6 | D7-F6_Li29419_S16 | 205 | 7 | 9.4 | 144299969 / 153229819 (94.17) | Yes |
| D7 M1 | D7-M1_Li29412_S9 | 126 | 7 | 9.5 | 135646150 / 143790919 (94.34) | Yes |
| D7 M2 | D7-M2_Li29413_S10 | 89 | 7 | 9.3 | 138636801 / 147210080 (94.18) | Yes |
| D7 M4 | D7-M4_Li29414_S11 | 144 | 7 | 9.6 | 113177165 / 119922840 (94.37) | Yes |
| D7 M7 | D7-M7_Li29415_S12 | 85 | 7 | 9.7 | 117210623 / 124680285 (94.01) | Yes |
| D10 F1 | D10-F1_Li29424_S21 | 100 | 10 | 9.6 | 126048861 / 132664380 (95.01) | Yes |
| D10 F2 | D10-F2_Li29425_S22 | 100 | 10 | 9.9 | 117253862 / 123536616 (94.91) | Yes |
| D10 F6 | D10-F6_Li29426_S23 | 80 | 10 | 9.7 | 128203029 / 136497304 (93.92) | No |
| D10 F7 | D10-F7_Li29427_S24 | 80 | 10 | 9.8 | 130193841 / 138092543 (94.28) | No |
| D10 F8 | D10-F8_Li31315_S1 | 80 | 10 | 9.4 | 166132630 / 173733692 (95.62) | Yes |
| D10 F9 | D10-F9_Li31316_S2 | 80 | 10 | 9.2 | 196647784 / 205996448 (95.46) | Yes |
| D10 F10 | D10-F10_Li31317_S3 | 80 | 10 | 9.3 | 170031664 / 178491143 (95.26) | Yes |
| D10 F11 | D10-F11_Li31318_S4 | 80 | 10 | 9.2 | 162920697 / 170951647 (95.30) | Yes |
| D10 F13 | D10-F13_Li31319_S5 | 80 | 10 | 9.5 | 147832599 / 155432294 (95.11) | Yes |
| D10 M1 | D10-M1_Li29420_S17 | 100 | 10 | 9.4 | 135805443 / 145141041 (93.57) | Yes |
| D10 M2 | D10-M2_Li29421_S18 | 100 | 10 | 9.6 | 128421348 / 137123098 (93.65) | Yes |
| D10 M3 | D10-M3_Li29422_S19 | 80 | 10 | 9.5 | 130675067 / 138649879 (94.25) | Yes |
| D10 M6 | D10-M6_Li29423_S20 | 80 | 10 | 9.8 | 110098546 / 116957271 (94.14) | Yes |
| D21 F5 | D21-F5_Li29432_S29 | 80 | 21 | 9.1 | 108798485 / 114364046 (95.13) | Yes |
| D21 F6 | D21-F6_Li29433_S30 | 80 | 21 | 9.5 | 100100633 / 104891911 (95.43) | Yes |
| D21 F11 | D21-F11_Li29434_S31 | 80 | 21 | 9.4 | 98844233 / 103850834 (95.18) | Yes |
| D21 F12 | D21-F12_Li29435_S32 | 80 | 21 | 4.9 | 100561928 / 105898940 (94.96) | Yes |
| D21 M5 | D21-M5_Li29428_S25 | 80 | 21 | 9 | 114067372 / 122106224 (93.42) | Yes |
| D21 M7 | D21-M7_Li29429_S26 | 80 | 21 | 8.7 | 122812424 / 131448504 (93.43) | Yes |
| D21 M12 | D21-M12_Li29430_S27 | 80 | 21 | 9.2 | 122775218 / 130863303 (93.82) | Yes |
| D21 M13 | D21-M13_Li29431_S28 | 80 | 21 | 9.1 | 104773522 / 112342173 (93.26) | Yes |

The sample groups were compared to each other using DESeq2 to find all transcripts that are statistically significantly differentially expressed between conditions. The numbers of transcripts up- and down-regulated between age-matched females and males or between adjacent timepoints among females or males are shown in Fig 2. In general, differences between males and females increase with age, whereas the biggest developmental differences are between the D7 and D10 timepoints for females and the D5 and D7 timepoints for males.

**Fig 2.**
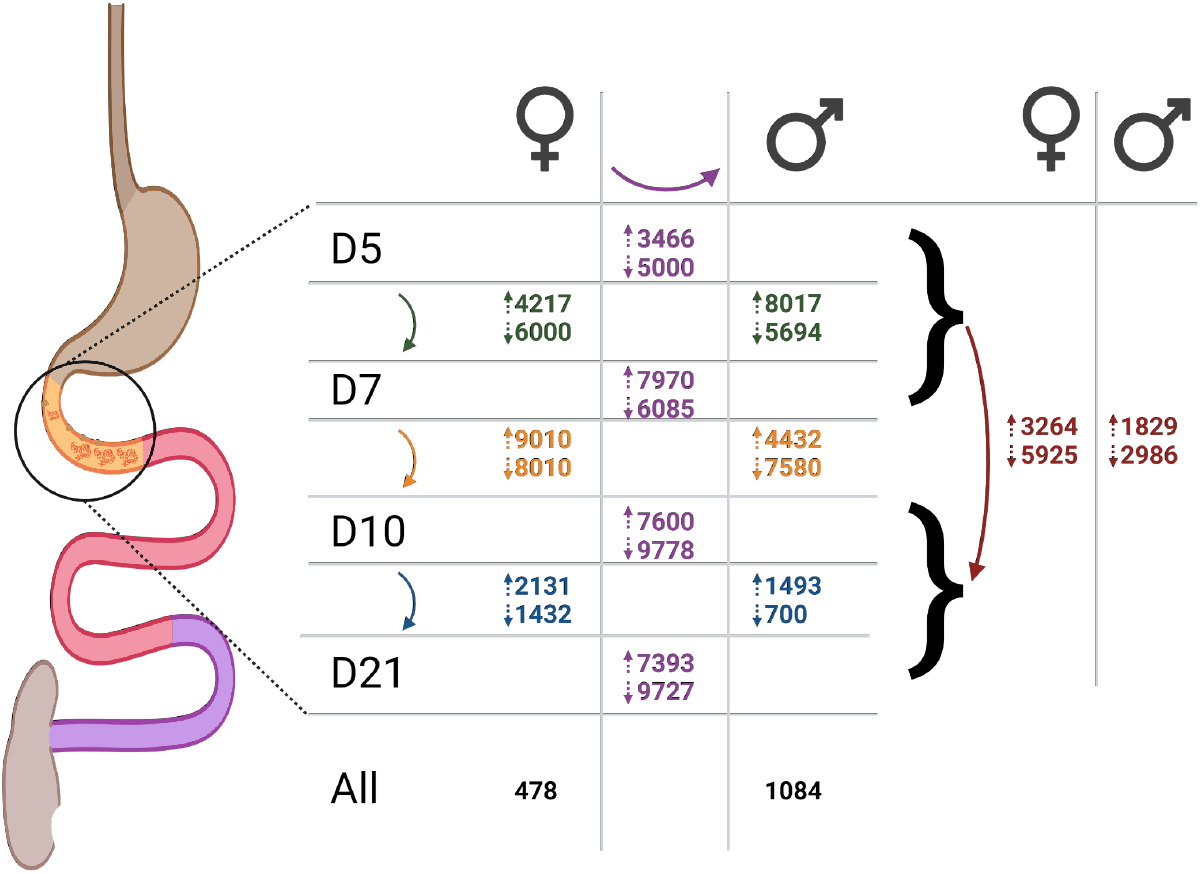
Numbers of transcripts statistically significantly up- and down-regulated in sample group comparisons. Numbers are positioned and coloured according to the comparison being described. Purple center: transcripts up-regulated (top number beside up arrow) or down-regulated (bottom number beside down arrow) in females vs. males at the age to the left, green: transcripts up- or down-regulated in D5 vs D7 females (left) or males (right), orange: transcripts up- or down-regulated in D7 vs D10 females or males, blue: transcripts up- or down-regulated in D10 vs D21 females or males, black: transcripts up-regulated in all female (URF) or male (URM) samples at all ages, red: tissue-dwelling vs lumen-dwelling comparisons (up: URTDF, URTDM; down: URLDF, URLDM).

### Transcription level differences between male and female worms

In addition to the obvious significance to reproductive biology and nematode transmission, sexual dimorphism has been found to be one of the biggest differences among lifecycle stages in a variety of nematodes [17–20]. *H. bakeri* does not have a sex-specific chromosome, like a Y chromosome as in many mammals or W chromosome in many birds. Rather, it has an XX/XO sex determination system like *C. elegans* [21,22] and therefore, there are no male-specific or female-specific genes in *H. bakeri*, only male-specific or female-specific expression of certain genes. To explore this sex-specific gene expression, we looked for transcripts that were consistently up-regulated in the males or females. These transcripts were statistically significantly differently transcribed (p_adj_ < 0.05 by DGE analysis, see Methods and Fig 3) between males and females of the same age that had higher expression in all the male samples (for up-regulated in males – URM) or in all the female samples (for up-regulated in females – URF) and were found in relevant modules of the co-expression networks (S3 Fig). Throughout all analyses, gene names used refer to *C. elegans* genes, with the corresponding *H. bakeri* locus tags from WormBase ParaSite (HPOL_XXXXXXXXXX) in S27 Table. Among the 1084 URM transcripts (S3 Table) were orthologs of genes in *C. elegans* with demonstrated association with males or demonstrated roles in male development (S4 Table). Notable examples include the transcriptionally regulated male development and patterning genes *mab-3* and *mab-23*; the male fate specification gene *her-1*; the male mating behaviour associated genes *eat-4, cil-7*, and *mapk-15*; and the spermatogenesis-related genes *spe-4, cpb-2, mib-1, fog-3*, and *cpb-1*. Additionally, among the URM transcripts were orthologs of genes in *C. elegans* with roles in the regulation of gene expression, whose expression in this *H. bakeri* context point to mechanisms by which male- or female-associated gene expression patterns may be established and/or maintained. These include genes involved in alternative splicing like *prmt-9* and *rsp-8*; glycosylation ZC250.2; and protein ubiquitination and protein folding *spop-1* and *cnx-1*. Finally, among the URM transcripts are orthologs of genes in *C. elegans* with roles in starvation, including *ser-6, gcy-35, pck-3, tre-2*, and *nemt-1*. Functional enrichment analysis of the entire URM set revealed 32 enriched gene ontology (GO) terms that predominantly describe protein modification processes and phosphorylation/dephosphorylation (S5 Table).

**Fig 3.**
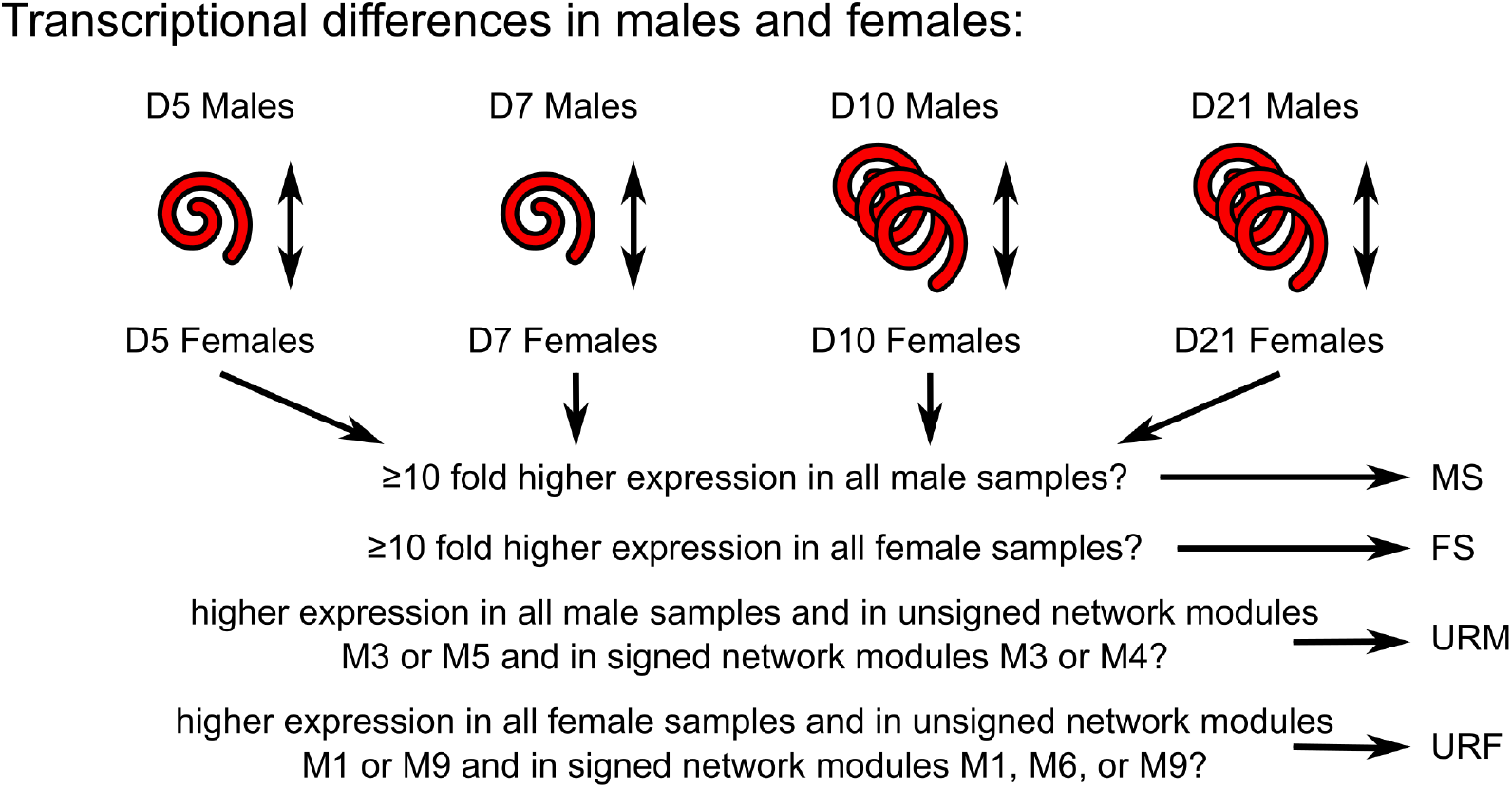
Flow diagram of male/female comparisons. The intersections of pairwise comparisons between age-matched male and female samples were filtered to yield the sets of transcripts discussed in the main text.

Among the 478 URF transcripts (S6 Table) were orthologs of genes in *C. elegans* with demonstrated roles in hermaphrodite development or maternal processes (S7 Table). These include genes involved in development of the vulva *soc-2* and *had-1*; development of the spermatheca *nhr-6*; the sex determination pathway *fox-1*; oogenesis *csn-1*; ovulation *ipp-5*; egg formation *perm-4* and *gna-2*; egg laying *bar-1, tmc-2*, and *sek-1*; and maternal roles in embryonic development *vha-7*. In contrast to the males, URF transcripts contain different orthologs of *C. elegans* genes involved in alternative splicing *ddx-15*; glycosylation *gale-1, ugt-64*, and ZK632.4; and protein ubiquitination *sli-1*, ZK430.7, and *cif-1*; as well as orthologs of genes involved in translational regulation of gene expression *eif-2Bα*, K07A12.4, *eif-3*.*E*, and *eif-3*.*D*. Moreover, rather than starvation-associated genes, URF transcripts include an ortholog of a *C. elegans* gene involved in eating *eat-20*. Finally, among the URF transcripts are orthologs of genes in *C. elegans* involved in stress responses including osmotic stress *nhr-1* and oxidative stress *aak-2, mek-1, mlk-1, trx-2*. Functional enrichment analysis of the entire URF set revealed 18 enriched GO terms that feature peroxisomes, mitochondria, and redox processes (S8 Table).

To further scrutinise the sex-specific gene expression, we defined male-specific transcripts as transcripts that were statistically significantly differently transcribed and that had ≥ 10-fold higher expression levels in all male samples compared to female samples of the same age (and vice versa for female-specific transcripts). These criteria (significantly different expression between males and hermaphrodites and ≥ 10-fold higher expression) when applied to *C. elegans* in a microarray study identified 285 male-specific and 160 hermaphrodite-specific genes [23]. Here in *H. bakeri*, among the 160 male-specific transcripts (S2 Table) are 103 transcripts with no annotation information, highlighting that many aspects of the male worms remain uncharacterized. Moreover, only two of these transcripts have an ortholog in the *C. elegans* genome: HPOL_0000701801 which is in a many-to-one relationship with W02D9.4 (an uncharacterized protein) and HPOL_0001987901 which is in a many-to-one relationship with F29B9.7 (an uncharacterized protein). The remaining 57 transcripts come from 50 genes that are shown in S26 Table. Functions of these genes, inferred from their annotation, include sperm production (HPOL_0001035601), male patterning (HPOL_0001902701), collagen synthesis and cross-linking (HPOL_0000750501, HPOL_0001117101, HPOL_0001117201, HPOL_0001117301, HPOL_0001117501), signalling cascades (HPOL_0000062301, HPOL_0000164101, HPOL_0000934501, HPOL_0000982501, HPOL_0001599701, HPOL_0001692501, HPOL_0001834101, HPOL_0001912801), and glycosylation (HPOL_0000317101). A high proportion of signalling cascade proteins has also been noted for spermatogenesis-enriched transcripts in a *C. elegans* microarray study [24]. Notably only one transcript fit the criteria as female-specific (HPOL_0000787001) and it is predicted to encode a peptidase (S26 Table).

Finding orthologs of known male-related genes in URM and female-related (hermaphrodite in *C. elegans*) genes in URF suggests that the filtering criteria used were appropriate to successfully recover transcripts important for the males and females, respectively. The URM and URF sets are, therefore, likely to reflect genuine differences between the males and females and not merely artifacts. The different sets of transcripts used between the males and females for alternative splicing, glycosylation, and protein ubiquitination point to the use of these processes to establish and/or maintain the important differences in gene expression between the males and females as well as potentially generate male/female isoforms of the targets of these gene products. It is surprising, however, to see oxidative and osmotic stress signatures among the URF transcripts. The collection of the worms from their host environment into medium is a stressful process that involves a transfer to a higher oxygen environment (See S1 Text for additional discussion on intestinal oxygen) as a source of oxidative stress. Additionally, it is possible the medium does not perfectly match the osmotic conditions of the mouse tissue and/or lumen, thus providing a source of osmotic stress. However, it is unclear why the females would be more responsive to these stresses than the males. Male worms have been found to die faster and in greater proportion when exposed to oxidative stressors like arsenite for *C. elegans* [25] and peroxide for *H. bakeri* [26]. Perhaps more sensitive and stronger stress responses in the female worms contributes to their increased tolerance of the stressors.

It is also curious that we find a starvation-like expression signature among the URM transcripts. It has been reported in *C. elegans* that the pharyngeal pumping rate decreases in males during mating from 180 pumps per minute to 50 [27]. However, since the D5 and D7 males are larvae that are still individually encysted in the intestinal tissue with no contact with a female they are not mating. Yet the starvation-like expression signature is present at these stages as well, which contradicts a mating-induced reduction in food intake in the males as an explanation for this transcript signature.

Additionally, starvation has been reported to inhibit mating behaviours in *C. elegans* males [28]. The infection conditions we used here to obtain the worms are also used to generate eggs to grow to infective L3 larvae to maintain the stock of worms in the laboratory, indicating that the adult worms do mate under these conditions. It is therefore unlikely that the male worms are truly starved of food, and using a bead-feeding assay, we found that males did feed (Fig 4 and S4 File). However, high feeding D10 adults were exclusively female (Fisher’s exact test, p=0.04), while the males only fed at low rates. This suggests it is likely that males ingest less food than the females (even after accounting for differences in body size) resulting in a net energy deficit during the parasitic phase of their life. Alternatively, the males may have a higher energy expenditure in general compared to the females, resulting in more pronounced liberation of energy stores and a higher drive to seek food, both of which are associated with starvation responses. In support of this, male *C. elegans* were found to have higher carbohydrate metabolism through the glycolytic pathway than hermaphrodites [23].

**Fig 4.**
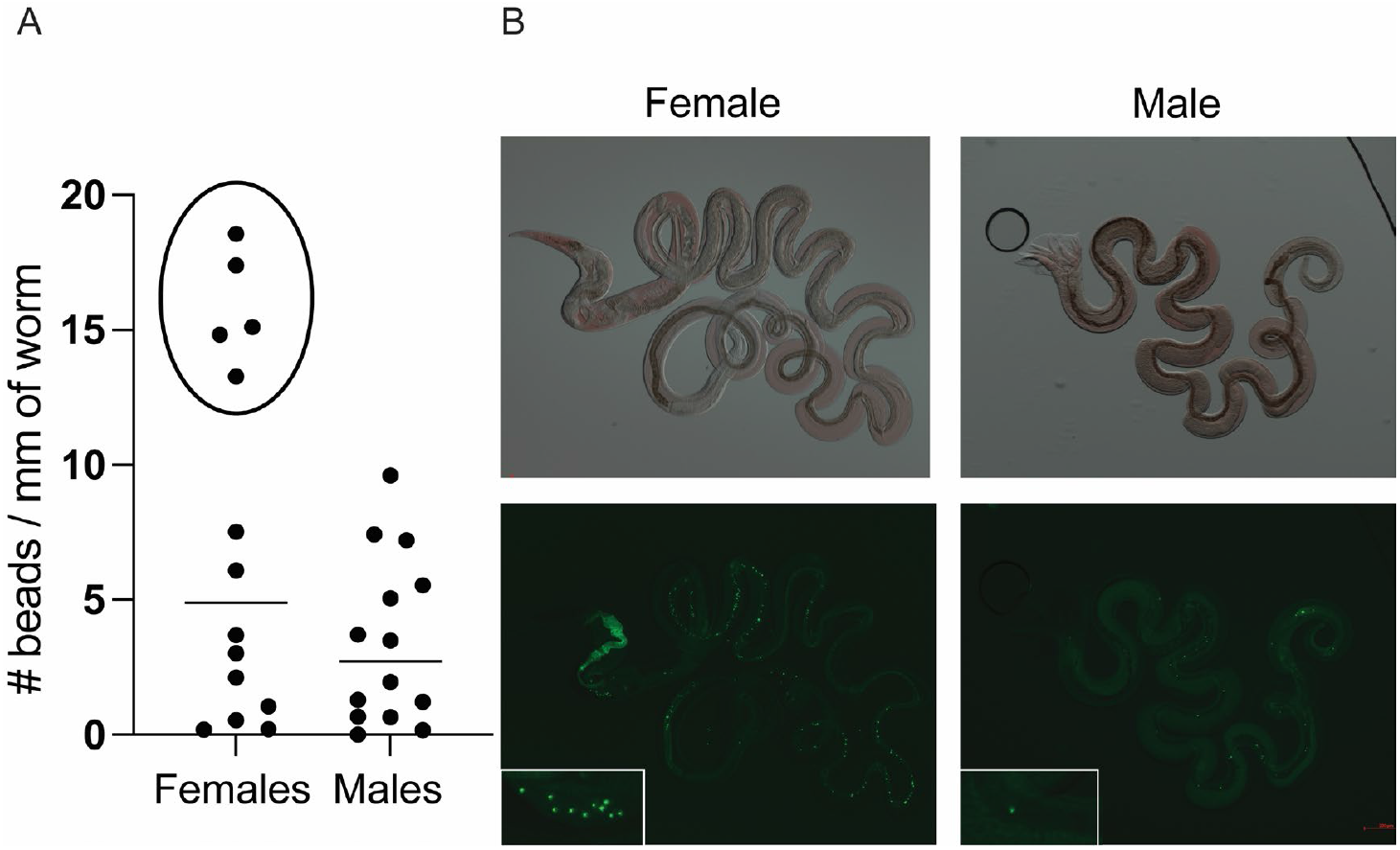
Comparison between the feeding of female and male D10 adults, both n=14. (A) The number of beads ingested normalised for the worms’ size difference. There was not a significant difference between the sexes (p=0.06), but the females and males were differently distributed. Only females ingested beads at the higher rate (circle). (B) Representative pictures of a female and male D10 worms. All the images are available in the S4 File.

The sheer number of transcripts that are significantly differentially expressed between males and females, both in URM + URF but also D10 (68.92%) and D21 (67.90%) (Fig 2 and Fig 5), reflects a significant level of sexual dimorphism at the transcriptional level in this nematode. We additionally analysed the RNA-seq datasets from *Haemonchus contortus* where the adult males and adult females were sequenced separately [20] and found 12669 of 21477 transcripts (58.99%) to be significantly differentially expressed between the adult males and females (using our same criteria as for *H. bakeri*). In *C. elegans*, a microarray study of adult males and hermaphrodites found 14488 / 26843 (53.97%) transcripts to be differentially expressed [23]. In all three nematodes, these proportions of transcripts that are differentially expressed between the adult sexes point to significant transcriptional sexual dimorphism in general. Our dataset additionally allows examination at the fourth larval stage, where sexual dimorphism is less prominent than in adults but is still notable (D5 – 33.58% and D7 – 55.74% transcripts differentially expressed between males and females; Fig 2). A microarray study from *C. elegans* L2, L3, and L4 worms that compared hermaphrodites to masculinized hermaphrodites (*tra-2(ar221ts)*; *xol-1(y9)* worms are strongly masculinized, fertile XX pseudomales) also found sexual dimorphism among the larval stages [29]. Though they identified fewer genes as sex-regulated than we have (possibly because they focused on somatic tissues while excluding germline tissues, and/or because of the lower sensitivity of microarrays compared to RNA-seq, and/or because of differences between the two species), they also found fewer genes to be sex-regulated among the larvae compared to adults, as well as many more male-enriched genes than hermaphrodite-enriched genes: both trends that we see here in *H. bakeri*. The greater number of male-enriched genes was postulated to be a consequence of the suppressive nature of the TRA-1 master sex-regulator in *C. elegans*, a gene that *H. bakeri* has a one-to-one ortholog of (HPOL_0000251701). These commonalities between the two worms support the view that TRA-1 suppressing male developmental programs in XX worms is a common feature among nematodes, whether they are dioecious like *H. bakeri* or androdioecious like *C. elegans*. Moreover, the same study found little overlap between the differentially expressed genes detected at early vs late larval stages [29]. This suggests significant, unexplored, sexual dimorphism exists even in the morphologically indistinct larval stages of other nematodes. Given the transcriptional sexual dimorphism in these nematodes, male/female differences should be taken into consideration in all future experiments, especially in dioecious species like *H. bakeri* or *H. contortus* where males and females exist in roughly uniform proportion.

**Fig 5.**
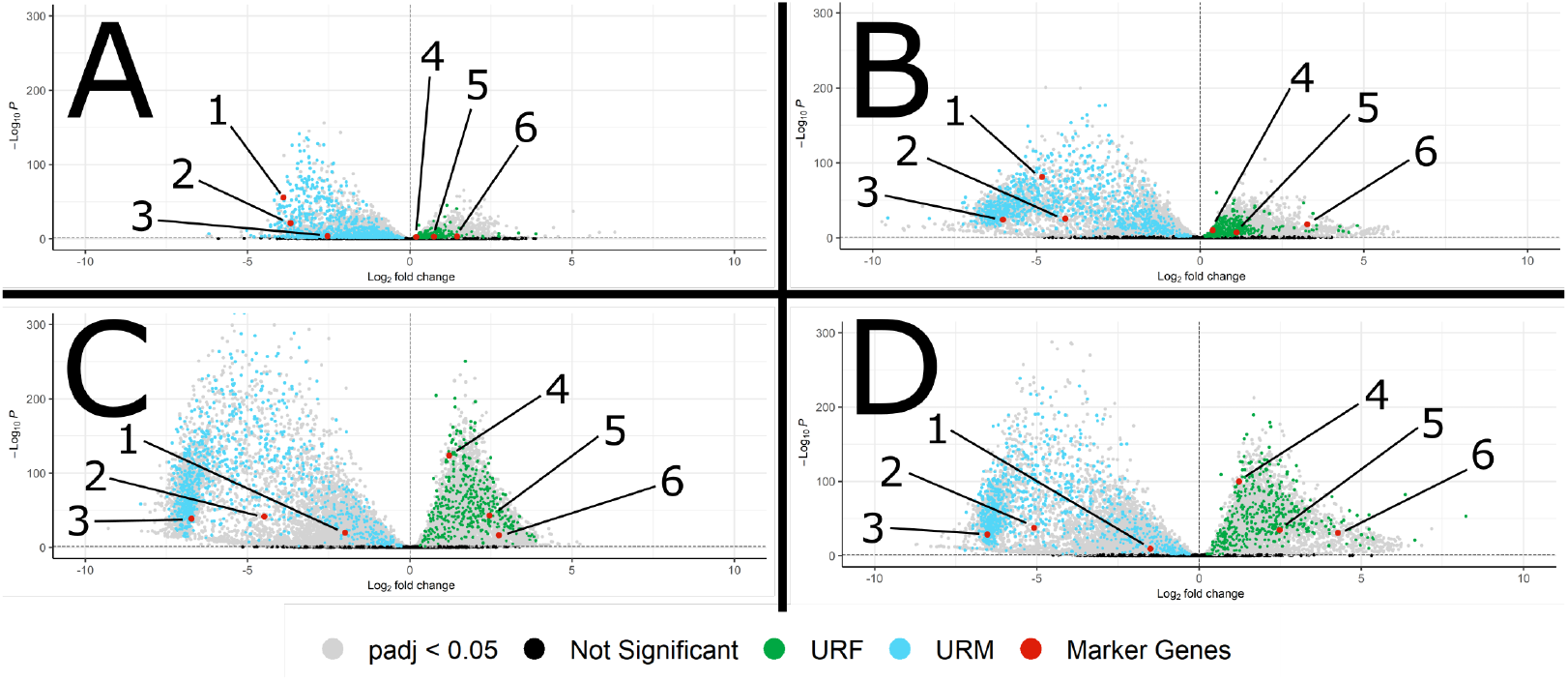
Volcano plots of male / female comparisons at D5 (A), D7 (B), D10 (C), and D21 (D). The transcripts in URM (blue) or URF (green) are shown. Negative log2FoldChanges denote higher expression in the male group while positive values indicate higher expression in the female group. Marker genes in red are 1) *her-1* (HPOL_0000740701), 2) *mab-3* (HPOL_0001902701), 3) *spe-4* (HPOL_0000308001), 4) *csn-1* (HPOL_0001830501), 5) *fox-1* (HPOL_0001264601), and 6) *perm-4* (HPOL_0000100601).

### Developmental transcriptional changes

The timepoints throughout infection for the samples used here include when *H. bakeri* larvae are encysted in the intestinal tissue (D5 and D7) and when the adults have emerged into the lumen (D10 and D21). To identify the transcriptional changes that occur throughout this final phase of development we defined, for both the males and the females, a set of transcripts up-regulated in the adults (lumen-dwelling) and a set of transcripts up-regulated in the larvae (tissue-dwelling). To be considered up-regulated in tissue-dwelling males (URTDM), transcripts had to: i) have higher expression in all D5 and D7 male samples than in all D10 and D21 male samples, ii) be in relevant modules of the co-expression networks (S3 Fig), and iii) have statistically significantly different expression in either of the tissue-dwelling vs lumen-dwelling pairwise comparisons being considered (See Methods and Fig 6). To be considered up-regulated in the lumen-dwelling males (URLDM), transcripts had to: i) have higher expression in all D10 and D21 male samples than in all D5 and D7 male samples, ii) be in relevant modules of the co-expression networks (S3 Fig), and iii) have statistically significantly different expression in either of the tissue-dwelling vs lumen-dwelling pairwise comparisons being considered. The same filtering criteria were applied to the female samples to find the transcripts up-regulated in the tissue-dwelling females (URTDF) and up-regulated in the lumen-dwelling females (URLDF).

**Fig 6.**
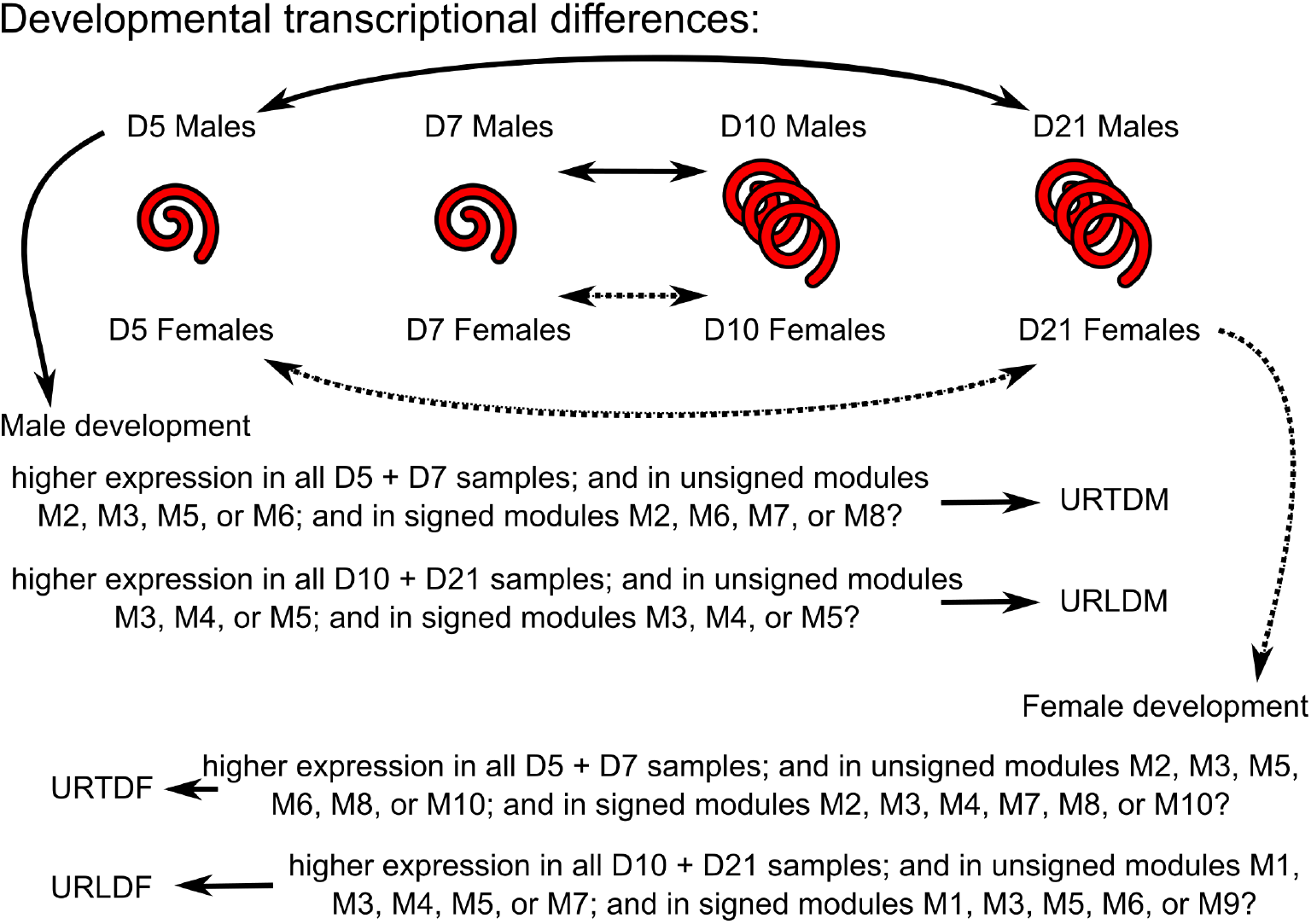
Flow diagram of developmental comparisons. Transcripts with expression patterns of interest were filtered according to co-expression network module and statistically significant differences in expression in pairwise comparisons between tissue-dwelling and lumen-dwelling worms to yield the sets of transcripts discussed in the main text.

Among the 1829 URTDM transcripts (S9 Table) were orthologs of genes in *C. elegans* with demonstrated roles in muscle development *lev-11*; epithelial development *ltd-1* and *efn-2*; cuticle synthesis and molting *dpy-31, phy-2, tsp-15, fkb-3*, and numerous cuticlins and collagens; and male tail development *lon-8* and *mab-7* (S10 Table). Additionally, among URTDM transcripts were orthologs of genes in *C. elegans* with roles in environmental sensing *nep-2* and signal transduction *rrc-1* and *pde-6*. Functional enrichment analysis of the entire URTDM set revealed 33 enriched GO terms that featured the plasma membrane and cuticle synthesis, including structural constituents and procollagen-proline dioxygenase activities (S11 Table). Within the 3264 URTDF transcripts (S12 Table) were orthologs of genes in *C. elegans* with known roles in muscle development *unc-52* and *stn-1*; epithelial development *efn-2*; nervous system development *irx-1, grdn-1, mig-13, kal-1, mnr-1*, and *mig-1*; cuticle synthesis and molting *bus-19, phy-2, mlt-7, zmp-2*, and numerous cuticlins and collagens; and regulation of body size and length *mua-3, sma-6*, and *lon-8* (S13 Table). Additionally, among URTDF transcripts were orthologs of genes in *C. elegans* with roles in aerobic respiration and oxygen sensing *ucr-2*.*3, isp-1, pdl-1*; environmental sensing *nep-2*; and nutrient absorption *sms-5*. Functional enrichment analysis of the entire URTDF set revealed 67 enriched GO terms that featured oxygen binding and transport, cuticle synthesis, and nervous system development (S14 Table). The URTDM and URTDF sets share 1560 transcripts (S15 Table). Functional enrichment of the shared up-regulated in tissue-dwelling worms transcripts revealed 31 enriched GO terms that feature adhesion, cuticle and molting cycle, and oxygen binding and transport (S16 Table).

Among the 2986 URLDM transcripts (S17 Table) was an ortholog of the *C. elegans* spermatogenesis-related gene, *ubxn-2*. Additionally, among the URLDM transcripts were orthologs of genes in *C. elegans* with roles in the cuticle *col-36*; and metabolism *sucg-1* and *tre-1* (S18 Table). Functional enrichment analysis of the entire URLDM set revealed 64 enriched GO terms that feature phosphorylation/dephosphorylation, ubiquitin ligase activity, and protein modification processes (S19 Table). Among the 5925 URLDF transcripts (S20 Table) were orthologs of genes in *C. elegans* with known roles in adult gonad maintenance and development *ippk-1, mys-1, tin-9*.*2*, and *evl-20*; egg laying *fahd-1* and *rfp-1*; maternal factors for embryonic development *mel-32, par-4, dnc-4*, and *emb-4*; and germline maintenance and gametogenesis *clk-2, dvc-1, stau-1*, and *etr-1* (S21 Table). Additionally, among URLDF transcripts were orthologs of genes in *C. elegans* with roles in mRNA splicing *prcc-1* and *prp-19*; translation *eif-3*.*C*; protein deubiquitination *otub-1*; heparan sulfate metabolism *hst-2* and *pst-2*; and negative regulation of aerobic respiration *blos-1* or response to reoxygenation *stl-1*. Functional enrichment analysis of the entire URLDF set revealed 262 enriched GO terms that feature regulation of gene expression (RNA processing, glycosylation, ubiquitination, histone modification, and translation terms) and cell cycle terms (S22 Table). The URLDM and URLDF sets share 1007 transcripts (S23 Table). Functional enrichment of the shared up-regulated in lumen-dwelling worms transcripts revealed 59 enriched GO terms that feature cell cycle and ubiquitin protein catabolic processes (S24 Table).

Since the tissue-dwelling worms are still developing, finding in URTDM and URTDF orthologs of genes in *C. elegans* with known roles in development suggests that the filtering criteria used were appropriate to successfully recover transcripts important for the tissue-dwelling worms. Likewise, finding in URLDM and URLDF orthologs of genes in *C. elegans* with known roles in gametogenesis and reproduction suggests that the filtering criteria used were appropriate to successfully recover transcripts important for the lumen-dwelling worms. It is, therefore, worth additional investigation into the importance heparan sulfate metabolism may be playing in the adult female worms and into whether the additional phosphorylation and dephosphorylation activities in the adult males are purely a consequence of spermatogenesis or whether this also reflects other critical male processes.

It has been reported previously that nematodes are capable of anaerobic respiration (ex. *Ascaris suum* [30], *Haemonchus contortus* [31], *C. elegans* [32]). Additionally, a study comparing three free-living nematodes (*C. elegans, Pristionchus pacificus*, and *Panagrolaimus superbus*) and one plant-parasitic nematode (*Bursaphelenchus xylophilus*) found the parasitic nematode survived anaerobic conditions much longer than the free-living ones [33], suggesting surviving low oxygen environments may be a particularly important adaptation for parasitic nematodes compared to their free-living relatives. The enriched GO terms involving oxygen binding, transport, and utilization among URTDM and URTDF transcripts (and the lack of such terms among the URLDM and URLDF transcripts) suggests that an increased role for anaerobic respiration, at least in *H. bakeri*, occurs during the transition from the L4 to the adult stage. This transition is when the worms migrate from the intestinal mucosa, a physiologically aerobic environment, to the intestinal lumen, a physiologically hypoxic environment [34] (See S1 Text for additional discussion on intestinal oxygen). The transition from the L4 to adult stage is also when the final molt occurs, along with synthesis of the final cuticle. This aligns with the abundance of collagens, cuticlins, and cuticle-related GO terms that are enriched in the URTDM and URTDF transcripts. Among the many steps involved in cuticle synthesis is the modification of certain proline residues in the collagen molecules to 4-hydroxyproline [35], a process requiring molecular oxygen [35,36], and the process most likely reflected in the enriched GO terms involving procollagen-proline dioxygenase activity (S11, S14, S16 Tables). Characterized human enzymes that perform this function (prolyl 4-hydroxylases) catalyze the reaction L-proline + 2-oxoglutarate + O_2_ → 4-hydroxyproline + succinate + CO_2_ [36]. In *H. bakeri*, this occurs when the L4 worms are in a physiologically aerobic environment and all other molts and synthesized cuticles in *H. bakeri* occur outside the host in a fully aerobic environment. We hypothesize that the levels of oxygen in the intestinal lumen are insufficient to support this collagen modification (and/or *H. bakeri* is unable to scavenge the required oxygen), which suggests that cuticle synthesis is a strictly aerobic process. In support of this, prolyl 4-hydroxylases are also involved in oxygen sensing through their oxygen-dependent modification and destruction of the hypoxia inducible factor [36], a transcription factor that is induced in the host epithelial cells lining the intestinal tract [34]. If luminal oxygen is indeed insufficient to support prolyl 4-hydroxylase activity, the need for molecular oxygen to get through the final molt within the host could be a driver of the worms encysting in the intestinal mucosa, a phenomenon that fully exposes the worms to the host immune system (in contrast to the lumen which is beyond the reach of many immune effectors). Many parasitic nematodes encyst within host tissue (ex. *Cooperia punctata* [37] and *Trichinella spiralis* [38], which both invade the intestinal mucosa during their development), migrate through aerobic host tissues (ex. *Ascaris lumbricoides* [39], *Necator americanus* [40], and *Nippostrongylus brasiliensis* [41], which all migrate through circulatory and pulmonary tissues), or actively seek host blood (ex. *Haemonchus contortus* [42] and *Ancylostoma duodenale* [40]), which would all provide a rich source of molecular oxygen for synthesis of the final cuticle regardless of where the adults ultimately reside in the host.

Since our timepoints span much of the adult life of *H. bakeri*, we investigated potential gene expression signatures of aging. The expression in *H. bakeri* of orthologs of genes in *C. elegans* that have been implicated in aging shows that our *H. bakeri* dataset has insufficient samples across the adult stage to robustly identify genes implicated in aging based on their expression patterns alone (S4 Fig and S1 Text for additional discussion on aging-related gene expression). However, the *H. bakeri* orthologs of the *C. elegans* IIS pathway [43] show marked differences in expression pattern between the males and the females (Fig 7), suggesting that molecular pathways involved in aging may differ between males and females. It has been reported previously in *C. elegans* that transcription of certain isoforms (d/f) of the *daf-16/FOXO* transcription factor is affected in a sex-specific manner by TRA-1 to increase *daf-16* activity [44]. Our results, however, show a notable decrease in the transcription of *daf-16* (HPOL_0000379201) in *H. bakeri* females upon reaching adulthood, in contrast to steadily increasing transcription in the males as their parasitic life progresses (Fig 7). It is worth noting, however, that in the current *H. bakeri* genome annotation there is only one isoform predicted for *daf-16* (HPOL_0000379201), which corresponds to the b/c isoforms in *C. elegans* (data not shown), highlighting that the annotations in *H. bakeri* still need work.

**Fig 7.**
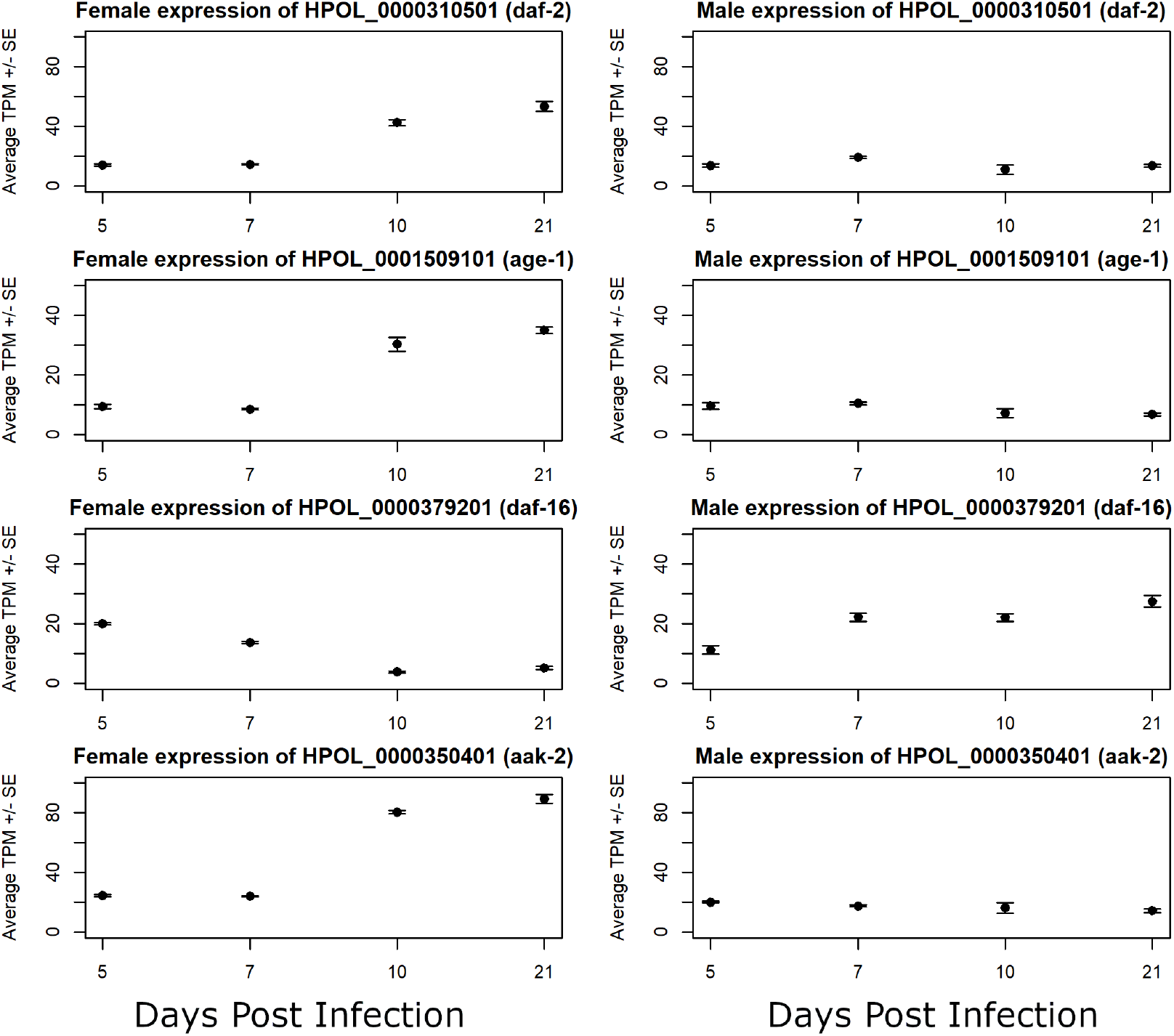
Expression in *H. bakeri* of orthologs of genes in *C. elegans* implicated in aging. Average TPM (transcript per million) are plotted for females (left) and males (right) at each timepoint. Error bars represent standard error. The insulin receptor (*daf-2*), kinase (*age-1*), and *daf-16/FOXO* transcription factor (*daf-16*) of the IIS pathway are shown along with the master energy regulator (*aak-2*).

### Expression of immunomodulatory genes and anthelmintic target genes

The majority of the male/female and developmental transcriptional responses described here center around the intestine, hypodermis, and gonad tissues of the worms. An estimate of tissue sizes in *C. elegans* places these three tissues as the largest in the worm, cumulatively accounting for more than three quarters of the worm’s tissue volume [45]. Since bulk RNA-seq pools together RNA from the whole worm, it is not surprising that the signatures of these tissues dominate our results. However, one phenotype of particular interest in *H. bakeri*, immunomodulation of its host, is not necessarily governed by these tissues. While some of the excreted/secreted products of *H. bakeri* with immunomodulatory activity were found to originate in the intestine (vesicles containing microRNAs) [15], the excreted/secreted proteins with described or implied immunomodulatory activity have no demonstrated tissue of origin in the worm. Excreted/secreted proteins in other parasitic worms have been found to originate from the uterine fluid and/or other sources, like the secretory apparatus [46]. Therefore, a possible source tissue for *H. bakeri* immunomodulatory proteins are excretory cells, which make up a very small proportion of whole *C. elegans* [45]. Comparative analyses may be able to uncover differential signal originating from rarer cell types like the excretory cells, especially if the differences between conditions are extreme. We therefore investigated the expression of loci that have been associated with immunomodulatory activity to see what comparisons or expression patterns might identify other immunomodulatory candidates. Most of the excretory/secretory proteins in *H. bakeri* that have been associated with immunomodulatory activity have not been identified but are members of certain classes of genes with common annotation descriptions (S25 Table). We therefore included all loci that matched each description. Expression patterns within these groups of genes vary considerably (S5 Fig). Moreover, even among described immunomodulatory proteins there are opposing expression patterns that exclude any one pairwise comparison from being more likely to identify immunomodulatory candidates (Table 2, Fig 8, and S1 Text for additional discussion on immunomodulatory gene expression). Hb-TGM-6 was identified as HPOL_0001864701 [47], which has higher expression in lumen-dwelling males than in tissue-dwelling males (as found with mixed-sex protein secretion) as well as sex-linked differences in expression (Fig 8 and S5 Fig). HbARI was identified as HPBE_0000813301 in the PRJEB1203 annotation [48], which when aligned against PRJEB15396 corresponds to HPOL_0001636401 (Fig 9), which has higher expression in the tissue-dwelling worms than in the lumen-dwelling worms (Fig 8 and S5 Fig). These data demonstrate that the expression of these two families of immunomodulators is tightly controlled. Moreover, our findings are consistent with the parasite’s need for these immunomodulators during infection. HbARI interferes with the release of IL-33, an alarmin which initiates the immune response to nematode infection and whose release is triggered by tissue damage [48]. IL-33 was found to be produced early in murine infection with the nematode *Trichuris muris*, which promotes worm expulsion in resistant mice [49]. Production and release of HbARI early in *H. bakeri* infection, when the worms are in the tissue and causing damage, helps regulate the host response to the worm’s damage and allows the worms to persist long enough to make it into the lumen where they are not causing the same level of tissue damage. In contrast, Hb-TGM is needed later in infection, where it stimulates TGF-β signalling and interferes with regulatory immune cells [47].

**Table 2.** Expression of immunomodulatory genes in *H. bakeri*.

| Protein Name | HbARI | HbBARI | HbBARI_Hom2 | Hb-TGM-6 |
| --- | --- | --- | --- | --- |

| *H. bakeri*<br>Locus Tag | HPOL_0001636401 | HPOL_0001228301 | HPOL_0001228401 | HPOL_0001864701 |
| --- | --- | --- | --- | --- |
| D5 Male<br>Average TPM | 1157.52 | 85.66 | 2189.10 | 11.08 |
| D5 Male vs D7<br>Male padj | 2.78E-01 | 1.81E-01 | 2.79E-25 | 1.14E-06 |
| D7 Male<br>Average TPM | 1137.80 | 71.52 | 599.96 | 47.05 |
| D7 Male vs<br>D10 Male padj | 3.91E-43 | 6.40E-01 | 2.18E-212 | 1.26E-02 |
| D10 Male<br>Average TPM | 145.43 | 53.57 | 5.80 | 63.04 |
| D10 Male vs<br>D21 Male padj | 1.48E-01 | 2.35E-07 | 9.88E-39 | 6.78E-01 |
| D21 Male<br>Average TPM | 201.43 | 217.55 | 0.65 | 87.04 |
| D5 Female<br>Average TPM | 1052.13 | 228.89 | 2097.85 | 16.25 |
| D5 Female vs<br>D7 Female<br>padj | 2.91E-01 | 9.60E-01 | 2.56E-08 | 5.92E-03 |
| D7 Female<br>Average TPM | 1070.11 | 202.22 | 824.01 | 30.71 |
| D7 Female vs<br>D10 Female<br>padj | 1.65E-174 | 2.62E-07 | 0.00E+00 | 1.85E-03 |
| D10 Female<br>Average TPM | 64.18 | 85.79 | 7.71 | 17.48 |
| D10 Female vs<br>D21 Female<br>padj | 7.07E-01 | 1.07E-01 | 4.38E-41 | 3.62E-01 |
| D21 Female<br>Average TPM | 63.68 | 140.27 | 0.06 | 13.54 |

**Fig 8.**
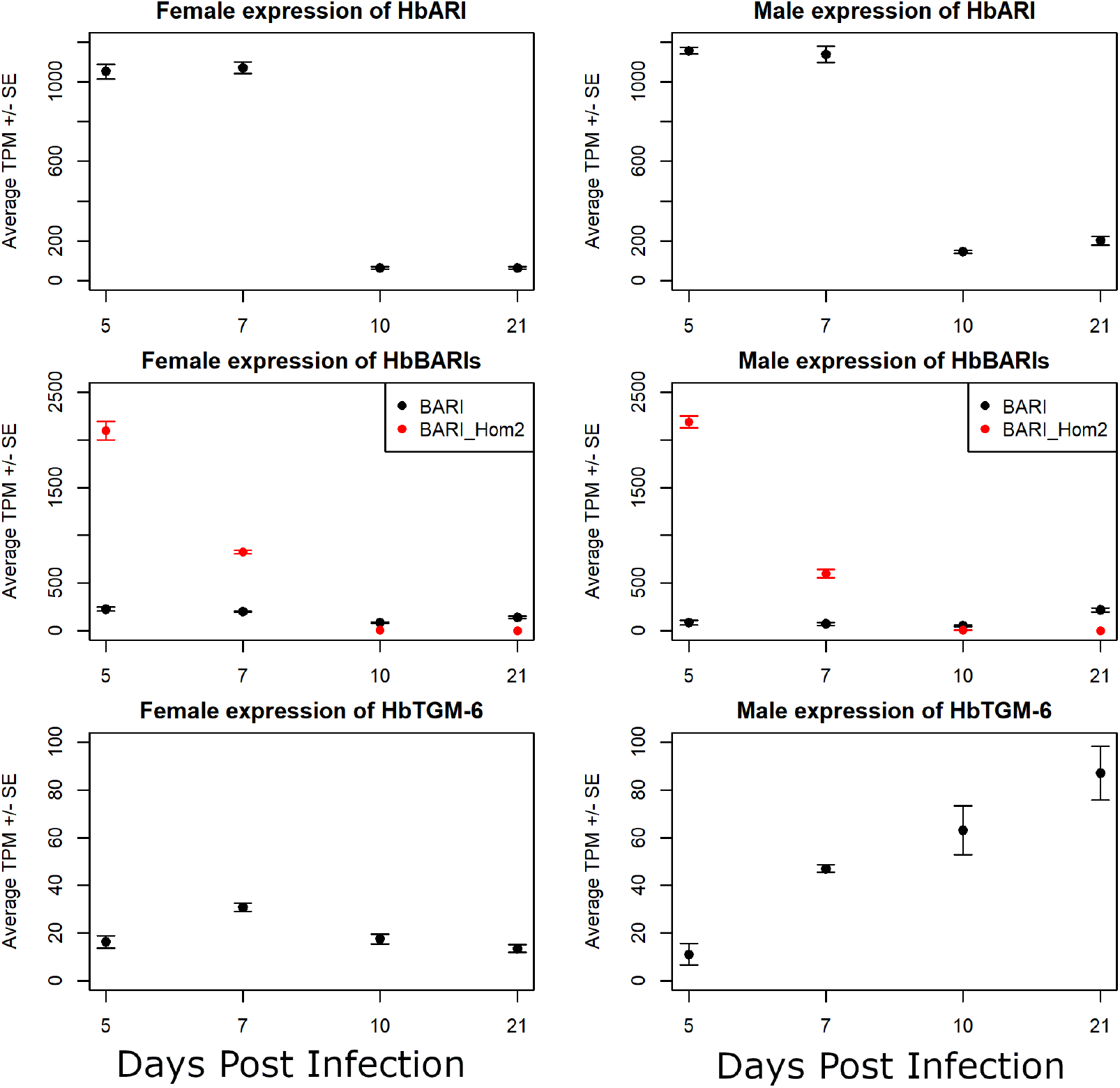
Expression of immunomodulatory genes in male and female *H. bakeri*. Average TPM (transcript per million) are plotted for females (left) and males (right) at each timepoint. Error bars represent standard error. The only identified immunomodulatory genes to date are plotted: HbARI (top row), HbBARI and HbBARI_Hom2(middle row), and HbTGM-6 (bottom row).

**Fig 9.**
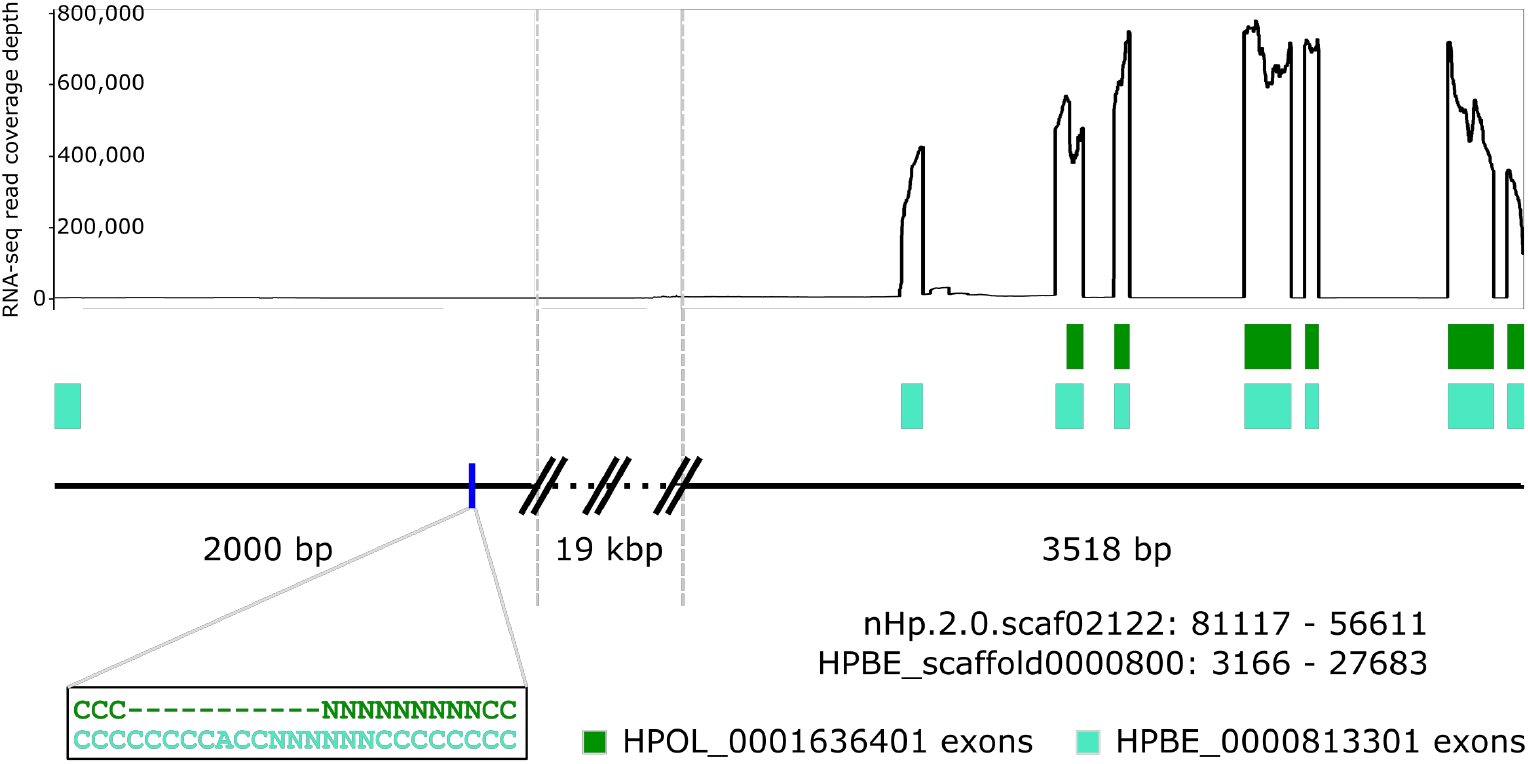
Schematic of the HbARI locus highlighting that the annotations in *H. bakeri* still need work. The two genome assemblies for *H. bakeri*, PRJEB1203 and PRJEB15396 (shown collectively as a black line), agree in this region with the exception of the area shown in the black box. The conflicting annotations for HbARI, HPBE_0000813301 (teal) and HPOL_0001636401 (green), are shown in their annotated positions and drawn to scale. HPOL_0001636401 is shorter at the 5’ end and consequently does not contain the signal peptide predicted to be in HPBE_0000813301. Since we know HbARI is a secreted protein because it was identified among HES, the PRJEB15396 annotation must be incorrect. As it is, HPOL_0001636401 is predicted to localize to the mitochondrion (data not shown). The coverage of all of the RNA-seq reads generated in this study in this region is shown as a line graph above the annotations. The HPBE_0000813301 annotation contains an extra 5’ exon that has no support in any of the RNA-seq datasets.

It is likely that *H. bakeri* secretes other proteins with immunomodulatory activity. However, these are unlikely to have uniform expression throughout the worm, may very well be secreted by a handful of specialized cells, and are not specific to males, females, or a lifecycle stage sampled here. Therefore, a higher resolution technique, like single-cell RNA-seq, would be needed to identify other immunomodulatory candidates based on their transcriptional expression patterns.

We also analyzed the expression profiles of known anthelmintic drug targets (Table 3 and Table 4). Of note, the majority of the drug targets have medium or low levels of transcription in both sexes at all stages sampled (TPMs range from 0 to 34856.8, with the top 10% of transcript expression corresponding to a TPM ≥ 66.9, top 30% corresponding to a TPM ≥ 14.7, and top 50% corresponding to a TPM ≥ 2.8). It is unclear if the medium and low expression reflect medium/low expression throughout the whole worm or high expression in a few cells and very low expression in the major tissues of the worms. The three highly expressed transcripts (top 10%) are the targets of benzimidazoles and a neuropeptide GPCR. Expression of the majority of the drug targets varies significantly between at least two of the sampled timepoints in both males and females, which may indicate higher drug efficacy on worms of particular ages or stages of development.

**Table 3.**
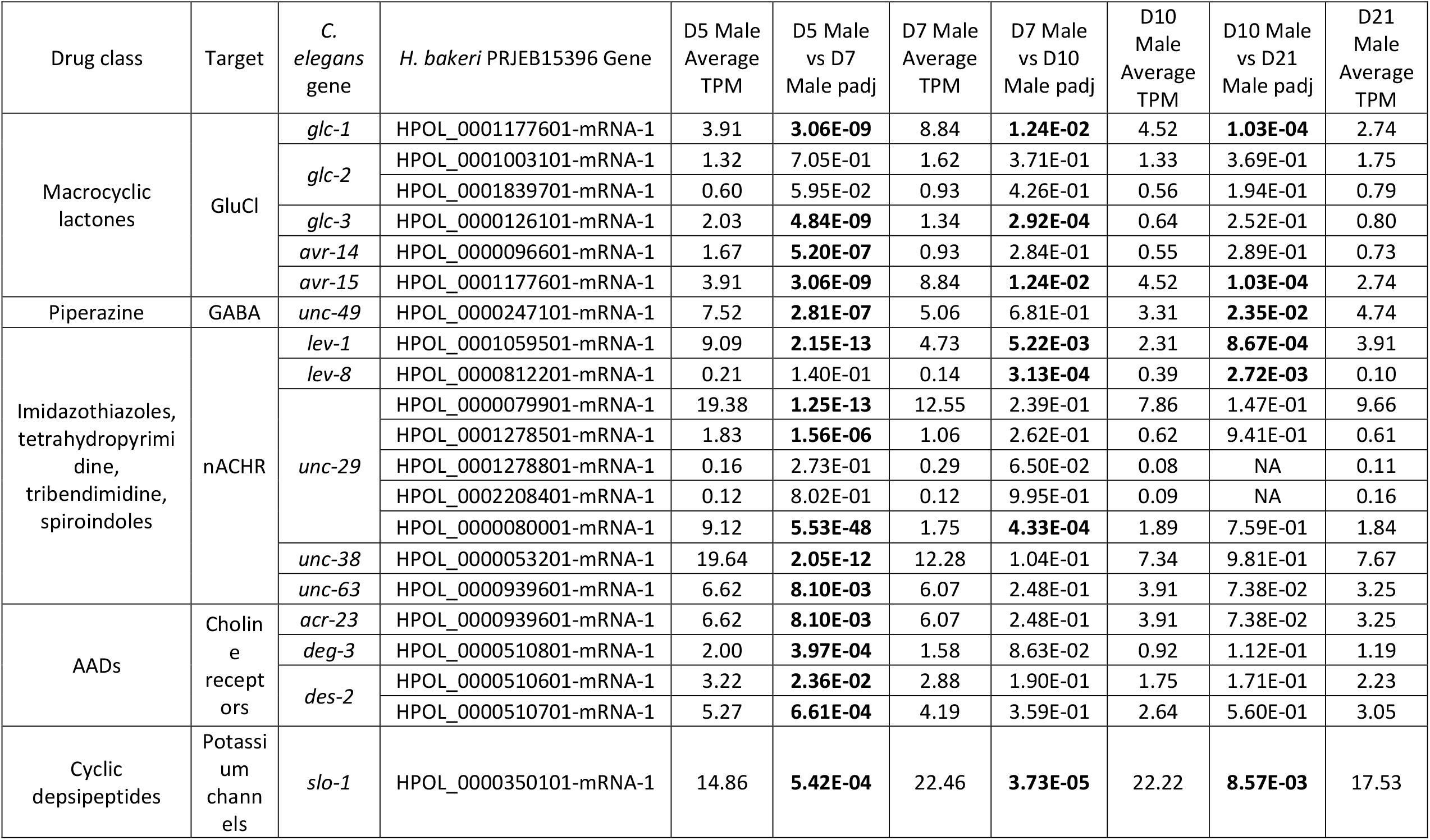
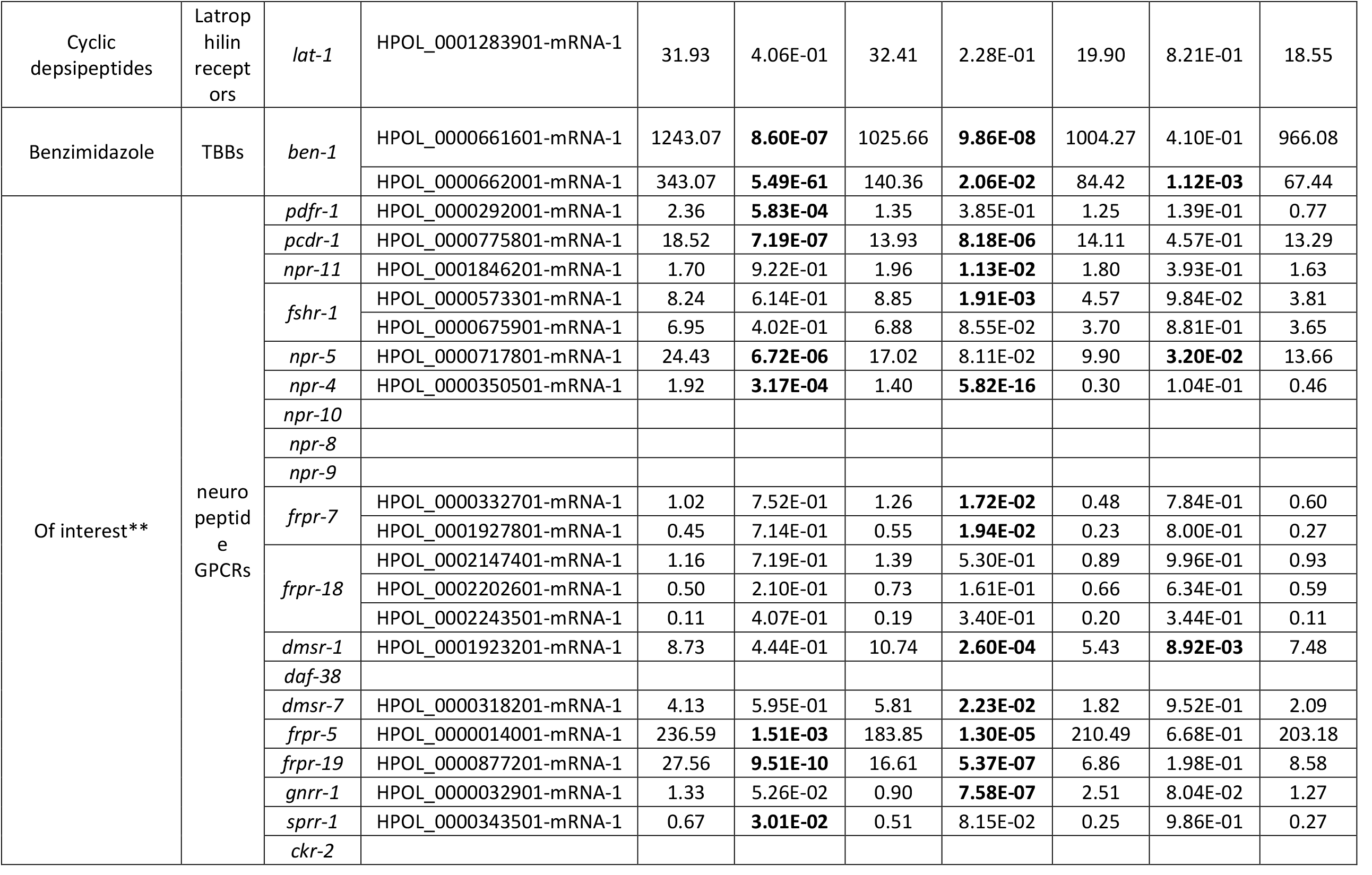
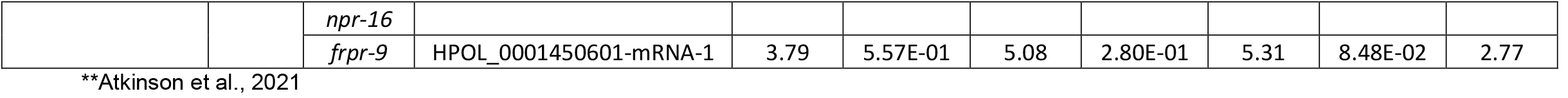
Expression of drug targets in male *H. bakeri*. Values in bold are padj < 0.05.

**Table 4.**
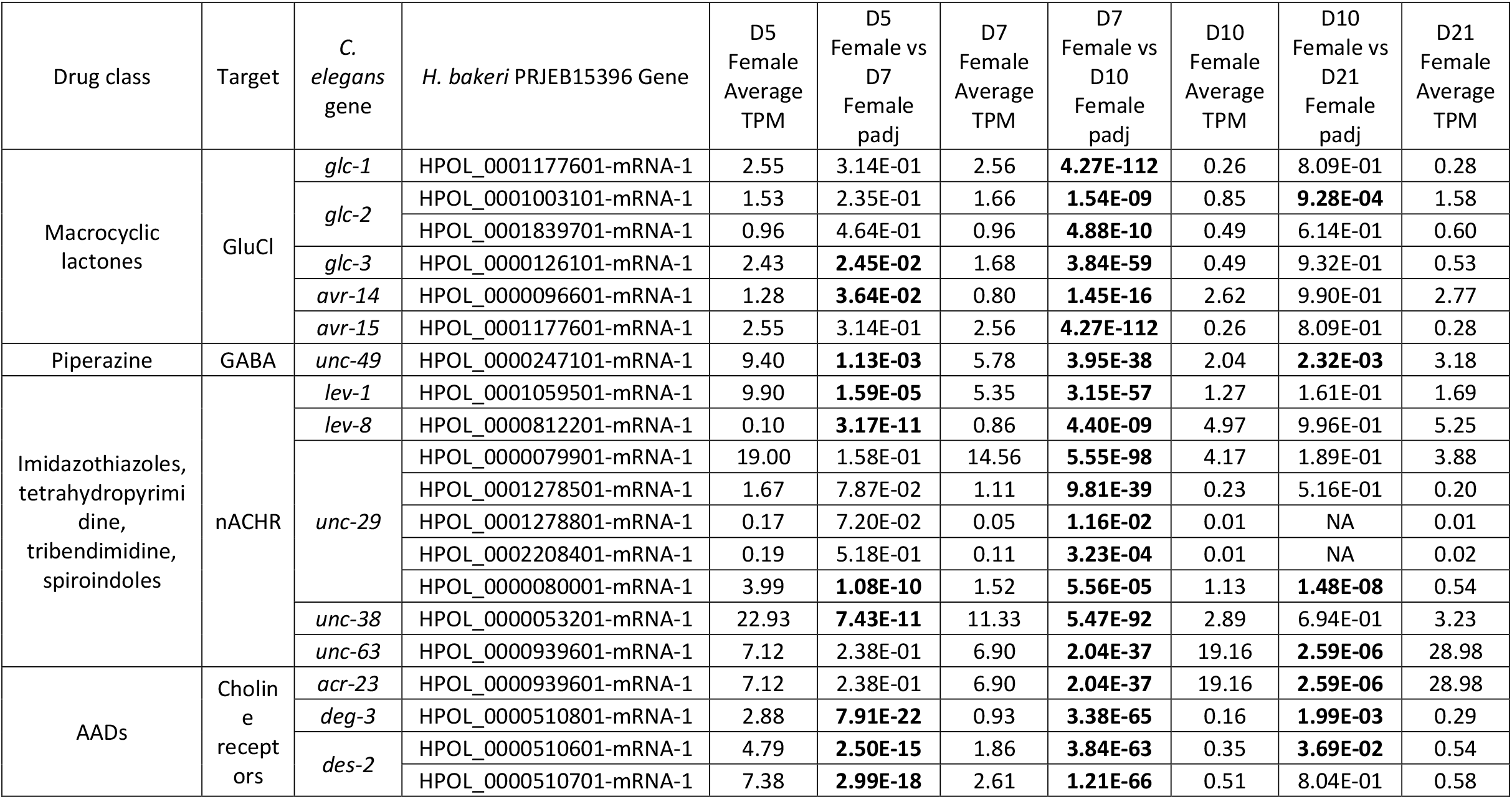
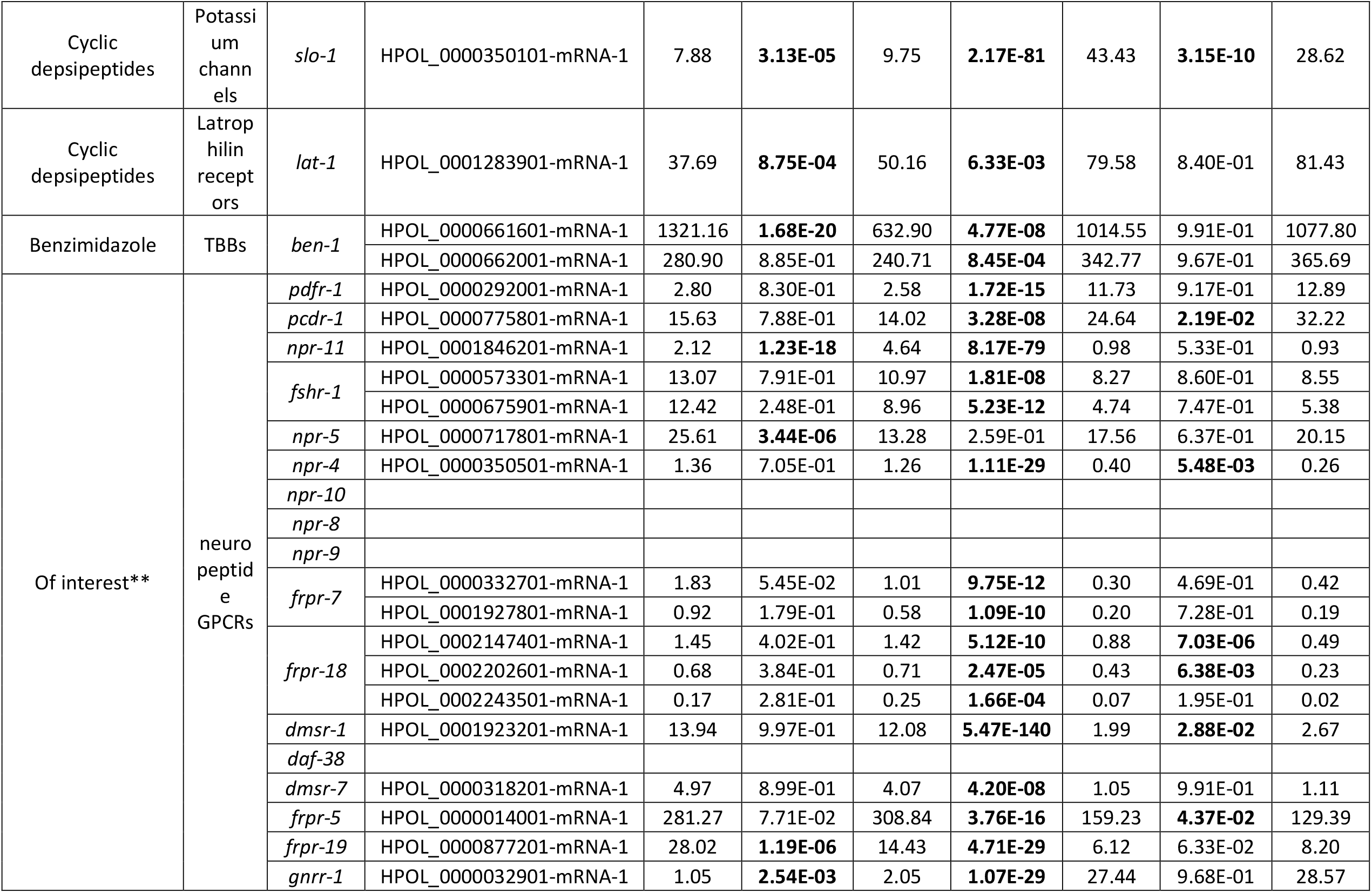
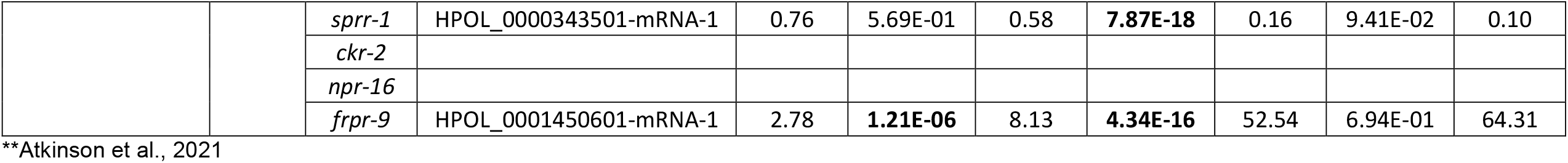
Expression of drug targets in female *H. bakeri*. Values in bold are padj < 0.05.

## Conclusions

By separating the males and females at each timepoint in our biologically replicated mRNA transcriptomes that span the parasitic phase of the *H. bakeri* lifecycle, we have been able to examine sexual dimorphism in this species and examine development without masking important sex-specific signals. We have uncovered inconsistencies in the available genome annotations for *H. bakeri*, along with other limitations that demonstrate a need to continue the work of annotating available genome sequences for this organism. We have identified processes that are key in establishing and/or maintaining sex-specific gene expression in this worm, without having sex-specific genes, including alternative splicing, glycosylation, and ubiquitination. Additionally, we find stronger oxidative and osmotic stress responses among female worms, potentially accounting for their previously reported better survival of oxidative assault. We hypothesize that oxygen concentration may be an important factor in the encysting behaviour of larval stage *H. bakeri* (in order to get through their last molt), a factor that could be more general among parasitic nematodes. We demonstrate the advantage of quantifying the levels of all transcripts in the worm by highlighting previously unknown transcription differences in immunomodulatory genes between worm sexes and lifecycle stages. Finally, the level of transcriptional sexual dimorphism we observe in this species (as well as in *H. contortus* and *C. elegans*) highlights the need to consider male/female differences in the worms in future experiments with dioecious nematodes.

## Materials and Methods

### Mice and parasites

Male C57Bl/6 mice aged 6-8 weeks (bred and maintained at the animal care facility, Department of Biological Sciences, University of Calgary) were used. All animal experiments were approved by the University of Calgary’s Life and Environmental Sciences Animal Care Committee (protocol AC17-0083). All protocols for animal use and euthanasia were in accordance with the Canadian Council for Animal Care (Canada). Infected mice were orally gavaged with 300 third stage *Heligmosomoides bakeri* larvae (maintained in house, original stock was a gift from Dr. Lisa Reynolds, University of Victoria, Canada) and euthanized at either 5, 7, 10, or 21 days post initial infection. Worms were removed from the intestinal tract, placed in Dulbecco’s modified eagle’s medium – high glucose (Sigma cat. D5796) where they were sexed and counted. The number of worms in each sample is shown in Table 1. Worms were snap frozen and kept at -80°C until RNA isolation.

### RNA isolation and quality control

Worms were lysed by adding 100 μL of Trizol and homogenizing in dry ice three times, for a total of three freeze-thaw cycles and a final volume of 300 μL. The mixture was centrifuged and the supernatant was collected into a new tube where RNA was extracted using the Zymo Direct-zol RNA Miniprep kit (Cat No. R2050) following the manufacturer’s instructions. RNA was digested once with DNase during the isolation, which has been found to sufficiently deplete DNA so as to not interfere with downstream RNA-seq analyses [50]. The RNA was then cleaned using the Zymo RNA Clean and Concentrator -5 kit (Cat No. R1015) following the manufacturer’s instructions. Quantity and quality of the total RNA was assessed on an Agilent 2200 TapeStation RNA ScreenTape following the manufacturer’s instructions.

### Library preparation and sequencing

Libraries were made from suitable RNA samples using the NEBNext Ultra II Directional RNA Library Prep Kit following the manufacturer’s instructions. Libraries were multiplexed and paired-end sequenced on an Illumina NovaSeq with a S2 flow cell or SP flow cell for 300 cycles (2×150bp) using the v1.5 reagent kit, following the manufacturer’s instructions. The resulting reads were deposited in the SRA under the accession number PRJNA750155.

### Species assignment

Ten million reads were randomly selected from all the libraries using seqtk [cite]. They were mapped to genome assemblies for *H. bakeri* (GCA_947359475.1_nxHelBake1.1, Lewis Stevens personal communication) and *H. polygyrus* (GCA_947396885.1_ngHelPoly1.1, Lewis Stevens personal communication) using STAR v2.7.10b with default parameters [51]. The STAR alignment score is recorded in the ‘AS’ for each read and a custom Perl script was used to determine to which assembly each read mapped better to.

### Aligning RNA-seq reads and counting reads per transcript

RNA-seq reads were aligned to the *H. bakeri* genome assembly obtained from WormBase ParaSite (PRJEB15396) using the splice-aware aligner STAR v2.7.3a [51]. The resulting BAM alignment file was used as input for the program featureCounts v2.0.3 [52] to count the fragments overlapping each transcript. Second stranded counts were used only on read pairs with both ends aligned in the proper orientation and fragments overlapping multiple transcripts were fractionally counted among all matches. RNA-seq reads were aligned to the transcript sequences predicted by the annotation accompanying the *H. bakeri* genome assembly PRJEB15396 using the non-splice-aware aligner bwa v0.7.17-r1188. The commands used in all analyses can be found in S2 File.

### Differential gene expression analysis

Read counts per transcript were rounded to the nearest integer and differential transcript expression was analyzed using DESeq2 v1.30.1 [53]. Within each pairwise comparison, transcripts with a false discovery rate adjusted p-value of less than 0.05 were considered differentially expressed.

For male / female comparisons (MS, FS, URM, and URF), pairwise comparisons considered included D5M vs D5F, D7M vs D7F, D10M vs D10F, and D21M vs D21F. Transcripts that were differentially expressed in all four pairwise comparisons were selected. Those that had ≥ 10 fold higher expression in the male samples of the pairwise comparisons were retained as the MS transcripts. Those that had ≥ 10 fold higher expression in the female samples of the pairwise comparisons were retained as the FS transcripts. Those that had higher expression in the male samples of the pairwise comparisons and were in unsigned network modules M3 or M5 and were in signed network modules M3 or M4 were retained as the URM transcripts. Those that had higher expression in the female samples of the pairwise comparisons and were in unsigned network modules M1 or M9 and were in signed network modules M1, M6, or M9 were retained as the URF transcripts.

For developmental comparisons (URTDM, URTDF, URLDM, and URLDF) counts per transcript in the DESeq2 object were VST-transformed to yield expression estimates for every transcript in every sample. For males and females separately, the expression estimates were then compared between every replicate of D5 and D7 samples vs every replicate of D10 and D21 samples to select all transcripts that were up-regulated in the tissue-dwelling phase or the lumen-dwelling phase. Transcripts were then filtered using the co-expression modules (URTDM – unsigned modules M2, M3, or M6 and signed modules M2, M6, M7, or M8; URLDM – unsigned modules M3, M4, or M5 and signed modules M3, M4, or M5; URTDF – unsigned modules M2, M3, M5, M6, M8, or M10 and signed modules M2, M3, M4, M7, M8, or M10; URLDF – unsigned modules M1, M3, M4, M5, or M7 and signed modules M1, M3, M5, M6, or M9). Transcripts that were differentially expressed in at least one of the D7 vs D10 or D5 vs D21 pairwise comparisons were retained.

### Co-expression network analysis

Read counts per transcript were normalized into fragments per kilobase per million mapped reads (FPKM) using the R package countToFPKM v1.0. The top 75% of transcripts by expression value were then grouped into co-expression modules with the R package CEMiTool v1.14.1 [54] using the FPKM values, using the variance-stabilizing transformation and the pearson correlation. Modules were visualized in R to allow determination of the pattern of module enrichment across the sample groups.

Identification of orthologs in *C. elegans*, Gene ontology functional enrichment analysis, Anthelmintic target identification, TPM calculation

All orthologs between *H. bakeri* and *C. elegans* were retrieved from WormBase Parasite. When transcripts of interest were identified in *H. bakeri* for which there was a one-to-one ortholog in *C. elegans*, the function of the *C. elegans* gene was identified through manual literature search.

GO terms enriched in any set of transcripts of interest were identified using the R package gprofiler2 v0.2.1 [55].

Orthologs of known anthelmintic target genes, as previously identified in *C. elegans* [56], were retrieved from WormBase Parasite. Additional targets of interest were taken from [57]. With the *H. bakeri* locus tags, expression of each target was examined. TPM (transcripts per million) were calculated using a bash script (See S2 file). For average TPMs, arithmetic means of the TPMs for all replicates within a sample group were calculated in R along with the standard error.

### Bead feeding assay

Adult worms were removed from the intestinal tract day 10 post-infection and isolated using a modified Baerman apparatus. They were washed three times with water and placed in Dulbecco’s modified eagle’s medium – high glucose (Sigma cat. D5796) where they were sexed and counted. Worms were either used immediately for feeding assays. Mixed sex groups of worms were incubated with or without beads ((108 beads per 5mL of media, Polysciences Inc., Cat. 18859-1for two days. Worms were euthanised with 70% ethanol. Images were acquired using the Zeiss Axio ZoomV.16 stereoscopic microscope (Bio Core Facility, Department of Biological Sciences, University of Calgary). Brightfield and fluorescence images were taken either under the the PlanNeoFluar Z 1x/0.25 FWD 56mm objective objective with the AxioCam High Resolution colour (HRc) and High-Resolution mono (HRm) cameras. Beads were counted manually by zooming into images by two independent observers.

## Supporting information

SOM figures

## Supporting Information

S1 Text. Supplementary Results and Discussion.

S1 Fig. Principal component biplots of the RNA-seq datasets used in this study. A) Biplot of the first two components from the PCA of the raw counts. B) Biplot of the first two components from the PCA of the VST-transformed counts.

S2 Fig. Hierarchical clustering dendrograms of the datasets used in this study. A) HC using Euclidean distance and average linkage. B) HC using Manhattan distance and average linkage. C) HC using Canberra distance and average linkage.

S3 Fig. Transcript co-expression networks. Normalized enrichment scores, reflecting the expression values of the genes in the module, are plotted for each module returned by the network. Unsigned networks allow strong negative correlations to be connected within a module, whereas signed networks do not. Both are shown.

S4 Fig. Expression of orthologs of genes in *C. elegans* that have been implicated in aging. Scatterplots of the VST-transformed read counts (which estimate expression level of the transcript) for all samples grouped by their age/sex combination. The transcript in *H. bakeri* and ortholog in *C. elegans* are shown above the plot.

S5 Fig. Expression of immunomodulatory genes. The arithmetic mean for each age/sex group of the VST-transformed read counts (which estimate expression level of the transcript) are plotted for every transcript matching the description above each plot. Average expression values for female samples are on the left and transcripts are plotted with an open circle connected by lines. Average expression values for male samples are on the right and transcripts are plotted with closed squares connected by lines.

Symbols and lines are colour coded to indicate the transcript being plotted with each being labelled in the legend to the right of the plot.

S1 File. Supplementary Tables.xlsx. All Supplementary Tables referenced in the main text. S2 File. Compressed file of Linux and R commands used in all analyses.

S3 File. Readcountmatrix_fcssall.xlsx. Matrix of read counts per transcript for all RNA-seq datasets included in this study.

S4 File. Compressed file of all the images used for the bead-feeding assay.

