## Supplementary material for "Transcriptional patterns of sexual dimorphism and in host developmental programs in the model parasitic nematode *Heligmosomoides bakeri*": SOM figures

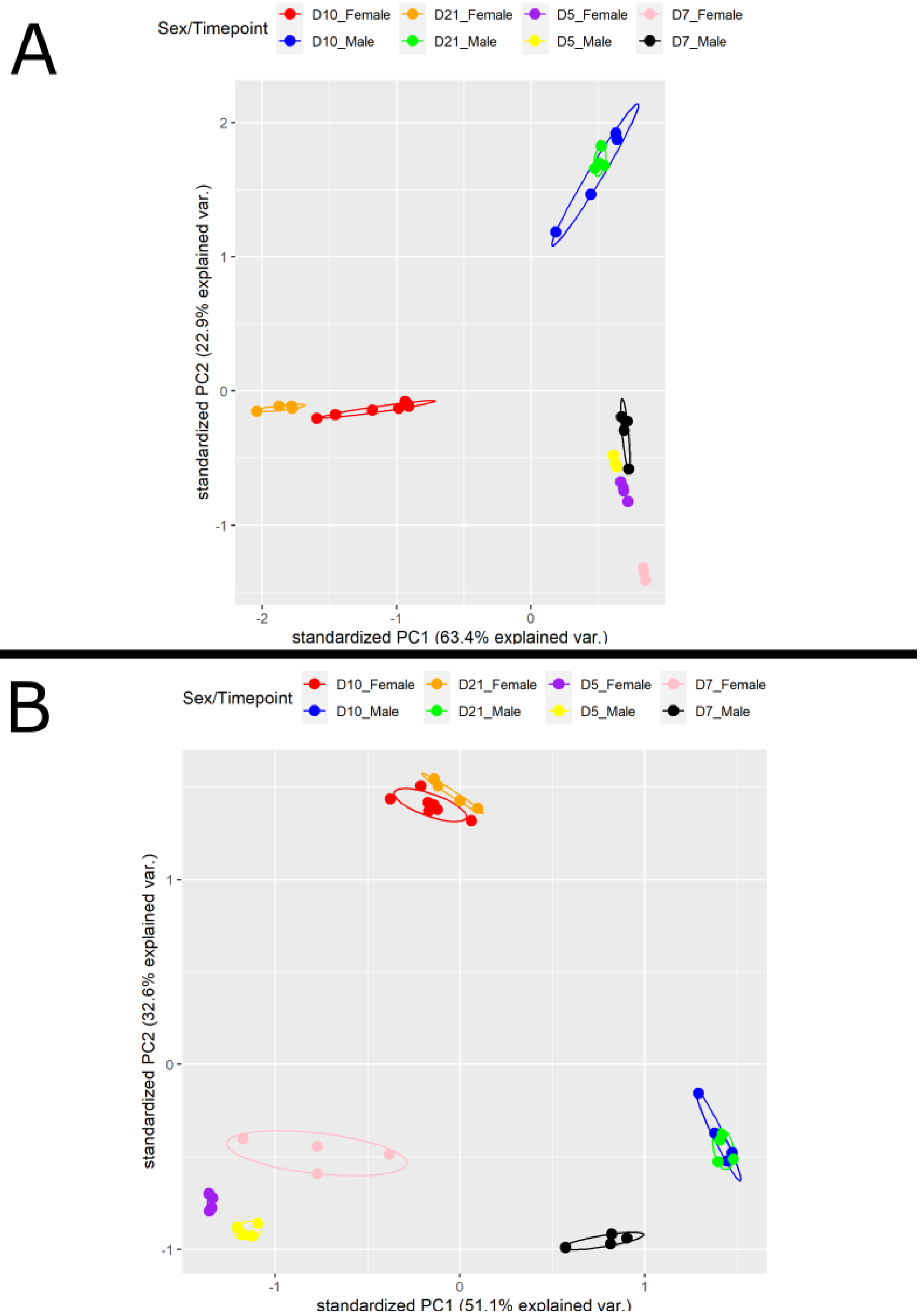


S1 Fig. PCA Biplots

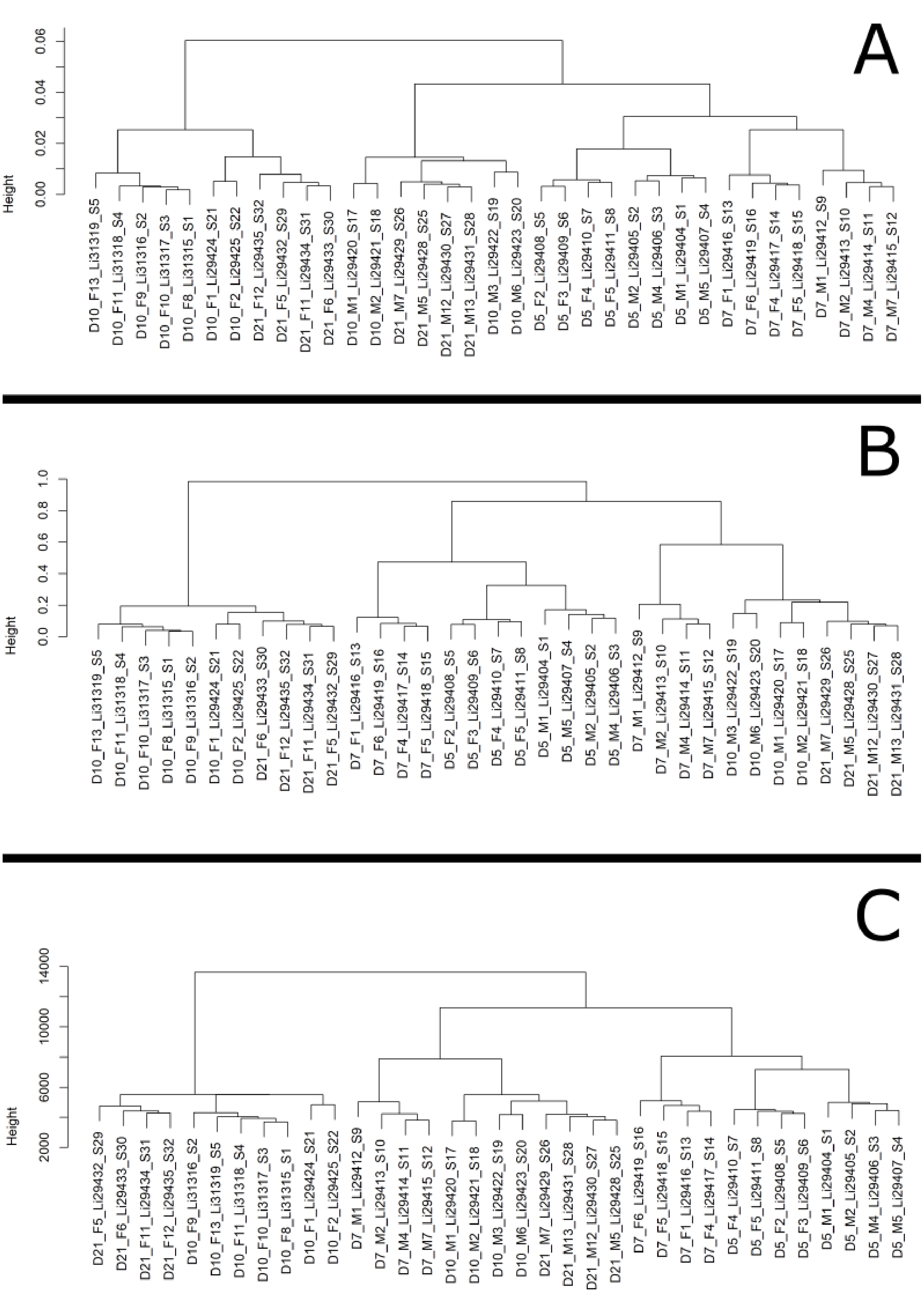


S2 Fig. HC Dendrograms

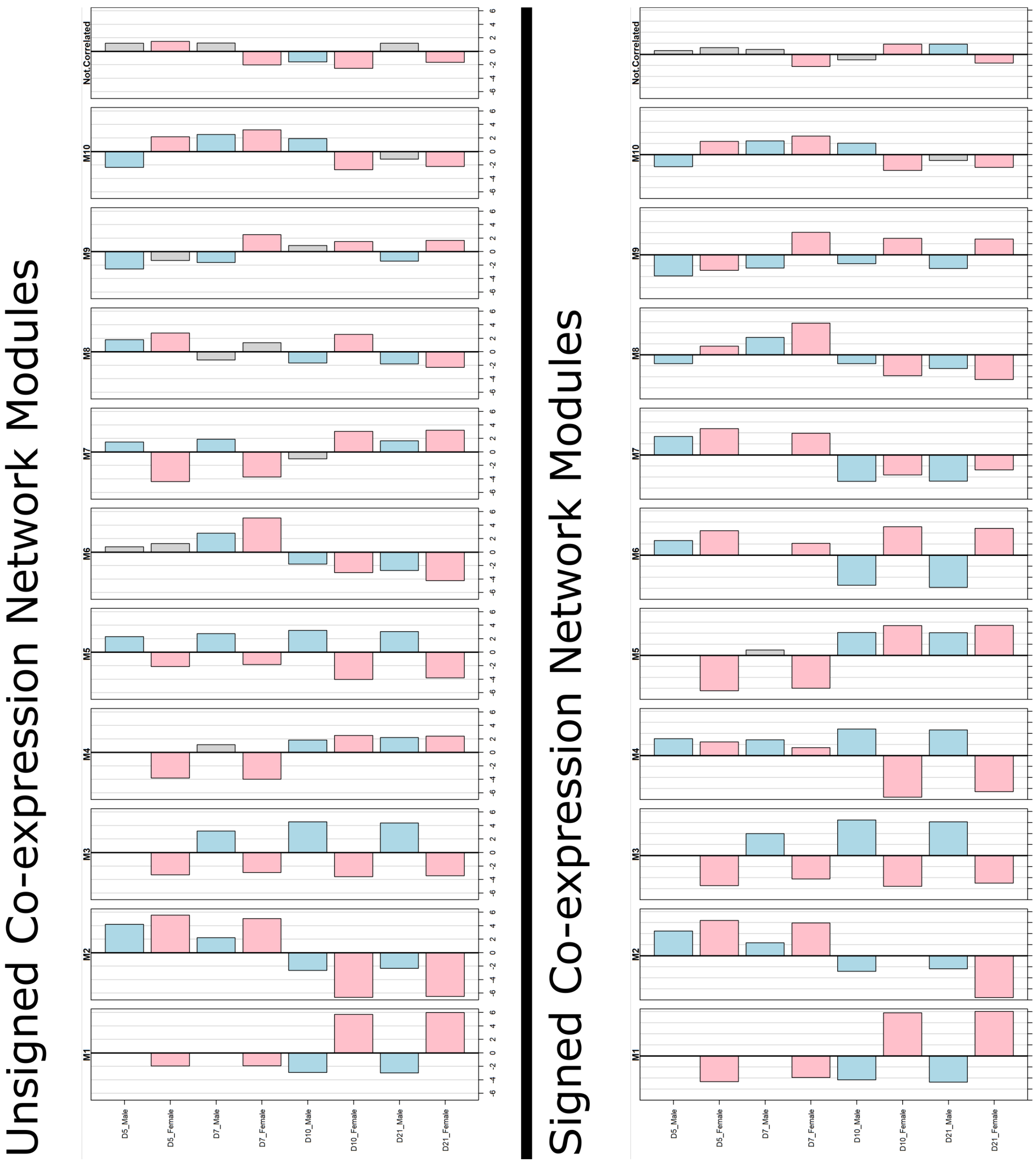


S3 Fig. Coexpression networks

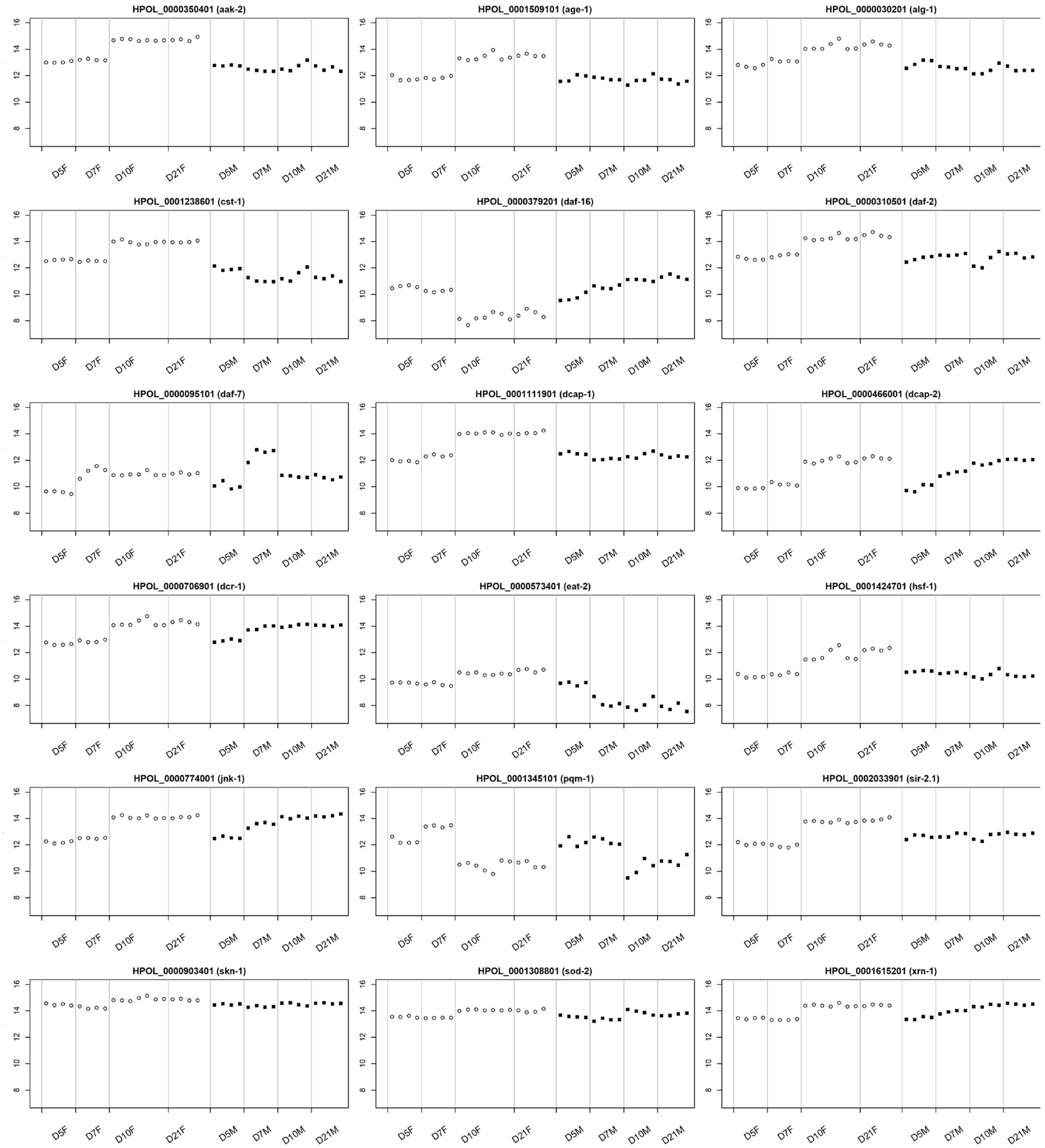


S4 Fig. hbakeri_celeagingorthos_expr

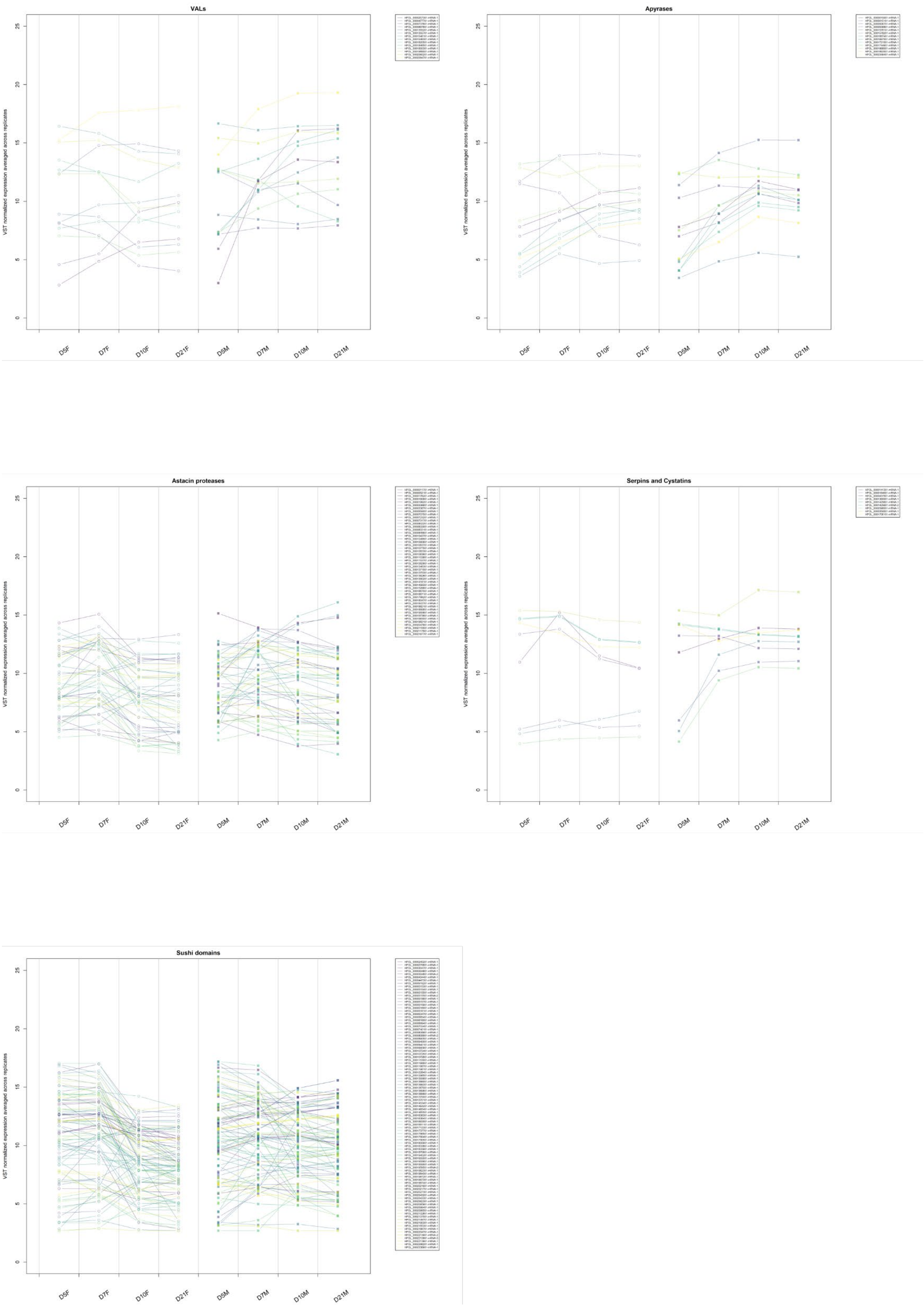


S5 Fig. Immunomod Exprs
