## Supplementary material for "Transcriptional patterns of sexual dimorphism and in host developmental programs in the model parasitic nematode *Heligmosomoides bakeri*": S2 File: calculate_tpms.sh

```
#Use the output of featureCounts for the first argument. This script parses the gene lengths from it

FC_COUNTS=$1
NAME=$2

#Strip first two lines out but put the second line as first line to final output
sed -n '2p' ${FC_COUNTS} | awk 'BEGIN {ORS=""} {print $1; for(i=7; i<=NF; i++){print "\t" $i} } END{print "\n"}' > ${NAME}_tpms.txt
sed '1,2d' ${FC_COUNTS} > c.tmp

awk 'BEGIN {ORS=""}; {print $1; for(i=7; i<=NF; i++){rpk=$i/$6*1000; print "\t" rpk;} print "\n"}' c.tmp > rpks.tmp
COLS=`head -1 rpks.tmp | wc -w`
COLS=$( expr ${COLS} - 1 + 1 )

for ((i=2; i<=${COLS}; i++))
do
	awk -v col=${i} 'BEGIN {ORS=""} {sum+=$col} END{sf=sum/1000000; print sf "\t"}' rpks.tmp >> sums.tmp
done

awk 'BEGIN {ORS=""} NR==FNR{for(i=1; i<=NF; i++){sfs[i]=$i} } {print $1; for(j=2; j<=NF; j++){sfsindex=j-1; tpm=$j/sfs[sfsindex]; print "\t" tpm;} print "\n"}' sums.tmp rpks.tmp > tpms.tmp

sed '1d' tpms.tmp >> ${NAME}_tpms.txt

rm c.tmp
rm rpks.tmp
rm sums.tmp
rm tpms.tmp
```
