## Supplementary material for "Transcriptional patterns of sexual dimorphism and in host developmental programs in the model parasitic nematode *Heligmosomoides bakeri*": S2 File: Hbakeri_drugtargetexpr.Rmd

````
---
title: "Drug_target_expr'"
author: "Stephen Pollo"
date: "02/08/2022"
output: html_document
---

```{r}
# Read in data
vst_exprs_df=readRDS("vst_exprs_df_hbakeri.rds")
hbakeri_WBPSgenandannots_fcssfracoverlapspaircheck_all_tpms <- read.delim("hbakeri_WBPSgenandannots_fcssfracoverlapspaircheck_all_tpms.txt")
res_fc_secondstranded = readRDS("Hbakeri_deseq2_3-21-22_res_fc_secondstranded.rds")
```

```{r}
#Make table

dtartpms <- function(l) {
  favg=data.frame()
  mavg=data.frame()
  qs=vector()
  for(i in 1:length(l)){
    query=paste("transcript:", l[i], sep="")
    if(grepl('-', query, fixed=TRUE)) {} else {query=paste(query, "-mRNA-1", sep="")}
    qs=append(qs, query)
    
    tmp=hbakeri_WBPSgenandannots_fcssfracoverlapspaircheck_all_tpms[hbakeri_WBPSgenandannots_fcssfracoverlapspaircheck_all_tpms$Geneid == query,]
    tmp2=tmp[,c(25:28,33:36,2:6,9:10,17:20)] # Females
    tmp3=tmp[,c(29:32,37:40,11:13,15,21:24)] #Males
    tmp4=c(mean(unlist(tmp2[1:4])), mean(unlist(tmp2[5:8])), mean(unlist(tmp2[9:15])), mean(unlist(tmp2[16:19]))) # Female avgs
    tmp5=c(mean(unlist(tmp3[1:4])), mean(unlist(tmp3[5:8])), mean(unlist(tmp3[9:12])), mean(unlist(tmp3[13:16]))) # Male avgs
    favg=rbind(favg, tmp4)
    mavg=rbind(mavg, tmp5)
  }
  
  colnames(favg)=c("D5_F", "D7_F", "D10_F", "D21_F")
  colnames(mavg)=c("D5_M", "D7_M", "D10_M", "D21_M")
  favg$gene=qs
  mavg$gene=qs
  
  tmp=list()
  tmp[[1]]=favg
  tmp[[2]]=mavg
  names(tmp)=c("Favgs", "Mavgs")
  
  return(tmp)
}

getadjpvals <- function(l) {
  fvals=data.frame()
  mvals=data.frame()
  qs=vector()
  for(i in 1:length(l)){
    query=paste("transcript:", l[i], sep="")
    if(grepl('-', query, fixed=TRUE)) {} else {query=paste(query, "-mRNA-1", sep="")}
    qs=append(qs, query)
    
    # Get female adj pvals
    tmp=vector()
    tmp=append(tmp, query)
    tmp=append(tmp, res_fc_secondstranded$D5_Female_vs_D7_Female[rownames(res_fc_secondstranded$D5_Female_vs_D7_Female) == query,6])
    tmp=append(tmp, res_fc_secondstranded$D10_Female_vs_D7_Female[rownames(res_fc_secondstranded$D10_Female_vs_D7_Female) == query,6])
    tmp=append(tmp, res_fc_secondstranded$D10_Female_vs_D21_Female[rownames(res_fc_secondstranded$D10_Female_vs_D21_Female) == query,6])
    
    tmp2=vector()
    tmp2=append(tmp2, query)
    tmp2=append(tmp2, res_fc_secondstranded$D5_Male_vs_D7_Male[rownames(res_fc_secondstranded$D5_Male_vs_D7_Male) == query,6])
    tmp2=append(tmp2, res_fc_secondstranded$D10_Male_vs_D7_Male[rownames(res_fc_secondstranded$D10_Male_vs_D7_Male) == query,6])
    tmp2=append(tmp2, res_fc_secondstranded$D10_Male_vs_D21_Male[rownames(res_fc_secondstranded$D10_Male_vs_D21_Male) == query,6])
    
    fvals=rbind(fvals, tmp)
    mvals=rbind(mvals, tmp2)
  }
  
  colnames(fvals)=c("Query", "D5F_vs_D7F", "D7F_vs_D10F", "D10F_vs_D21F")
  colnames(mvals)=c("Query", "D5M_vs_D7M", "D7M_vs_D10M", "D10M_vs_D21M")
  
  tmp=list()
  tmp[[1]]=fvals
  tmp[[2]]=mvals
  names(tmp)=c("Fvals", "Mvals")
  
  return(tmp)
}

genes=c("HPOL_0001177601", "HPOL_0001003101", "HPOL_0001839701", "HPOL_0000126101", "HPOL_0000096601", "HPOL_0001177601", "HPOL_0000247101", "HPOL_0001059501", "HPOL_0000812201", "HPOL_0000079901", "HPOL_0001278501", "HPOL_0001278801", "HPOL_0002208401", "HPOL_0000080001", "HPOL_0000053201", "HPOL_0000939601", "HPOL_0000939601", "HPOL_0000510801", "HPOL_0000510601", "HPOL_0000510701", "HPOL_0000350101", "HPOL_0001283901", "HPOL_0000661601", "HPOL_0000662001", "HPOL_0000292001", "HPOL_0000775801", "HPOL_0001846201", "HPOL_0000573301", "HPOL_0000675901", "HPOL_0000717801", "HPOL_0000350501", "HPOL_0000332701", "HPOL_0001927801", "HPOL_0002147401", "HPOL_0002202601", "HPOL_0002243501", "HPOL_0001923201", "HPOL_0000318201", "HPOL_0000014001", "HPOL_0000877201", "HPOL_0000032901", "HPOL_0000343501", "HPOL_0001450601")

t1=dtartpms(genes)
t2=getadjpvals(genes)

t3=merge(t1$Favgs, t2$Fvals, by.x=0, by.y=0)
t4=merge(t1$Mavgs, t2$Mvals, by.x=0, by.y=0)

ftable=t3[,c(7,2,8,3,9,4,10,5,1)]
mtable=t4[,c(7,2,8,3,9,4,10,5,1)]

```
````
