## Supplementary material for "Transcriptional patterns of sexual dimorphism and in host developmental programs in the model parasitic nematode *Heligmosomoides bakeri*": S2 File: Hbakeri_immunomod_mainfig.Rmd

````
---
title: "Hbakeri_immunomod_mainfig"
author: "Stephen Pollo"
date: "09/08/2022"
output: html_document
---

```{r}
# Read in data
hbakeri_WBPSgenandannots_fcssfracoverlapspaircheck_all_tpms <- read.delim("C:/Users/Stephen/Desktop/hbakeri_WBPSgenandannots_fcssfracoverlapspaircheck_all_tpms.txt")
```

```{r}
# Get avg TPMs

dtartpms <- function(l) {
  favg=data.frame()
  mavg=data.frame()
  qs=vector()
  for(i in 1:length(l)){
    query=paste("transcript:", l[i], sep="")
    if(grepl('-', query, fixed=TRUE)) {} else {query=paste(query, "-mRNA-1", sep="")}
    qs=append(qs, query)
    
    tmp=hbakeri_WBPSgenandannots_fcssfracoverlapspaircheck_all_tpms[hbakeri_WBPSgenandannots_fcssfracoverlapspaircheck_all_tpms$Geneid == query,]
    tmp2=tmp[,c(25:28,33:36,2:6,9:10,17:20)] # Females
    tmp3=tmp[,c(29:32,37:40,11:13,15,21:24)] #Males
    tmp4=c(mean(unlist(tmp2[1:4])), mean(unlist(tmp2[5:8])), mean(unlist(tmp2[9:15])), mean(unlist(tmp2[16:19]))) # Female avgs
    tmp5=c(mean(unlist(tmp3[1:4])), mean(unlist(tmp3[5:8])), mean(unlist(tmp3[9:12])), mean(unlist(tmp3[13:16]))) # Male avgs
    favg=rbind(favg, tmp4)
    mavg=rbind(mavg, tmp5)
  }
  
  colnames(favg)=c("D5_F", "D7_F", "D10_F", "D21_F")
  colnames(mavg)=c("D5_M", "D7_M", "D10_M", "D21_M")
  favg$gene=qs
  mavg$gene=qs
  
  tmp=list()
  tmp[[1]]=favg
  tmp[[2]]=mavg
  names(tmp)=c("Favgs", "Mavgs")
  
  return(tmp)
}

# Get standard errors of avg TPMs
dtarsds <- function(l) {
     fsd=data.frame()
     msd=data.frame()
     qs=vector()
     for(i in 1:length(l)){
         query=paste("transcript:", l[i], sep="")
         if(grepl('-', query, fixed=TRUE)) {} else {query=paste(query, "-mRNA-1", sep="")}
         qs=append(qs, query)
         
         tmp=hbakeri_WBPSgenandannots_fcssfracoverlapspaircheck_all_tpms[hbakeri_WBPSgenandannots_fcssfracoverlapspaircheck_all_tpms$Geneid == query,]
         tmp2=tmp[,c(25:28,33:36,2:6,9:10,17:20)] # Females
         tmp3=tmp[,c(29:32,37:40,11:13,15,21:24)] #Males
         tmp4=c(sd(unlist(tmp2[1:4]))/sqrt(4), sd(unlist(tmp2[5:8]))/sqrt(4), sd(unlist(tmp2[9:15]))/sqrt(7), sd(unlist(tmp2[16:19]))/sqrt(4)) # Female ses
         tmp5=c(sd(unlist(tmp3[1:4]))/sqrt(4), sd(unlist(tmp3[5:8]))/sqrt(4), sd(unlist(tmp3[9:12]))/sqrt(4), sd(unlist(tmp3[13:16]))/sqrt(4)) # Male ses
         fsd=rbind(fsd, tmp4)
         msd=rbind(msd, tmp5)
     }
     
     colnames(fsd)=c("D5_F", "D7_F", "D10_F", "D21_F")
     colnames(msd)=c("D5_M", "D7_M", "D10_M", "D21_M")
     fsd$gene=qs
     msd$gene=qs
     
     tmp=list()
     tmp[[1]]=fsd
     tmp[[2]]=msd
     names(tmp)=c("Fses", "Mses")
     
     return(tmp)
}

# Not including HPOL_0002154401
genes=c("HPOL_0001636401", "HPOL_0001228301", "HPOL_0001228401", "HPOL_0001864701")

```

```{r}
# Get vals and plot
avgs=dtartpms(genes)
ses=dtarsds(genes)

x=c(1:4)

```

```{r}
# Draw plot
png(file="hbakeri_immunomod_exprfig_updated.png", width=2000, height=1800, res=300)

par(mfrow=c(3,2))
par(mar=c(2,5,2,1))
plot(x, avgs$Favgs[1,1:4], ylim=c(0,1200), pch=19, xlab="Days Post Infection", ylab="Average TPM +/- SE", main="Female expression of HbARI", xaxt='n')
axis(1, at=1:4, labels=c("5", "7", "10", "21"))
arrows(x, unlist(avgs$Favgs[1,1:4]-ses$Fses[1,1:4]), x, unlist(avgs$Favgs[1,1:4]+ses$Fses[1,1:4]), length=0.05, angle = 90, code=3)

plot(x, avgs$Mavgs[1,1:4], ylim=c(0,1200), pch=19, xlab="Days Post Infection", ylab="Average TPM +/- SE", main="Male expression of HbARI", xaxt='n')
axis(1, at=1:4, labels=c("5", "7", "10", "21"))
arrows(x, unlist(avgs$Mavgs[1,1:4]-ses$Mses[1,1:4]), x, unlist(avgs$Mavgs[1,1:4]+ses$Mses[1,1:4]), length=0.05, angle = 90, code=3)

plot(x, avgs$Favgs[2,1:4], ylim=c(0,2500), pch=19, xlab="Days Post Infection", ylab="Average TPM +/- SE", main="Female expression of HbBARIs", xaxt='n')
axis(1, at=1:4, labels=c("5", "7", "10", "21"))
arrows(x, unlist(avgs$Favgs[2,1:4]-ses$Fses[2,1:4]), x, unlist(avgs$Favgs[2,1:4]+ses$Fses[2,1:4]), length=0.05, angle = 90, code=3)
points(x, avgs$Favgs[3,1:4], pch=16, col="red")
arrows(x, unlist(avgs$Favgs[3,1:4]-ses$Fses[3,1:4]), x, unlist(avgs$Favgs[3,1:4]+ses$Fses[3,1:4]), length=0.05, angle = 90, code=3, col="red")
legend("topright", legend=c("BARI", "BARI_Hom2"), box.lty=1, pch=c(16,16), col=c("black", "red"))

plot(x, avgs$Mavgs[2,1:4], ylim=c(0,2500), pch=19, xlab="Days Post Infection", ylab="Average TPM +/- SE", main="Male expression of HbBARIs", xaxt='n')
axis(1, at=1:4, labels=c("5", "7", "10", "21"))
arrows(x, unlist(avgs$Mavgs[2,1:4]-ses$Mses[2,1:4]), x, unlist(avgs$Mavgs[2,1:4]+ses$Mses[2,1:4]), length=0.05, angle = 90, code=3)
points(x, avgs$Mavgs[3,1:4], pch=16, col="red")
arrows(x, unlist(avgs$Mavgs[3,1:4]-ses$Mses[3,1:4]), x, unlist(avgs$Mavgs[3,1:4]+ses$Mses[3,1:4]), length=0.05, angle = 90, code=3, col="red")
legend("topright", legend=c("BARI", "BARI_Hom2"), box.lty=1, pch=c(16,16), col=c("black", "red"))

plot(x, avgs$Favgs[4,1:4], ylim=c(0,100), pch=19, xlab="Days Post Infection", ylab="Average TPM +/- SE", main="Female expression of HbTGM-6", xaxt='n')
axis(1, at=1:4, labels=c("5", "7", "10", "21"))
arrows(x, unlist(avgs$Favgs[4,1:4]-ses$Fses[4,1:4]), x, unlist(avgs$Favgs[4,1:4]+ses$Fses[4,1:4]), length=0.05, angle = 90, code=3)

plot(x, avgs$Mavgs[4,1:4], ylim=c(0,100), pch=19, xlab="Days Post Infection", ylab="Average TPM +/- SE", main="Male expression of HbTGM-6", xaxt='n')
axis(1, at=1:4, labels=c("5", "7", "10", "21"))
arrows(x, unlist(avgs$Mavgs[4,1:4]-ses$Mses[4,1:4]), x, unlist(avgs$Mavgs[4,1:4]+ses$Mses[4,1:4]), length=0.05, angle = 90, code=3)

dev.off()

```

```{r}
# Aging genes plot
genes2=c("HPOL_0000310501", "HPOL_0001509101", "HPOL_0000379201", "HPOL_0000350401")

avgs2=dtartpms(genes2)
ses2=dtarsds(genes2)

x=c(1:4)
```

```{r}
# Draw aging plot
png(file="hbakeri_aging_exprfig_updated.png", width=2200, height=1800, res=300)

par(mfrow=c(4,2))
par(mar=c(2,5,2,1))
plot(x, avgs2$Favgs[1,1:4], ylim=c(0,100), pch=19, xlab="Days Post Infection", ylab="Average TPM +/- SE", main="Female expression of HPOL_0000310501 (daf-2)", xaxt='n')
axis(1, at=1:4, labels=c("5", "7", "10", "21"))
arrows(x, unlist(avgs2$Favgs[1,1:4]-ses2$Fses[1,1:4]), x, unlist(avgs2$Favgs[1,1:4]+ses2$Fses[1,1:4]), length=0.05, angle = 90, code=3)

plot(x, avgs2$Mavgs[1,1:4], ylim=c(0,100), pch=19, xlab="Days Post Infection", ylab="Average TPM +/- SE", main="Male expression of HPOL_0000310501 (daf-2)", xaxt='n')
axis(1, at=1:4, labels=c("5", "7", "10", "21"))
arrows(x, unlist(avgs2$Mavgs[1,1:4]-ses2$Mses[1,1:4]), x, unlist(avgs2$Mavgs[1,1:4]+ses2$Mses[1,1:4]), length=0.05, angle = 90, code=3)

plot(x, avgs2$Favgs[2,1:4], ylim=c(0,50), pch=19, xlab="Days Post Infection", ylab="Average TPM +/- SE", main="Female expression of HPOL_0001509101 (age-1)", xaxt='n')
axis(1, at=1:4, labels=c("5", "7", "10", "21"))
arrows(x, unlist(avgs2$Favgs[2,1:4]-ses2$Fses[2,1:4]), x, unlist(avgs2$Favgs[2,1:4]+ses2$Fses[2,1:4]), length=0.05, angle = 90, code=3)

plot(x, avgs2$Mavgs[2,1:4], ylim=c(0,50), pch=19, xlab="Days Post Infection", ylab="Average TPM +/- SE", main="Male expression of HPOL_0001509101 (age-1)", xaxt='n')
axis(1, at=1:4, labels=c("5", "7", "10", "21"))
arrows(x, unlist(avgs2$Mavgs[2,1:4]-ses2$Mses[2,1:4]), x, unlist(avgs2$Mavgs[2,1:4]+ses2$Mses[2,1:4]), length=0.05, angle = 90, code=3)

plot(x, avgs2$Favgs[3,1:4], ylim=c(0,50), pch=19, xlab="Days Post Infection", ylab="Average TPM +/- SE", main="Female expression of HPOL_0000379201 (daf-16)", xaxt='n')
axis(1, at=1:4, labels=c("5", "7", "10", "21"))
arrows(x, unlist(avgs2$Favgs[3,1:4]-ses2$Fses[3,1:4]), x, unlist(avgs2$Favgs[3,1:4]+ses2$Fses[3,1:4]), length=0.05, angle = 90, code=3)

plot(x, avgs2$Mavgs[3,1:4], ylim=c(0,50), pch=19, xlab="Days Post Infection", ylab="Average TPM +/- SE", main="Male expression of HPOL_0000379201 (daf-16)", xaxt='n')
axis(1, at=1:4, labels=c("5", "7", "10", "21"))
arrows(x, unlist(avgs2$Mavgs[3,1:4]-ses2$Mses[3,1:4]), x, unlist(avgs2$Mavgs[3,1:4]+ses2$Mses[3,1:4]), length=0.05, angle = 90, code=3)

plot(x, avgs2$Favgs[4,1:4], ylim=c(0,100), pch=19, xlab="Days Post Infection", ylab="Average TPM +/- SE", main="Female expression of HPOL_0000350401 (aak-2)", xaxt='n')
axis(1, at=1:4, labels=c("5", "7", "10", "21"))
arrows(x, unlist(avgs2$Favgs[4,1:4]-ses2$Fses[4,1:4]), x, unlist(avgs2$Favgs[4,1:4]+ses2$Fses[4,1:4]), length=0.05, angle = 90, code=3)

plot(x, avgs2$Mavgs[4,1:4], ylim=c(0,100), pch=19, xlab="Days Post Infection", ylab="Average TPM +/- SE", main="Male expression of HPOL_0000350401 (aak-2)", xaxt='n')
axis(1, at=1:4, labels=c("5", "7", "10", "21"))
arrows(x, unlist(avgs2$Mavgs[4,1:4]-ses2$Mses[4,1:4]), x, unlist(avgs2$Mavgs[4,1:4]+ses2$Mses[4,1:4]), length=0.05, angle = 90, code=3)

dev.off()

```
````
