## Supplementary material for "Transcriptional patterns of sexual dimorphism and in host developmental programs in the model parasitic nematode *Heligmosomoides bakeri*": S2 File: hbakeri_infection_biplots.Rmd

````
---
title: "H. bakeri infections transcriptomes biplots"
author: "Stephen Pollo"
date: "05/08/2021"
output: html_document
---

```{r}
# Load libraries
library(ggbiplot)

# Import the matrices
hbakeri_featurecount_output <- read.table("hbakeri_WBPSgenandannots_fc_ss_all_fractionoverlaps_paircheck.txt", row.names=1, comment.char = "#")

# Fix col names
colnames(hbakeri_featurecount_output) = c("Chr","Start","End","Strand","Length", "D10_F10_Li31317_S3", "D10_F11_Li31318_S4", "D10_F13_Li31319_S5", "D10_F1_Li29424_S21","D10_F2_Li29425_S22","D10_F6_Li29426_S23","D10_F7_Li29427_S24", "D10_F8_Li31315_S1", "D10_F9_Li31316_S2", "D10_M1_Li29420_S17","D10_M2_Li29421_S18","D10_M3_Li29422_S19", "D10_M3_Li31320_S6","D10_M6_Li29423_S20", "D10_M6_Li31321_S7","D21_F11_Li29434_S31","D21_F12_Li29435_S32","D21_F5_Li29432_S29","D21_F6_Li29433_S30","D21_M12_Li29430_S27","D21_M13_Li29431_S28","D21_M5_Li29428_S25","D21_M7_Li29429_S26","D5_F2_Li29408_S5","D5_F3_Li29409_S6","D5_F4_Li29410_S7","D5_F5_Li29411_S8","D5_M1_Li29404_S1","D5_M2_Li29405_S2","D5_M4_Li29406_S3","D5_M5_Li29407_S4","D7_F1_Li29416_S13","D7_F4_Li29417_S14","D7_F5_Li29418_S15","D7_F6_Li29419_S16","D7_M1_Li29412_S9","D7_M2_Li29413_S10","D7_M4_Li29414_S11","D7_M7_Li29415_S12")
featurecounts_second_stranded_counts_all=as.data.frame(apply(hbakeri_featurecount_output[-1,-1:-5], 2, as.numeric))
rownames(featurecounts_second_stranded_counts_all)=rownames(hbakeri_featurecount_output[-1,])

```

```{r}
# Functions

# Create matrix for ggbiplot
createGGMatrix <- function(counts_file) {
  tmp = t(counts_file)
  
  # GGbiplot doesn't like genes that are all 0 so they have to be removed
  tmp2 = tmp[,apply(tmp, 2, var, na.rm=TRUE) != 0]
  
  return(tmp2)
}

# Draw the biplots
drawMyBiplot <- function(pca_obj) {
  ggbiplot(pca_obj, ellipse=TRUE, groups=groups, var.axes=FALSE) + 
    scale_color_manual(name="Sex/Timepoint", values=c("darkred", "red", "blue", "darkblue", "orange", "green", "purple", "yellow", "pink", "black")) +
    scale_shape_manual(name="Sex/Timepoint", values=c(19,19,19,19,19,19,19,19,19,19)) +
    geom_point(aes(colour=groups, shape=groups), size=3) +
    theme(legend.direction="horizontal", legend.position="top")
}

```

``` {r}
# Normalize the counts by library size
fcss_normby_assigned=apply(featurecounts_second_stranded_counts_all, 2, norm<-function(x){return (x/sum(x))})

# Create required matrix
groups=c(rep("D10_Female_new",3), rep("D10_Female_old",4), rep("D10_Female_new",2), rep("D10_Male_old",3), rep("D10_Male_new",1), rep("D10_Male_old",1), rep("D10_Male_new",1), rep("D21_Female",4), rep("D21_Male",4), rep("D5_Female",4), rep("D5_Male",4), rep("D7_Female",4), rep("D7_Male",4))

bip_fcss = createGGMatrix(fcss_normby_assigned)

# Run PCA
pca_fcss = prcomp(bip_fcss)

```

```{r}
# Export biplots to files

png(file="hbakeri_pca_39_infection_transcriptomes.png", width=2000, height=1800, res=300)
drawMyBiplot(pca_fcss)
dev.off()
```

```{r}
# Biplots with vectors

library(factoextra)

# Use the same PCA from above
fviz_pca_biplot(pca_fcss, repel = TRUE, select.var = list(contrib=10))
```

```{r}
# Run hierarchical clustering analysis

rev=t(fcss_normby_assigned)
#rownames(rev)=c("D10_F1","D10_F2","D10_F6","D10_F7","D10_M1","D10_M2","D10_M3","D10_M6","D21_F11","D21_F12","D21_F5","D21_F6","D21_M12","D21_M13","D21_M5","D21_M7","D5_F2","D5_F3","D5_F4","D5_F5","D5_M1","D5_M2","D5_M4","D5_M5","D7_F1","D7_F4","D7_F5","D7_F6","D7_M1","D7_M2","D7_M4","D7_M7")

dm_euc=dist(rev, method = 'euclidean')
dm_man=dist(rev, method = 'manhattan')
dm_can=dist(rev, method = 'canberra')

hclust_euc_avg=hclust(dm_euc, method = 'average')
hclust_man_avg=hclust(dm_man, method = 'average')
hclust_can_avg=hclust(dm_can, method = 'average')

```

```{r}
# Write hierarchical clustering plots to files

png(file="hbakeri_hc_avglinkage_euclideandist.png", width=3600, height=1800, res=300)
plot(hclust_euc_avg)
dev.off()

png(file="hbakeri_hc_avglinkage_manhattandist.png", width=3600, height=1800, res=300)
plot(hclust_man_avg)
dev.off()

png(file="hbakeri_hc_avglinkage_canberradist.png", width=3600, height=1800, res=300)
plot(hclust_can_avg)
dev.off()

```
````
