## Supplementary material for "Transcriptional patterns of sexual dimorphism and in host developmental programs in the model parasitic nematode *Heligmosomoides bakeri*": S2 File: hbakeri_infection_coexpression_networks.Rmd

````
---
title: "H. bakeri co-expression networks"
author: "Stephen Pollo"
date: "16/08/2021"
output: html_document
---

```{r}
# Load libraries and import data
library(CEMiTool)
library(countToFPKM)

hbakeri_featurecount_output <- read.table("hbakeri_WBPSgenandannots_fc_ss_all_fractionoverlaps_paircheck.txt", row.names=1, comment.char = "#")

# Fix col names
colnames(hbakeri_featurecount_output) = c("Chr","Start","End","Strand","Length", "D10_F10_Li31317_S3", "D10_F11_Li31318_S4", "D10_F13_Li31319_S5", "D10_F1_Li29424_S21","D10_F2_Li29425_S22","D10_F6_Li29426_S23","D10_F7_Li29427_S24", "D10_F8_Li31315_S1", "D10_F9_Li31316_S2", "D10_M1_Li29420_S17","D10_M2_Li29421_S18","D10_M3_Li29422_S19", "D10_M3_Li31320_S6","D10_M6_Li29423_S20", "D10_M6_Li31321_S7","D21_F11_Li29434_S31","D21_F12_Li29435_S32","D21_F5_Li29432_S29","D21_F6_Li29433_S30","D21_M12_Li29430_S27","D21_M13_Li29431_S28","D21_M5_Li29428_S25","D21_M7_Li29429_S26","D5_F2_Li29408_S5","D5_F3_Li29409_S6","D5_F4_Li29410_S7","D5_F5_Li29411_S8","D5_M1_Li29404_S1","D5_M2_Li29405_S2","D5_M4_Li29406_S3","D5_M5_Li29407_S4","D7_F1_Li29416_S13","D7_F4_Li29417_S14","D7_F5_Li29418_S15","D7_F6_Li29419_S16","D7_M1_Li29412_S9","D7_M2_Li29413_S10","D7_M4_Li29414_S11","D7_M7_Li29415_S12")
featurecounts_second_stranded_counts_all=as.data.frame(apply(hbakeri_featurecount_output[-1,-1:-5], 2, as.numeric))
rownames(featurecounts_second_stranded_counts_all)=rownames(hbakeri_featurecount_output[-1,])

# Remove the samples that are being excluded
keeps=c("D10_F10_Li31317_S3", "D10_F11_Li31318_S4", "D10_F13_Li31319_S5", "D10_F1_Li29424_S21", "D10_F2_Li29425_S22", "D10_F8_Li31315_S1", "D10_F9_Li31316_S2", "D10_M1_Li29420_S17","D10_M2_Li29421_S18","D10_M3_Li29422_S19", "D10_M6_Li29423_S20", "D21_F11_Li29434_S31", "D21_F12_Li29435_S32", "D21_F5_Li29432_S29", "D21_F6_Li29433_S30", "D21_M12_Li29430_S27", "D21_M13_Li29431_S28", "D21_M5_Li29428_S25", "D21_M7_Li29429_S26", "D5_F2_Li29408_S5", "D5_F3_Li29409_S6", "D5_F4_Li29410_S7", "D5_F5_Li29411_S8", "D5_M1_Li29404_S1", "D5_M2_Li29405_S2", "D5_M4_Li29406_S3", "D5_M5_Li29407_S4", "D7_F1_Li29416_S13", "D7_F4_Li29417_S14", "D7_F5_Li29418_S15", "D7_F6_Li29419_S16", "D7_M1_Li29412_S9", "D7_M2_Li29413_S10", "D7_M4_Li29414_S11", "D7_M7_Li29415_S12")
featurecounts_second_stranded_counts_all=featurecounts_second_stranded_counts_all[keeps]

```

```{r}
# Normalize raw counts into FPKM
# Need gene lengths (calculated on ARC)
h_bakeri_gene_lengths <- read.csv("h_bakeri_gene_lengths.csv", header=FALSE)
rownames(h_bakeri_gene_lengths) = h_bakeri_gene_lengths[,1]
lengths=h_bakeri_gene_lengths[,-1]

# Need average read lengths per library (taken from STAR mapping final log)
average_read_lengths <- read.table("average_read_lengths.txt", quote="\"", comment.char="")
read_lengths=average_read_lengths[,-2:-6]
rownames(read_lengths) = read_lengths[,1]
read_lengths=read_lengths[keeps,]
read_lengths=read_lengths[,-1]

# Calculate fpkms and transform to data frame for CEMiTool
fpkms=fpkm(featurecounts_second_stranded_counts_all, lengths, read_lengths)
fpkms_df=as.data.frame(fpkms)

```

```{r}
# Run CEMiTool
# Build sample annotation data frame
snames=colnames(featurecounts_second_stranded_counts_all)
groups=c(rep("D10_Female",7), rep("D10_Male",4), rep("D21_Female",4), rep("D21_Male",4), rep("D5_Female",4), rep("D5_Male",4), rep("D7_Female",4), rep("D7_Male",4))
groups2=c(rep("Female",7), rep("Male",4), rep("Female",4), rep("Male",4), rep("Female",4), rep("Male",4), rep("Female",4), rep("Male",4))
groups3=c(rep("Lumen",19), rep("Tissue",16))
groups4=c(rep("D10",11), rep("D21",8), rep("D5",8), rep("D7",8))
sannots=data.frame(snames, groups)
colnames(sannots)=c("SampleName", "Class")

# CEMiTool can't find an appropriate beta for this data (it looks for a value that gives a r2 > 0.8, which doesn't happen), so it must be set manually, and thus, needs to be optimized
hbakeri_coexp_network = cemitool(expr = fpkms_df, annot = sannots, verbose = TRUE, apply_vst = TRUE, filter_pval = 1, set_beta = 10, gsea_max_size = 15000, network_type = "unsigned")

#hbakeri_coexp_network = cemitool(expr = fpkms_df, annot = sannots, verbose = TRUE, apply_vst = TRUE, filter_pval = 1, set_beta = 16, gsea_max_size = 15000, network_type = "signed")

```

```{r}
# Save output of CEMiTool
generate_report(hbakeri_coexp_network, force=TRUE)
write_files(hbakeri_coexp_network, force=TRUE)
save_plots(hbakeri_coexp_network, "all", force=TRUE)
diagnostic_report(hbakeri_coexp_network, force=TRUE)

```

```{r}
# Make a new visulization of the module enrichment

# Unsigned Network
# Import the data if this was run on ARC
enrichment_nes_unsigned <- read.delim("enrichment_nes.tsv")
enrichment_padj_unsigned <- read.delim("enrichment_padj.tsv", row.names=1)

pvals=as.data.frame(t(enrichment_padj))
pvals$dpi=c(10,10,21,21,5,5,7,7)
pvals=pvals[ order(as.numeric(pvals$dpi), decreasing=TRUE), ]

dat=t(enrichment_nes)
colnames(dat)=dat[1,]
dat=dat[-1,]
n=rownames(dat)
dat=as.data.frame(apply(dat, 2, as.numeric))
rownames(dat)=n
dat$group=rownames(dat)
dat$dpi=c(10,10,21,21,5,5,7,7)
dat=dat[ order(as.numeric(dat$dpi), decreasing = TRUE), ]

drawBar <- function(vals, name, fills) {
  barplot(vals, main = name, col = fills, xlim = c(-7,6.5), horiz = TRUE, las=1)
  grid(nx=NULL, ny=NA, lwd=1, lty=1, col="lightgray")
  barplot(vals, main = name, col = fills, xlim = c(-7,6.5), horiz = TRUE, las=1, add=TRUE)
  abline(v=0, lwd=2)
  box(which = "plot")
}

assignColsByPval <- function(vals, pmat) {
  tmp=c()
  i=0
  for (p in pmat[, vals]) {
    i=i+1
    if (!is.na(p) & p < 0.05) { # NES is significant
      if (i%%2 == 1) { # Female
        tmp=append(tmp, "pink", after = length(tmp))
      } else { # Male
        tmp=append(tmp, "lightblue", after = length(tmp))
      }
    } else { # NES NOT significant
      tmp=append(tmp, "lightgray", after = length(tmp))
    }
  }
  
  return(tmp)
}

numpanels=ncol(dat)-2
order=c(seq(0,numpanels))
ws=c(1,rep(2,numpanels))
gs=enrichment_nes_unsigned$pathway
o=c(1,3,4,5,6,7,8,9,10,2,11)
gs=gs[o]

png(file="hbakeri_unsigned_modenrich.png", width=5000, height=1800, res=300)
par(mar=c(2,1,1,0))
layout(mat=matrix(order, nrow=1, ncol=(numpanels+1)), heights = c(), widths = ws)
i=0
for (mod in gs) {
  i=i+1
  cols=assignColsByPval(mod, pvals)
  drawBar(dat[[mod]], mod, cols)
  if (i == 1) {
    axis(2, outer=FALSE, labels=dat$group, at=c(0.7,1.9,3.1,4.3,5.5,6.7,7.9,9.1), line = 0, las=1)
  }
}
dev.off()

# Signed Network
# Import the data if this was run on ARC
enrichment_nes_signed <- read.delim("enrichment_nes.tsv", row.names=1)
enrichment_padj_signed <- read.delim("enrichment_padj.tsv", row.names=1)

pvals_s=as.data.frame(t(enrichment_padj_signed))
pvals_s$dpi=c(10,10,21,21,5,5,7,7)
pvals_s=pvals_s[ order(as.numeric(pvals_s$dpi), decreasing=TRUE), ]

dat_s=as.data.frame(t(enrichment_nes_signed))
dat_s$group=rownames(dat_s)
dat_s$dpi=c(10,10,21,21,5,5,7,7)
dat_s=dat_s[ order(as.numeric(dat_s$dpi), decreasing = TRUE), ]

drawBar2 <- function(vals, name, fills) {
  barplot(vals, main = name, col = fills, xlim = c(-8,8.5), horiz = TRUE, las=1, xaxt='n')
  axis(1, at=c(-8,-6,-4,-2,0,2,4,6,8), labels=FALSE)
  abline(v=c(-6,-4,-2,2,4,6,8), lwd=1, col="lightgray")
  axis(1, at=c(-8,-4,0,4,8), labels=c(-8,-4,0,4,8))
  barplot(vals, main = name, col = fills, xlim = c(-8,8.5), horiz = TRUE, las=1, xaxt='n', add=TRUE)
  abline(v=0, lwd=2)
  box(which = "plot")
}

numpanels_s=ncol(dat_s)-2
order=c(seq(0,numpanels_s))
ws=c(1,rep(2,numpanels_s))
gs=rownames(enrichment_nes_signed)
o=c(1,3,4,5,6,7,8,9,10,2,11)
gs=gs[o]

png(file="hbakeri_signed_modenrich.png", width=5000, height=1800, res=300)
par(mar=c(2,1,1,0))
layout(mat=matrix(order, nrow=1, ncol=(numpanels_s+1)), heights = c(), widths = ws)
i=0
for (mod in gs) {
  i=i+1
  cols=assignColsByPval(mod, pvals_s)
  drawBar2(dat_s[[mod]], mod, cols)
  if (i == 1) {
    axis(2, outer=FALSE, labels=dat_s$group, at=c(0.7,1.9,3.1,4.3,5.5,6.7,7.9,9.1), line = 0, las=1)
  }
}
dev.off()

```
````
