## Supplementary material for "Transcriptional patterns of sexual dimorphism and in host developmental programs in the model parasitic nematode *Heligmosomoides bakeri*": S2 File: hbakeri_infection_DGE.Rmd

````
---
title: "H. bakeri infection transcriptomes DGE"
author: "Stephen Pollo"
date: "03/08/2021"
output: html_document
---

```{r}
# Load libraries
library(DESeq2)

hbakeri_featurecount_output <- read.table("hbakeri_WBPSgenandannots_fc_ss_all_fractionoverlaps_paircheck.txt", row.names=1, comment.char = "#")

# Fix col names
colnames(hbakeri_featurecount_output) = c("Chr","Start","End","Strand","Length", "D10_F10_Li31317_S3", "D10_F11_Li31318_S4", "D10_F13_Li31319_S5", "D10_F1_Li29424_S21","D10_F2_Li29425_S22","D10_F6_Li29426_S23","D10_F7_Li29427_S24", "D10_F8_Li31315_S1", "D10_F9_Li31316_S2", "D10_M1_Li29420_S17","D10_M2_Li29421_S18","D10_M3_Li29422_S19", "D10_M3_Li31320_S6","D10_M6_Li29423_S20", "D10_M6_Li31321_S7","D21_F11_Li29434_S31","D21_F12_Li29435_S32","D21_F5_Li29432_S29","D21_F6_Li29433_S30","D21_M12_Li29430_S27","D21_M13_Li29431_S28","D21_M5_Li29428_S25","D21_M7_Li29429_S26","D5_F2_Li29408_S5","D5_F3_Li29409_S6","D5_F4_Li29410_S7","D5_F5_Li29411_S8","D5_M1_Li29404_S1","D5_M2_Li29405_S2","D5_M4_Li29406_S3","D5_M5_Li29407_S4","D7_F1_Li29416_S13","D7_F4_Li29417_S14","D7_F5_Li29418_S15","D7_F6_Li29419_S16","D7_M1_Li29412_S9","D7_M2_Li29413_S10","D7_M4_Li29414_S11","D7_M7_Li29415_S12")
featurecounts_second_stranded_counts_all=as.data.frame(apply(hbakeri_featurecount_output[-1,-1:-5], 2, as.numeric))
rownames(featurecounts_second_stranded_counts_all)=rownames(hbakeri_featurecount_output[-1,])

```

```{r}
# Functions

# Pull all result combinations for given deseq experiment
getAllResultCombos <- function(deseq_exp) {
  conditions = c("D10_Female", "D10_Male", "D21_Female", "D21_Male", "D5_Female", "D5_Male", "D7_Female", "D7_Male")
  
  tmp = list()
  
  for (i in 1:(length(conditions)-1)) {
    for (j in (i+1):length(conditions)) {
      tmp[[paste(conditions[i], "_vs_", conditions[j], sep="")]] = results(deseq_exp, contrast=c("groups", conditions[i], conditions[j]))
    }
  }
  
  return(tmp)
}

# Find genes common to two lists of interest
getSharedGenes <- function(list1, list2) {
  tmp = intersect(rownames(list1), rownames(list2))
  
  return(tmp)
}

# Find genes common to two vectors
getSharedGenesVector <- function(vector1, vector2) {
  tmp = intersect(vector1, vector2)
  
  return(tmp)
}

# Get genes not found in a given list
# Vector 2 is all genes, vector 1 is genes to remove
getComplementGenes <- function(vector1, vector2) {
  tmp <- vector2[is.na(pmatch(vector2, vector1))]
  
  return(tmp)
}

# Filter a list of DESeq2 results by a specified pvalue cutoff. Returns a subsetted list
filterpadj <- function(DESeq2_results_list, padj_cutoff) {
  tmp=list()
  
  for (i in 1:length(DESeq2_results_list)) {
   tmp[[names(DESeq2_results_list)[i]]] = subset(DESeq2_results_list[[i]], padj < padj_cutoff)
  }
  
  return(tmp)
}

# Filter a list of DESeq2 results by a specified log2FoldChange cutoff. Returns a subsetted list
filterlog2FC <- function(DESeq2_results_list, log2FC_cutoff) {
  tmp=list()
  
  for (i in 1:length(DESeq2_results_list)) {
   tmp[[names(DESeq2_results_list)[i]]] = subset(DESeq2_results_list[[i]], abs(log2FoldChange) > log2FC_cutoff)
  }
  
  return(tmp)
}

# Print out a mini-table of number of genes in comps of interest in relevant list
printNumGenesInCompsOfInterest <- function(DESeq2_results_list, compsofinterestnames) {
  for (i in 1:length(compsofinterestnames)) {
     print(paste(compsofinterestnames[i], nrow(DESeq2_results_list[[compsofinterestnames[i]]]), sep=': '))
  }
}

# Print out mini-table of number of genes common to comps of interest in relevant list
printNumSharedGenesBetweenCompsOfInterest <- function(DESeq2_results_list, compsofinterestnames) {
  for (i in 1:(length(compsofinterestnames)-1)) {
     print(paste("Shared in: ", compsofinterestnames[i], " and ", compsofinterestnames[i+1], ": ",  length(getSharedGenesVector(rownames(DESeq2_results_list[[compsofinterestnames[i]]]), rownames(DESeq2_results_list[[compsofinterestnames[i+1]]]))), sep=''))
  }
}

# Print out a single line of the number of genes in all comps of interest in relevant list
printNumGenesSharedInAllCompsOfInterest <- function(DESeq2_results_list, compsofinterestnames) {
  comps=length(compsofinterestnames)
  num=0
  
  if(comps==2) {
    num=length(getSharedGenesVector(rownames(DESeq2_results_list[[compsofinterestnames[1]]]), rownames(DESeq2_results_list[[compsofinterestnames[2]]])))
  } else {
    share=rownames(DESeq2_results_list[[compsofinterestnames[1]]])
    for (i in 2:comps) {
      share=getSharedGenesVector(rownames(DESeq2_results_list[[compsofinterestnames[i]]]), share)
      
    }
    num=length(share)
  }
  
  print(paste("Shared in all: ", num, sep=""))
}

```

```{r}
# Round the fractional counts to integers
featurecounts_second_stranded_counts_all=round(featurecounts_second_stranded_counts_all)

# Remove the samples that are being excluded
keeps=c("D10_F10_Li31317_S3", "D10_F11_Li31318_S4", "D10_F13_Li31319_S5", "D10_F1_Li29424_S21", "D10_F2_Li29425_S22", "D10_F8_Li31315_S1", "D10_F9_Li31316_S2", "D10_M1_Li29420_S17","D10_M2_Li29421_S18","D10_M3_Li29422_S19", "D10_M6_Li29423_S20", "D21_F11_Li29434_S31", "D21_F12_Li29435_S32", "D21_F5_Li29432_S29", "D21_F6_Li29433_S30", "D21_M12_Li29430_S27", "D21_M13_Li29431_S28", "D21_M5_Li29428_S25", "D21_M7_Li29429_S26", "D5_F2_Li29408_S5", "D5_F3_Li29409_S6", "D5_F4_Li29410_S7", "D5_F5_Li29411_S8", "D5_M1_Li29404_S1", "D5_M2_Li29405_S2", "D5_M4_Li29406_S3", "D5_M5_Li29407_S4", "D7_F1_Li29416_S13", "D7_F4_Li29417_S14", "D7_F5_Li29418_S15", "D7_F6_Li29419_S16", "D7_M1_Li29412_S9", "D7_M2_Li29413_S10", "D7_M4_Li29414_S11", "D7_M7_Li29415_S12")
featurecounts_second_stranded_counts_all=featurecounts_second_stranded_counts_all[keeps]

# Construct DESeq object
groups=c(rep("D10_Female",7), rep("D10_Male",4), rep("D21_Female",4), rep("D21_Male",4), rep("D5_Female",4), rep("D5_Male",4), rep("D7_Female",4), rep("D7_Male",4))

fc_second_stranded=DESeqDataSetFromMatrix(featurecounts_second_stranded_counts_all, DataFrame(groups), ~groups)
de_fc_second_stranded=DESeq(fc_second_stranded)

saveRDS(fc_second_stranded, "hbakeri_deseq2_3-21-22_de_fc_second_stranded.rds")

# Pull results and write to files
res_fc_secondstranded = getAllResultCombos(de_fc_second_stranded)

saveRDS(res_fc_secondstranded, "Hbakeri_deseq2_3-21-22_res_fc_secondstranded.rds")

# for (k in 1:length(res_fc_secondstranded)) {
#   write.csv(res_fc_secondstranded[k], paste("Hbakeri_fc_ss_", names(res_fc_secondstranded)[k], ".csv", sep=""))
# }

```

```{r}
# Investigate effect of padj and log2FC cutoffs on DGE
dge_p0.05_nl2fc=filterpadj(res_fc_secondstranded, 0.05)
dge_p0.01_nl2fc=filterpadj(res_fc_secondstranded, 0.01)
dge_p0.001_nl2fc=filterpadj(res_fc_secondstranded, 0.001)

dge_p0.05_l2fc1=filterlog2FC(dge_p0.05_nl2fc, 1)
dge_p0.01_l2fc1=filterlog2FC(dge_p0.01_nl2fc, 1)
dge_p0.001_l2fc1=filterlog2FC(dge_p0.001_nl2fc, 1)

dge_p0.05_l2fc2=filterlog2FC(dge_p0.05_nl2fc, 2)
dge_p0.01_l2fc2=filterlog2FC(dge_p0.01_nl2fc, 2)
dge_p0.001_l2fc2=filterlog2FC(dge_p0.001_nl2fc, 2)

# Get number of genes in comparisons of interest
compsofinterest=c("D5_Female_vs_D5_Male", "D7_Female_vs_D7_Male", "D10_Female_vs_D10_Male", "D21_Female_vs_D21_Male")

filtered_DEGs=c("dge_p0.05_nl2fc", "dge_p0.01_nl2fc", "dge_p0.001_nl2fc", "dge_p0.05_l2fc1", "dge_p0.01_l2fc1", "dge_p0.001_l2fc1", "dge_p0.05_l2fc2", "dge_p0.01_l2fc2", "dge_p0.001_l2fc2")

for (i in 1:length(filtered_DEGs)) {
  print(filtered_DEGs[i])
  printNumGenesInCompsOfInterest(get(filtered_DEGs[i]), compsofinterest)
  print(" ")
}

# Also print shared genes between comparisons of interest
for (i in 1:length(filtered_DEGs)) {
  print(filtered_DEGs[i])
  printNumSharedGenesBetweenCompsOfInterest(get(filtered_DEGs[i]), compsofinterest)
  printNumGenesSharedInAllCompsOfInterest(get(filtered_DEGs[i]), compsofinterest)
  print(" ")
}

```

```{r}
# Investigate male/female comparison with different grouping and effect of padj and log2FC

# Construct DESeq object, assuming featurecounts file is already rounded
groups2=c(rep("Female",7), rep("Male",4), rep("Female",4), rep("Male",4), rep("Female",4), rep("Male",4), rep("Female",4), rep("Male",4))

fc_second_stranded_mfgroups=DESeqDataSetFromMatrix(featurecounts_second_stranded_counts_all, DataFrame(groups2), ~groups2)
de_fc_second_stranded_mfgroups=DESeq(fc_second_stranded_mfgroups)

# Pull result and write to file
res_fc_secondstranded_mfgroups = results(de_fc_second_stranded_mfgroups, contrast=c("groups2", "Male", "Female"))

write.csv(res_fc_secondstranded_mfgroups, "Hbakeri_fc_ss_mfgroups_male_vs_female.csv")

# Investigate effect of padj and log2FC cutoffs on DGE
dge_mfg_p0.05_nl2fc=subset(res_fc_secondstranded_mfgroups, padj < 0.05)
dge_mfg_p0.01_nl2fc=subset(res_fc_secondstranded_mfgroups, padj < 0.01)
dge_mfg_p0.001_nl2fc=subset(res_fc_secondstranded_mfgroups, padj < 0.001)

dge_mfg_p0.05_l2fc1=subset(dge_mfg_p0.05_nl2fc, abs(log2FoldChange) > 1)
dge_mfg_p0.01_l2fc1=subset(dge_mfg_p0.01_nl2fc, abs(log2FoldChange) > 1)
dge_mfg_p0.001_l2fc1=subset(dge_mfg_p0.001_nl2fc, abs(log2FoldChange) > 1)

dge_mfg_p0.05_l2fc2=subset(dge_mfg_p0.05_nl2fc, abs(log2FoldChange) > 2)
dge_mfg_p0.01_l2fc2=subset(dge_mfg_p0.01_nl2fc, abs(log2FoldChange) > 2)
dge_mfg_p0.001_l2fc2=subset(dge_mfg_p0.001_nl2fc, abs(log2FoldChange) > 2)

# Get number of DE genes
filtered_mfg_DEGs=c("dge_mfg_p0.05_nl2fc", "dge_mfg_p0.01_nl2fc", "dge_mfg_p0.001_nl2fc", "dge_mfg_p0.05_l2fc1", "dge_mfg_p0.01_l2fc1", "dge_mfg_p0.001_l2fc1", "dge_mfg_p0.05_l2fc2", "dge_mfg_p0.01_l2fc2", "dge_mfg_p0.001_l2fc2")

for (i in 1:length(filtered_DEGs)) {
  print(filtered_DEGs[i])
  print(nrow(get(filtered_mfg_DEGs[i])))
  print(" ")
}

```

```{r}
# Find putative housekeeping genes
num_de_fc_nss=list()
for (i in 1:length(res_fc_secondstranded)) {
   num_de_fc_nss[[names(res_fc_secondstranded)[i]]] = subset(res_fc_secondstranded[[i]], padj > 0.05)
}

genes=rownames(featurecounts_second_stranded_counts_all)

for (i in 1:length(num_de_fc_nss)) {
    
    if (i == 1) {
        tmp=getSharedGenesVector(rownames(num_de_fc_nss[[i]]), genes)
    } else {
        tmp=getSharedGenesVector(rownames(num_de_fc_nss[[i]]), tmp)
    }
}
putative_housekeeping_genes=tmp

for (i in 1:length(putative_housekeeping_genes)) {
    print(res_fc_secondstranded$D10_Female_vs_D10_Male[putative_housekeeping_genes[i],])
}

for (i in 1:length(res_fc_secondstranded)) {
    print(res_fc_secondstranded[[i]][putative_housekeeping_genes[7],])
}

# Write putative housekeeping genes to file
write.csv(putative_housekeeping_genes, "putative_housekeeping_gene_names.csv")

```
````
