## Supplementary material for "Transcriptional patterns of sexual dimorphism and in host developmental programs in the model parasitic nematode *Heligmosomoides bakeri*": S2 File: Hbakeri_integrating_DGEandCoexp.Rmd

````
---
title: "Integrating DGE and Co-expression networks"
author: "Stephen Pollo"
date: "19/11/2021"
output: html_document
---

```{r}
# Read in data

Hbakeri_fc_ss_MFcompsANDcoexpnetworks <- readRDS("Hbakeri_fc_ss_MFcompsANDcoexpnetworks.rds") # Table of all DGE results and the coexpression network module for both networks for all transcripts

WBPS_hbakeri_celegans_orthos_mart_export <- read.delim("WBPS_hbakeri_celegans_orthos_mart_export.txt") # All orthologs from WormBase ParaSite

heligmosomoides_bakeri.PRJEB15396.WBPS14.annotations <- read.delim("heligmosomoides_polygyrus.PRJEB15396.WBPS14.annotations.gff3", header=FALSE, comment.char="#")
hbakeri_annot_descriptions=heligmosomoides_bakeri.PRJEB15396.WBPS14.annotations[heligmosomoides_bakeri.PRJEB15396.WBPS14.annotations$V3 == 'mRNA', ]
colnames(hbakeri_annot_descriptions)=c("Contig","Source","Feature_Type","Start","End","Score","Strand","Phase","Attributes")
hbakeri_annot_descriptions$GeneID = substr(hbakeri_annot_descriptions$Attributes, 15, 29)
hbakeri_annot_descriptions$Gene_Description = substr(hbakeri_annot_descriptions$Attributes, 94, nchar(hbakeri_annot_descriptions$Attributes))
hbakeri_annots_nondup = hbakeri_annot_descriptions[!duplicated(hbakeri_annot_descriptions$GeneID),]

```

```{r}
# For genes up-regulated in males
allm=subset(Hbakeri_fc_ss_MFcompsANDcoexpnetworks, D5_Female_vs_D5_Male.padj < 0.05 & D7_Female_vs_D7_Male.padj < 0.05 & D10_Female_vs_D10_Male.padj < 0.05 & D21_Female_vs_D21_Male.padj < 0.05)
upm=subset(allm, D5_Female_vs_D5_Male.log2FoldChange < 0 & D7_Female_vs_D7_Male.log2FoldChange < 0 & D10_Female_vs_D10_Male.log2FoldChange < 0 & D21_Female_vs_D21_Male.log2FoldChange < 0)

data.frame(table(upm$`Unsigned Network modules`))
data.frame(table(upm$`Signed Network Modules`))

upm_uns=upm[is.na(upm$`Unsigned Network modules`)==FALSE & (upm$`Unsigned Network modules`=='M3' | upm$`Unsigned Network modules`=='M5'),]
upm_b=upm_uns[is.na(upm_uns$`Signed Network Modules`)==FALSE & (upm_uns$`Signed Network Modules`=='M3' | upm_uns$`Signed Network Modules`=='M4'),]

upm_b$BiggestLog2FC=pmax(abs(upm_b$D5_Female_vs_D5_Male.log2FoldChange), abs(upm_b$D7_Female_vs_D7_Male.log2FoldChange), abs(upm_b$D10_Female_vs_D10_Male.log2FoldChange), abs(upm_b$D21_Female_vs_D21_Male.log2FoldChange))

#write.csv(upm_b, "Hbakeri_p0.05log2fcl0inall4comps_malemodules.csv")

```

```{r}
# Check which genes have homologs in C. elegans
upm_b$Gene_ID=substr(upm_b$...1, 12, 26)
orthopairs=WBPS_hbakeri_celegans_orthos_mart_export[,-1]

upm_orths=merge(upm_b, orthopairs, by.x = "Gene_ID", by.y = "Gene.stable.ID", all.x = TRUE, all.y = FALSE)
upm_noh=upm_orths[is.na(upm_orths$Homology.type),]
upm_m2m=subset(upm_orths, Homology.type=="ortholog_many2many")
upm_o2m=subset(upm_orths, Homology.type=="ortholog_one2many")
upm_o2o=subset(upm_orths, Homology.type=="ortholog_one2one")

# Coarsely classify genes by presence of a homolog in C. elegans or not
uniq_upm=data.frame(table(upm_b$Gene_ID))
uniq_upm_noh=data.frame(table(upm_noh$Gene_ID))
uniq_upm_m2m=data.frame(table(upm_m2m$Gene_ID))
uniq_upm_o2m=data.frame(table(upm_o2m$Gene_ID))
uniq_upm_o2o=data.frame(table(upm_o2o$Gene_ID))

# Get lists of genes with their C. elegans homolog for looking up
uniq_upm_o2o_withdesc=upm_o2o[!duplicated(upm_o2o$Gene_ID),]
uniq_upm_o2m_withdesc=upm_o2m[!duplicated(upm_o2m$Gene_ID),]
uniq_upm_m2m_withdesc=upm_m2m[!duplicated(upm_m2m$Gene_ID),]
# write.csv(uniq_upm_o2o_withdesc, "Hbakeri_upm_wcelehomo2o.csv")
# write.csv(uniq_upm_o2m_withdesc, "Hbakeri_upm_wcelehomo2m.csv")
# write.csv(uniq_upm_m2m_withdesc, "Hbakeri_upm_wcelehomm2m.csv")

# Genes with no homologs have to rely on H. bakeri annotation
uniq_upm_noh_withdesc=merge(uniq_upm_noh, hbakeri_annots_nondup, by.x = "Var1", by.y = "GeneID", all.x = TRUE, all.y = FALSE)

# How many hypothetical proteins?
nrow(subset(uniq_upm_noh_withdesc, Gene_Description == ""))

# Get genes with annotation info
uniq_upm_noh_annotated=subset(uniq_upm_noh_withdesc, Gene_Description != "")
#write.csv(uniq_upm_noh_annotated, "Hbakeri_upm_noh_annotated.csv")

```

```{r}
# For genes up-regulated in females
allm=subset(Hbakeri_fc_ss_MFcompsANDcoexpnetworks, D5_Female_vs_D5_Male.padj < 0.05 & D7_Female_vs_D7_Male.padj < 0.05 & D10_Female_vs_D10_Male.padj < 0.05 & D21_Female_vs_D21_Male.padj < 0.05)
upf=subset(allm, D5_Female_vs_D5_Male.log2FoldChange > 0 & D7_Female_vs_D7_Male.log2FoldChange > 0 & D10_Female_vs_D10_Male.log2FoldChange > 0 & D21_Female_vs_D21_Male.log2FoldChange > 0)

data.frame(table(upf$`Unsigned Network modules`))
data.frame(table(upf$`Signed Network Modules`))

upf_uns=upf[is.na(upf$`Unsigned Network modules`)==FALSE & (upf$`Unsigned Network modules`=='M1' | upf$`Unsigned Network modules`=='M9'),]
upf_b=upf_uns[is.na(upf_uns$`Signed Network Modules`)==FALSE & (upf_uns$`Signed Network Modules`=='M1' | upf_uns$`Signed Network Modules`=='M6' | upf_uns$`Signed Network Modules`=='M9'),]

upf_b$BiggestLog2FC=pmax(abs(upf_b$D5_Female_vs_D5_Male.log2FoldChange), abs(upf_b$D7_Female_vs_D7_Male.log2FoldChange), abs(upf_b$D10_Female_vs_D10_Male.log2FoldChange), abs(upf_b$D21_Female_vs_D21_Male.log2FoldChange))

#write.csv(upf_b, "Hbakeri_p0.05log2fcg0inall4comps_femalemodules.csv")

# Unsigned M3 is a male-enriched module. What is the deal with these?
weird_m3s=upf[is.na(upf$`Unsigned Network modules`)==FALSE & (upf$`Unsigned Network modules`=='M3'),]

# Check which genes have homologs in C. elegans
upf_b$Gene_ID=substr(upf_b$...1, 12, 26)
orthopairs=WBPS_hbakeri_celegans_orthos_mart_export[,-1]

upf_orths=merge(upf_b, orthopairs, by.x = "Gene_ID", by.y = "Gene.stable.ID", all.x = TRUE, all.y = FALSE)
upf_noh=upf_orths[is.na(upf_orths$Homology.type),]
upf_m2m=subset(upf_orths, Homology.type=="ortholog_many2many")
upf_o2m=subset(upf_orths, Homology.type=="ortholog_one2many")
upf_o2o=subset(upf_orths, Homology.type=="ortholog_one2one")

# Coarsely classify genes by presence of a homolog in C. elegans or not
uniq_upf=data.frame(table(upf_b$Gene_ID))
uniq_upf_noh=data.frame(table(upf_noh$Gene_ID))
uniq_upf_m2m=data.frame(table(upf_m2m$Gene_ID))
uniq_upf_o2m=data.frame(table(upf_o2m$Gene_ID))
uniq_upf_o2o=data.frame(table(upf_o2o$Gene_ID))

# Get lists of genes with their C. elegans homolog for looking up
uniq_upf_o2o_withdesc=upf_o2o[!duplicated(upf_o2o$Gene_ID),]
uniq_upf_o2m_withdesc=upf_o2m[!duplicated(upf_o2m$Gene_ID),]
uniq_upf_m2m_withdesc=upf_m2m[!duplicated(upf_m2m$Gene_ID),]
# write.csv(uniq_upf_o2o_withdesc, "Hbakeri_upf_wcelehomo2o.csv")
# write.csv(uniq_upf_o2m_withdesc, "Hbakeri_upf_wcelehomo2m.csv")
# write.csv(uniq_upf_m2m_withdesc, "Hbakeri_upf_wcelehomm2m.csv")

# Genes with no homologs have to rely on H. bakeri annotation
uniq_upf_noh_withdesc=merge(uniq_upf_noh, hbakeri_annots_nondup, by.x = "Var1", by.y = "GeneID", all.x = TRUE, all.y = FALSE)

# How many hypothetical proteins?
nrow(subset(uniq_upf_noh_withdesc, Gene_Description == ""))

# Get genes with annotation info
uniq_upf_noh_annotated=subset(uniq_upf_noh_withdesc, Gene_Description != "")

```

```{r}
# Developmental analysis

library(DESeq2)

# Import the DESeq2 results
de_fc_second_stranded = readRDS("hbakeri_deseq2_3-21-22_de_fc_second_stranded.rds")
res_fc_secondstranded = readRDS("Hbakeri_deseq2_3-21-22_res_fc_secondstranded.rds")

# Import the signed and unsigned module assignments
signed_modules <- read.delim("signed/module.tsv", row.names=1)
unsigned_modules <- read.delim("unsigned/module.tsv", row.names=1)
genemods=merge(unsigned_modules, signed_modules, by=0, suffixes=c(".unsigned", ".signed"))
rownames(genemods)=genemods[,1]
genemods=genemods[,-1]

# Get expression estimates for every gene in all samples
vst_exprs=vst(de_fc_second_stranded, blind = FALSE)
vst_exprs_mat=assay(vst_exprs)
vst_exprs_df=as.data.frame(vst_exprs_mat)
saveRDS(vst_exprs_df, "vst_exprs_df_hbakeri.rds")

# Construct the sets of pairwise comparisons to use
tmpm1=merge(as.data.frame(res_fc_secondstranded$D5_Male_vs_D7_Male), as.data.frame(res_fc_secondstranded$D10_Male_vs_D7_Male), by=0, suffixes=c(".D5MvsD7M", ".D10MvsD7M"))
tmpm2=merge(as.data.frame(res_fc_secondstranded$D10_Male_vs_D21_Male), as.data.frame(res_fc_secondstranded$D21_Male_vs_D5_Male), by=0, suffixes=c(".D10MvsD21M", ".D21MvsD5M"))
rownames(tmpm1)=tmpm1[,1]
tmpm1=tmpm1[,-1]
rownames(tmpm2)=tmpm2[,1]
tmpm2=tmpm2[,-1]
tmpm3=merge(tmpm1, tmpm2, by=0)
rownames(tmpm3)=tmpm3[,1]
tmpm3=tmpm3[,-1]
devpcsmale=merge(tmpm3, genemods, by=0, all.x=TRUE)
rownames(devpcsmale)=devpcsmale[,1]
devpcsmale=devpcsmale[,-1]

tmpf1=merge(as.data.frame(res_fc_secondstranded$D5_Female_vs_D7_Female), as.data.frame(res_fc_secondstranded$D10_Female_vs_D7_Female), by=0, suffixes=c(".D5FvsD7F", ".D10FvsD7F"))
tmpf2=merge(as.data.frame(res_fc_secondstranded$D10_Female_vs_D21_Female), as.data.frame(res_fc_secondstranded$D21_Female_vs_D5_Female), by=0, suffixes=c(".D10FvsD21F", ".D21FvsD5F"))
rownames(tmpf1)=tmpf1[,1]
tmpf1=tmpf1[,-1]
rownames(tmpf2)=tmpf2[,1]
tmpf2=tmpf2[,-1]
tmpf3=merge(tmpf1, tmpf2, by=0)
rownames(tmpf3)=tmpf3[,1]
tmpf3=tmpf3[,-1]
devpcsfemale=merge(tmpf3, genemods, by=0, all.x=TRUE)
rownames(devpcsfemale)=devpcsfemale[,1]
devpcsfemale=devpcsfemale[,-1]

rm(tmpm1)
rm(tmpm2)
rm(tmpm3)
rm(tmpf1)
rm(tmpf2)
rm(tmpf3)

# Function for doing the comparisons. Need to compare all replicates against all replicates for each group
# rel is relationship between group1 and group2: gr group1 > group2, lt group1 < group2, function returns 0 otherwise
compExpVals <- function(df, group1, group2, rel) {
  i1=switch(group1, "d10f" = 1:7, "d10m" = 8:11, "d21f" = 12:15, "d21m" = 16:19, "d5f" = 20:23, "d5m" = 24:27, "d7f" = 28:31, "d7m" = 32:35)
  j1=switch(group2, "d10f" = 1:7, "d10m" = 8:11, "d21f" = 12:15, "d21m" = 16:19, "d5f" = 20:23, "d5m" = 24:27, "d7f" = 28:31, "d7m" = 32:35)
  
  tmp=df
  
  for (i in i1) { # iterate through the first group
    for (j in j1) { #iterate through the second group
      if(rel == "gr") {
        tmp=tmp[tmp[,i] > tmp[,j],]
      }
      else if(rel == "lt") {
        tmp=tmp[tmp[,i] < tmp[,j],]
      }
      else {return(0)}
    }
  }
  
  return(tmp)
}

# Get strictly increasing genes
tmp=compExpVals(vst_exprs_df, "d7m", "d5m", "gr")
tmp2=compExpVals(tmp, "d10m", "d7m", "gr")
strictlyincreasingmales=compExpVals(tmp2, "d21m", "d10m", "gr")

tmp=compExpVals(vst_exprs_df, "d7f", "d5f", "gr")
tmp2=compExpVals(tmp, "d10f", "d7f", "gr")
strictlyincreasingfemales=compExpVals(tmp2, "d21f", "d10f", "gr")

# Get strictly decreasing genes
tmp=compExpVals(vst_exprs_df, "d7m", "d5m", "lt")
tmp2=compExpVals(tmp, "d10m", "d7m", "lt")
strictlydecreasingmales=compExpVals(tmp2, "d21m", "d10m", "lt")

tmp=compExpVals(vst_exprs_df, "d7f", "d5f", "lt")
tmp2=compExpVals(tmp, "d10f", "d7f", "lt")
strictlydecreasingfemales=compExpVals(tmp2, "d21f", "d10f", "lt")

# Get increased lumen over tissue
tmp=compExpVals(vst_exprs_df, "d10m", "d5m", "gr")
tmp2=compExpVals(vst_exprs_df, "d10m", "d7m", "gr")
tmp3=compExpVals(vst_exprs_df, "d21m", "d5m", "gr")
tmp4=compExpVals(vst_exprs_df, "d21m", "d7m", "gr")
tmp5=intersect(rownames(tmp), rownames(tmp2))
tmp6=intersect(rownames(tmp3), rownames(tmp4))
tmp7=intersect(tmp5, tmp6)
ilvstmales=vst_exprs_df[tmp7,]

tmp=compExpVals(vst_exprs_df, "d10f", "d5f", "gr")
tmp2=compExpVals(vst_exprs_df, "d10f", "d7f", "gr")
tmp3=compExpVals(vst_exprs_df, "d21f", "d5f", "gr")
tmp4=compExpVals(vst_exprs_df, "d21f", "d7f", "gr")
tmp5=intersect(rownames(tmp), rownames(tmp2))
tmp6=intersect(rownames(tmp3), rownames(tmp4))
tmp7=intersect(tmp5, tmp6)
ilvstfemales=vst_exprs_df[tmp7,]

# Get decreased lumen over tissue
tmp=compExpVals(vst_exprs_df, "d10m", "d5m", "lt")
tmp2=compExpVals(vst_exprs_df, "d10m", "d7m", "lt")
tmp3=compExpVals(vst_exprs_df, "d21m", "d5m", "lt")
tmp4=compExpVals(vst_exprs_df, "d21m", "d7m", "lt")
tmp5=intersect(rownames(tmp), rownames(tmp2))
tmp6=intersect(rownames(tmp3), rownames(tmp4))
tmp7=intersect(tmp5, tmp6)
dlvstmales=vst_exprs_df[tmp7,]

tmp=compExpVals(vst_exprs_df, "d10f", "d5f", "lt")
tmp2=compExpVals(vst_exprs_df, "d10f", "d7f", "lt")
tmp3=compExpVals(vst_exprs_df, "d21f", "d5f", "lt")
tmp4=compExpVals(vst_exprs_df, "d21f", "d7f", "lt")
tmp5=intersect(rownames(tmp), rownames(tmp2))
tmp6=intersect(rownames(tmp3), rownames(tmp4))
tmp7=intersect(tmp5, tmp6)
dlvstfemales=vst_exprs_df[tmp7,]

rm(tmp)
rm(tmp2)
rm(tmp3)
rm(tmp4)
rm(tmp5)
rm(tmp6)
rm(tmp7)

# See how many genes with expression patterns of interest are statistically significant in relevant pairwise comparisons
# Male
sim_sig=devpcsmale[rownames(strictlyincreasingmales),]
sdm_sig=devpcsmale[rownames(strictlydecreasingmales),]
ilvstm_sig=devpcsmale[rownames(ilvstmales),]
dlvstm_sig=devpcsmale[rownames(dlvstmales),]
# Female
sif_sig=devpcsfemale[rownames(strictlyincreasingfemales),]
sdf_sig=devpcsfemale[rownames(strictlydecreasingfemales),]
ilvstf_sig=devpcsfemale[rownames(ilvstfemales),]
dlvstf_sig=devpcsfemale[rownames(dlvstfemales),]

# Filter using coexpression modules
# Function to filter based on coexpression modules
filterCoexpMods <- function(x, col_name, vals_list) {
  coli=grep(col_name, colnames(x))
  tmp=x[is.na(x[coli])==FALSE,]
  for (val in vals_list) {
    if(val == vals_list[1]) {
      tmp2=tmp[tmp[,coli] == val,]
    } else {
      tmp3=tmp[tmp[,coli] == val,]
      tmp2=rbind(tmp2, tmp3)
    }
  }
  return(tmp2)
}

# Male
sim_sig_coexpuns=filterCoexpMods(sim_sig, 'modules.unsigned', c('M4', 'M3', 'M5'))
sim_sig_coexpboth=filterCoexpMods(sim_sig_coexpuns, 'modules.signed', c('M5', 'M3', 'M4'))
sdm_sig_coexpuns=filterCoexpMods(sdm_sig, 'modules.unsigned', c('M2', 'M6', 'M3'))
sdm_sig_coexpboth=filterCoexpMods(sdm_sig_coexpuns, 'modules.signed', c('M2', 'M6', 'M7'))

ilvstm_sig_coexpuns=filterCoexpMods(ilvstm_sig, 'modules.unsigned', c('M4', 'M3', 'M5'))
ilvstm_sig_coexpboth=filterCoexpMods(ilvstm_sig_coexpuns, 'modules.signed', c('M5', 'M3', 'M4'))
dlvstm_sig_coexpuns=filterCoexpMods(dlvstm_sig, 'modules.unsigned', c('M2', 'M3', 'M5', 'M6'))
dlvstm_sig_coexpboth=filterCoexpMods(dlvstm_sig_coexpuns, 'modules.signed', c('M2', 'M6', 'M7', 'M8'))

# Female
sif_sig_coexpuns=filterCoexpMods(sif_sig, 'modules.unsigned', c('M1', 'M3', 'M4', 'M7'))
sif_sig_coexpboth=filterCoexpMods(sif_sig_coexpuns, 'modules.signed', c('M1', 'M3', 'M5'))
sdf_sig_coexpuns=filterCoexpMods(sdf_sig, 'modules.unsigned', c('M2', 'M3', 'M5'))
sdf_sig_coexpboth=filterCoexpMods(sdf_sig_coexpuns, 'modules.signed', c('M2', 'M3', 'M4', 'M7'))

ilvstf_sig_coexpuns=filterCoexpMods(ilvstf_sig, 'modules.unsigned', c('M1', 'M3', 'M4', 'M5', 'M7'))
ilvstf_sig_coexpboth=filterCoexpMods(ilvstf_sig_coexpuns, 'modules.signed', c('M1', 'M3', 'M5', 'M6', 'M9'))
dlvstf_sig_coexpuns=filterCoexpMods(dlvstf_sig, 'modules.unsigned', c('M2', 'M3', 'M5', 'M6', 'M8', 'M10'))
dlvstf_sig_coexpboth=filterCoexpMods(dlvstf_sig_coexpuns, 'modules.signed', c('M2', 'M3', 'M4', 'M7', 'M8', 'M10'))

# Function to extract genes that are statistically significantly different in all relevant pairwise comparisons
getGenesSSAllComps <- function(x, cutoff) {
  colis=grep("padj", colnames(x))
  tmp=x
  for(i in colis) {
    tmp=tmp[is.na(tmp[,i]) == FALSE & tmp[,i] < cutoff,]
  }
  return(tmp)
}

# Function to extract genes that are not statistically signficantly different in any relevant pairwise comparison
# An uncomputed adjusted pvalue (padj = NA) is included as not statistically significantly different
getGenesNSSAnyComp <- function(x, cutoff){
  colis=grep("padj", colnames(x))
  tmp=x
  for(i in colis) {
    tmp=tmp[is.na(tmp[,i]) == TRUE | tmp[,i] >= cutoff,]
  }
  return(tmp)
}

# Narrow down lists for manual ortholog checking
sim_coexpboth_sigall=getGenesSSAllComps(sim_sig_coexpboth, 0.05)
sdm_coexpboth_sigall=getGenesSSAllComps(sdm_sig_coexpboth, 0.05)
ilvstm_coexpboth_sigall=getGenesSSAllComps(ilvstm_sig_coexpboth, 0.05)
dlvstm_coexpboth_sigall=getGenesSSAllComps(dlvstm_sig_coexpboth, 0.05)

sif_coexpboth_sigall=getGenesSSAllComps(sif_sig_coexpboth, 0.05)
sdf_coexpboth_sigall=getGenesSSAllComps(sdf_sig_coexpboth, 0.05)
ilvstf_coexpboth_sigall=getGenesSSAllComps(ilvstf_sig_coexpboth, 0.05)
dlvstf_coexpboth_sigall=getGenesSSAllComps(dlvstf_sig_coexpboth, 0.05)

# Check which genes have homologs in C. elegans
orthopairs=WBPS_hbakeri_celegans_orthos_mart_export[,-1]

# Function to get genes with an ortholog in C. elegans
# Get one2one orthos with o2o, one2many orthos with o2m, many2many orthos with m2m, no orthos with none. Returns 0 for anything else
getCeleOrthos <- function(x, ortho_type){
  tmp=x
  tmp$Gene_ID=substr(rownames(tmp), 12, 26)
  tmp2=merge(tmp, orthopairs, by.x = "Gene_ID", by.y = "Gene.stable.ID", all.x = TRUE, all.y = FALSE)
  if(ortho_type == "o2o") {
    return(tmp2[tmp2$Homology.type == "ortholog_one2one" & is.na(tmp2$Homology.type) == FALSE,])
  } else if(ortho_type == "o2m") {
    return(tmp2[tmp2$Homology.type == "ortholog_one2many" & is.na(tmp2$Homology.type) == FALSE,])
  } else if(ortho_type == "m2m") {
    return(tmp2[tmp2$Homology.type == "ortholog_many2many" & is.na(tmp2$Homology.type) == FALSE,])
  } else if(ortho_type == "none") {
    return(tmp2[is.na(tmp2$Homology.type) == TRUE,])
  } else {
    return(0)
  }
}

tmp=getCeleOrthos(sim_coexpboth_sigall, "o2o")
sim_coexpboth_sigall_o2oorthos=tmp[!duplicated(tmp$Gene_ID),]
tmp=getCeleOrthos(sdf_coexpboth_sigall, "o2o")
sdm_coexpboth_sigall_o2oorthos=tmp[!duplicated(tmp$Gene_ID),]
tmp=getCeleOrthos(ilvstm_coexpboth_sigall, "o2o")
ilvstm_coexpboth_sigall_o2oorthos=tmp[!duplicated(tmp$Gene_ID),]
tmp=getCeleOrthos(dlvstm_coexpboth_sigall, "o2o")
dlvstm_coexpboth_sigall_o2oorthos=tmp[!duplicated(tmp$Gene_ID),]

tmp=getCeleOrthos(sif_coexpboth_sigall, "o2o")
sif_coexpboth_sigall_o2oorthos=tmp[!duplicated(tmp$Gene_ID),]
tmp=getCeleOrthos(sdf_coexpboth_sigall, "o2o")
sdf_coexpboth_sigall_o2oorthos=tmp[!duplicated(tmp$Gene_ID),]
tmp=getCeleOrthos(ilvstf_coexpboth_sigall, "o2o")
ilvstf_coexpboth_sigall_o2oorthos=tmp[!duplicated(tmp$Gene_ID),]
tmp=getCeleOrthos(dlvstf_coexpboth_sigall, "o2o")
dlvstf_coexpboth_sigall_o2oorthos=tmp[!duplicated(tmp$Gene_ID),]

rm(tmp)

# Define full lists of genes in each category (will include all the orthos from above) incorporating deseq2 stats
# For strictly increasing or strictly decreasing genes, genes can be statistically significantly different in any of the pairwise comparisons being considered (ie. remove genes that are not stat. sig. diff. in any pairwise comp)
sim_coexpboth_sigalo=sim_sig_coexpboth[setdiff(rownames(sim_sig_coexpboth), rownames(getGenesNSSAnyComp(sim_sig_coexpboth, 0.05))),]
sdm_coexpboth_sigalo=sdm_sig_coexpboth[setdiff(rownames(sdm_sig_coexpboth), rownames(getGenesNSSAnyComp(sdm_sig_coexpboth, 0.05))),]
sif_coexpboth_sigalo=sif_sig_coexpboth[setdiff(rownames(sif_sig_coexpboth), rownames(getGenesNSSAnyComp(sif_sig_coexpboth, 0.05))),]
sdf_coexpboth_sigalo=sdf_sig_coexpboth[setdiff(rownames(sdf_sig_coexpboth), rownames(getGenesNSSAnyComp(sdf_sig_coexpboth, 0.05))),]

# For genes increasing or decreasing in the lumen relative to the tissue, genes must be statistically significantly different in at least one lumen vs tissue comparison (but don't need to be in tissue vs tissue or lumen vs lumen comarisons)
# Function to extract genes that are statistically significantly different in specified pairwise comparisons
getGenesSSCertainComps <- function(x, cutoff, padj_colnums) {
  tmp=x
  removes=tmp
  for(i in padj_colnums) {
    throws=removes[is.na(removes[,i]) == TRUE | removes[,i] >= cutoff,]
    removes=throws
  }
  tmp2=tmp[setdiff(rownames(tmp), rownames(removes)),]
  return(tmp2)
}

ilvstm_coexpboth_sigrel=getGenesSSCertainComps(ilvstm_sig_coexpboth, 0.05, c(12, 24))
dlvstm_coexpboth_sigrel=getGenesSSCertainComps(dlvstm_sig_coexpboth, 0.05, c(12, 24))
ilvstf_coexpboth_sigrel=getGenesSSCertainComps(ilvstf_sig_coexpboth, 0.05, c(12, 24))
dlvstf_coexpboth_sigrel=getGenesSSCertainComps(dlvstf_sig_coexpboth, 0.05, c(12, 24))

# Get shared tissue-dwelling transcripts
tdshared=intersect(rownames(dlvstm_coexpboth_sigrel), rownames(dlvstf_coexpboth_sigrel))
tds_transcripts=dlvstm_coexpboth_sigrel[tdshared,]

# Get shared lumen-dwelling transcripts
ldshared=intersect(rownames(ilvstm_coexpboth_sigrel), rownames(ilvstf_coexpboth_sigrel))
lds_transcripts=ilvstm_coexpboth_sigrel[ldshared,]

# Maybe for use later
table(sim_sig$padj.D5MvsD7M >= 0.05)
nrow(subset(sim_sig, padj.D5MvsD7M >= 0.05 & padj.D10MvsD7M >= 0.05 & padj.D10MvsD21M >= 0.05 & padj.D21MvsD5M >= 0.05))
nrow(subset(sim_sig, padj.D5MvsD7M < 0.05 & padj.D10MvsD7M < 0.05 & padj.D10MvsD21M < 0.05 & padj.D21MvsD5M < 0.05))
test=subset(devpcsmale, padj.D5MvsD7M < 0.05 & padj.D10MvsD7M < 0.05 & padj.D10MvsD21M < 0.05 & padj.D21MvsD5M < 0.05)

# Function to look up a given gene locus tag in the 4 final lists
lookupGeneInWorms <- function(lt) {
  rn=paste("transcript:", lt, "-mRNA-1", sep="")
  
  print("Female dev pairwise comps")
  print(devpcsfemale[rownames(devpcsfemale) == rn,])
  print("Male dev pairwise comps")
  print(devpcsmale[rownames(devpcsmale) == rn,])
  
  print("Tissue females:")
  tmp=dlvstf_coexpboth_sigrel[rownames(dlvstf_coexpboth_sigrel) == rn,]
  if (nrow(tmp) == 0) {print("  Gene not special to me")} else {print(tmp)}
  print("Lumen females:")
  tmp=ilvstf_coexpboth_sigrel[rownames(ilvstf_coexpboth_sigrel) == rn,]
  if (nrow(tmp) == 0) {print("  Gene not special to me")} else {print(tmp)}
  print("Tissue males:")
  tmp=dlvstm_coexpboth_sigrel[rownames(dlvstm_coexpboth_sigrel) == rn,]
  if (nrow(tmp) == 0) {print("  Gene not special to me")} else {print(tmp)}
  print("Lumen males:")
  tmp=ilvstm_coexpboth_sigrel[rownames(ilvstm_coexpboth_sigrel) == rn,]
  if (nrow(tmp) == 0) {print("  Gene not special to me")} else {print(tmp)}
}

```

```{r}
# Functional enrichment analysis
library(gprofiler2)

# M/F
upm_fe = gost(query=substr(upm_b$...1, 12, 100), organism="hepolyprjeb15396")
upf_fe = gost(query=substr(upf_b$...1, 12, 100), organism="hepolyprjeb15396")

# Dev series
ilvstf_fe = gost(query=substr(rownames(ilvstf_coexpboth_sigrel), 12, 100), organism="hepolyprjeb15396")
dlvstf_fe = gost(query=substr(rownames(dlvstf_coexpboth_sigrel), 12, 100), organism="hepolyprjeb15396")
ilvstm_fe = gost(query=substr(rownames(ilvstm_coexpboth_sigrel), 12, 100), organism="hepolyprjeb15396")
dlvstm_fe = gost(query=substr(rownames(dlvstm_coexpboth_sigrel), 12, 100), organism="hepolyprjeb15396")

# Common devs
tdshare_fe=gost(query=substr(tdshared, 12, 100), organism="hepolyprjeb15396")
ldshare_fe=gost(query=substr(ldshared, 12, 100), organism="hepolyprjeb15396")

# Can look through results with 
ilvstf_fe$result
ilvstf_fe$result$term_name

# Can generate interactive plots with
p=gostplot(ilvstf_fe, capped = FALSE, interactive = TRUE)

```

```{r}
# Defining sex-specific genes
# Male
# Want genes to have at least 10-fold higher expression in all male samples than female samples - 10-fold = log2FC of -3.321928
ms=subset(upm, D5_Female_vs_D5_Male.log2FoldChange <= -3.321928 & D7_Female_vs_D7_Male.log2FoldChange <= -3.321928 & D10_Female_vs_D10_Male.log2FoldChange <= -3.321928 & D21_Female_vs_D21_Male.log2FoldChange <= -3.321928)
ms$Gene_ID=substr(ms$...1, 12, 26)
ms_withdescs=merge(ms, hbakeri_annots_nondup, by.x="Gene_ID", by.y="GeneID", all.x=TRUE)

# Female
# Want genes to have at least 10-fold higher expression in all female samples than male samples - 10-fold - log2FC of 3.321928
fs=subset(upf, D5_Female_vs_D5_Male.log2FoldChange >= 3.321928 & D7_Female_vs_D7_Male.log2FoldChange >= 3.321928 & D10_Female_vs_D10_Male.log2FoldChange >= 3.321928 & D21_Female_vs_D21_Male.log2FoldChange >= 3.321928)
fs$Gene_ID=substr(fs$...1, 12, 26)
fs_withdescs=merge(fs, hbakeri_annots_nondup, by.x="Gene_ID", by.y="GeneID", all.x=TRUE)

```

```{r}
# Immunomodulatory gene expression

#Do all immunomod genes as lines of avg at each timepoint
library(viridis)

immlineplot <- function(l, mt) {
  par(mar=c(5,4,2,15))
  favg=data.frame()
  mavg=data.frame()
  qs=vector()
  cols=viridis(nrow(l), option="D", alpha = 0.5)
  for(i in 1:nrow(l)){
    query=paste("transcript:", l[i,2], sep="")
    if(grepl('-', query, fixed=TRUE)) {} else {query=paste(query, "-mRNA-1", sep="")}
    qs=append(qs, substr(query, 12, nchar(query)))
    tmp=vst_exprs_df[rownames(vst_exprs_df) == query,]
    tmp2=tmp[,c(20:23,28:31,1:7,12:15)] # Females
    tmp3=tmp[,c(24:27,32:35,8:11,16:19)] #Males
    tmp4=c(mean(unlist(tmp2[1:4])), mean(unlist(tmp2[5:8])), mean(unlist(tmp2[9:15])), mean(unlist(tmp2[16:19]))) # Female avgs
    tmp5=c(mean(unlist(tmp3[1:4])), mean(unlist(tmp3[5:8])), mean(unlist(tmp3[9:12])), mean(unlist(tmp3[13:16]))) # Male avgs
    favg=rbind(favg, tmp4)
    mavg=rbind(mavg, tmp5)
  }
  plot(2, favg[1,1], ylim=c(0,25), xlim=c(0,32), xaxt='n', ylab="VST normalized expression averaged across replicates", xlab=NA,  main=mt, type="l")
  abline(v=c(0,4,8,12,16,20,24,28), col="grey")
  axis(1, at=c(0,4,8,12,16,20,24,28), labels=FALSE)
  text(c(3,7,11,15,19,23,27,31), labels=c("D5F", "D7F", "D10F", "D21F", "D5M", "D7M", "D10M", "D21M"), srt=35, cex=1.2, y=par("usr")[3]-1.5, xpd=NA, adj=0.99)
  for(i in 1:nrow(l)){
    lines(seq(2,14,by=4), favg[i,], type="b", col=cols[i])
    lines(seq(18,30,by=4), mavg[i,], type="b", pch=15, col=cols[i])
  }
  legend("topright", inset = c(-0.20,0), legend=qs, col=cols, lty=1, cex=0.5, xpd=TRUE)
}

Identified_classes_of_immunomods=readRDS("Identified_classes_of_immunomods.rds") # Table of gene types and the H bakeri transcripts that match

vals=Identified_classes_of_immunomods[106:119,]
sushi=Identified_classes_of_immunomods[12:105,]
apy=Identified_classes_of_immunomods[122:134,]
aritgm=Identified_classes_of_immunomods[c(1,6,11),]
ast=Identified_classes_of_immunomods[135:182,]
serpcyst=Identified_classes_of_immunomods[c(183:189,120:121),]

immlineplot(serpcyst, "Serpins and Cystatins")
immlineplot(aritgm, "HpARI and HpTGMs")
immlineplot(ast,"Astacin proteases")
immlineplot(apy,"Apyrases")
immlineplot(sushi,"Sushi domains")
immlineplot(vals,"VALs")

png(file="hbakeri_vals_expr.png", width=4500, height=3500, res=300)
immlineplot(vals,"VALs")
dev.off()

png(file="hbakeri_sushi_expr.png", width=4500, height=3500, res=300)
immlineplot(sushi,"Sushi domains")
dev.off()

png(file="hbakeri_apyrases_expr.png", width=4500, height=3500, res=300)
immlineplot(apy,"Apyrases")
dev.off()

png(file="hbakeri_astacinproteases_expr.png", width=4500, height=3500, res=300)
immlineplot(ast,"Astacin proteases")
dev.off()

png(file="hbakeri_serpinscystatins_expr.png", width=4500, height=3500, res=300)
immlineplot(serpcyst, "Serpins and Cystatins")
dev.off()

png(file="hbakeri_aritgm_expr.png", width=4500, height=3500, res=300)
immlineplot(aritgm, "HpARI and HpTGMs")
dev.off()

```

```{r}
# Investigate aging
# C. elegans is an established model for studying aging
# Import list of C. elegans genes known to be involved in aging
celegans_aging_genes <- read.table("celegans_aging_genes.csv", quote="\"", comment.char="")

# Cross reference with H. bakeri - C. elegans orthos to find H. bakeri genes
hp_agingorthos=merge(celegans_aging_genes, orthopairs, by.x="V1", by.y="Caenorhabditis.elegans..PRJNA13758...WS279..gene.name", all.x=T, all.y=F)
# Only keep unique HPOLs
hp_agingorthos_uniqs=hp_agingorthos[!duplicated(hp_agingorthos$Gene.stable.ID),]

# Get expression pattern of the genes
indplot <- function(g, celename) {
  query=paste("transcript:", g, "-mRNA-1", sep="")
  tmp=vst_exprs_df[rownames(vst_exprs_df) == query,]
  tmp2=tmp[,c(20:23,28:31,1:7,12:15)] # Females
  tmp3=tmp[,c(24:27,32:35,8:11,16:19)] #Males
  
  plot(0.5, tmp2[1], ylim=c(7,16), xlim=c(0,35), xaxt='n', ylab="VST normalized expression", xlab=NA,  main=paste(g, " (", celename, ")", sep=""))
  abline(v=c(0,4,8,15,20,24,28,32), col="grey")
  points(seq(0.5,18.5), tmp2)
  points(seq(20.5,35.5), tmp3, pch=15)
  axis(1, at=c(0,4,8,15,20,24,28,32), labels=FALSE)
  text(c(3,7,11,17,23,27,31,35), labels=c("D5F", "D7F", "D10F", "D21F", "D5M", "D7M", "D10M", "D21M"), srt=35, cex=1.2, y=par("usr")[3]-1.5, xpd=NA, adj=0.99)
  
}

png(file="hbakeri_celeagingorthos_expr.png", width=4500, height=5000, res=300)
par(mar=c(2,1,1,0))
par(mfrow = c(6,3))
par(mar=c(5,3,2,0))
for (gene in 1:length(hp_agingorthos_uniqs$Gene.stable.ID)) {
  if(is.na(hp_agingorthos_uniqs$Gene.stable.ID[gene]) == FALSE) {
    indplot(hp_agingorthos_uniqs$Gene.stable.ID[gene], hp_agingorthos_uniqs$V1[gene])
  }
}
dev.off()

gs=c("daf-2", "age-1", "daf-16", "aak-2")
dat=vector()
for (g in gs) {
  d=hp_agingorthos_uniqs[hp_agingorthos_uniqs$V1 == g,]
  dat=rbind(dat, d)
}

png(file="hbakeri_celeagingorthos_expr_main4.png", width=2500, height=5000, res=300)
par(mar=c(2,1,1,0))
par(mfrow = c(4,1))
par(mar=c(5,3,3,0))
for (gene in 1:length(dat$Gene.stable.ID)) {
  if(is.na(dat$Gene.stable.ID[gene]) == FALSE) {
    indplot(dat$Gene.stable.ID[gene], dat$V1[gene])
  }
}
dev.off()

# No obvious patterns for 'aging' investigation. Don't have enough time points to really get into patterns of interest :(

```

```{r}
# Write the tables and supplementary tables to file
# Supp ms
ms_table=ms_withdescs[,c(1,2,4,8,10,14,16,20,22,26,27,28,38)]
rownames(ms_table)=substr(ms_table$...1, 12, nchar(ms_table$...1))
ms_table=ms_table[,-2]
write.csv(ms_table, "ms_transcripts.csv")

# Main ms/fs
msfs=ms_table[ms_table$Gene_Description != "",]
fstoadd=fs_withdescs[,c(1,2,4,8,10,14,16,20,22,26,27,28,38)]
rownames(fstoadd)=substr(fstoadd$...1, 12, nchar(fstoadd$...1))
fstoadd=fstoadd[,-2]
msfs=rbind(fstoadd, msfs)
write.csv(msfs, "msfs_transcripts_withannots.csv")

# Supp urm
urm=upm_b[,c(1,29,2:27)]
urm=merge(urm, hbakeri_annots_nondup, by.x="Gene_ID", by.y="GeneID", all.x=T)
rownames(urm)=substr(urm$...1, 12, nchar(urm$...1))
urm=urm[,c(1,4,8,10,14,16,20,22,26,27,28,38)]
write.csv(urm, "urm_transcripts.csv")

# Supp urm with o2o
urm_o2o=uniq_upm_o2o_withdesc[,c(1,2,4,8,10,14,16,20,22,26,27,28,33,31)]
rownames(urm_o2o)=substr(urm_o2o$...1, 12, nchar(urm_o2o$...1))
urm_o2o=urm_o2o[,-2]
write.csv(urm_o2o, "urm_transcripts_witho2ocele.csv")

# Function to turn GO results parents column into text for export
convertGOResP <- function(rearranged_results_obj) {
  rearranged_results_obj$parents=as.character(rearranged_results_obj$parents)
  rearranged_results_obj$parents=gsub(',', ';', rearranged_results_obj$parents)
  rearranged_results_obj$parents=gsub('c', '', rearranged_results_obj$parents)
  rearranged_results_obj$parents=gsub('"', '', rearranged_results_obj$parents)
  rearranged_results_obj$parents=gsub('[(]', '', rearranged_results_obj$parents)
  rearranged_results_obj$parents=gsub('[)]', '', rearranged_results_obj$parents)
}

# Supp urm GOs
urm_fe=upm_fe$result[,c(11, 10, 2:9, 12:14)]
urm_fe$parents=convertGOResP(urm_fe)
write.csv(urm_fe, "urm_fe_terms.csv")

# Supp urf
urf=upf_b[,c(1,29,2:27)]
urf=merge(urf, hbakeri_annots_nondup, by.x="Gene_ID", by.y="GeneID", all.x=T)
rownames(urf)=substr(urf$...1, 12, nchar(urf$...1))
urf=urf[,c(1,4,8,10,14,16,20,22,26,27,28,38)]
write.csv(urf, "urf_transcripts.csv")

# Supp urf with o2o
urf_o2o=uniq_upf_o2o_withdesc[,c(1,2,4,8,10,14,16,20,22,26,27,28,33,31)]
rownames(urf_o2o)=substr(urf_o2o$...1, 12, nchar(urf_o2o$...1))
urf_o2o=urf_o2o[,-2]
write.csv(urf_o2o, "urf_transcripts_witho2ocele.csv")

# Supp urf GOs
urf_fe=upf_fe$result[,c(11, 10, 2:9, 12:14)]
urf_fe$parents=convertGOResP(urf_fe)
urf_fe$parents[1]="" # This one was blank but then became 'harater0' based on the changes above
write.csv(urf_fe, "urf_fe_terms.csv")

# Supp urtdm
urtdm=dlvstm_coexpboth_sigrel[,c(2,6,8,12,14,18,20,24,25,26)]
urtdm$t=rownames(urtdm)
urtdm$Gene_ID=substr(rownames(urtdm), 12, 26)
urtdm=merge(urtdm, hbakeri_annots_nondup, by.x="Gene_ID", by.y="GeneID", all.x=T)
rownames(urtdm)=substr(urtdm$t, 12, nchar(urtdm$t))
urtdm=urtdm[,-12:-21]
write.csv(urtdm, "urtdm_transcripts.csv")

# Supp urtdm with o2o
urtdm_o2o=urtdm
urtdm_o2o$t=rownames(urtdm_o2o)
urtdm_o2o=merge(urtdm_o2o, orthopairs, by.x="Gene_ID", by.y="Gene.stable.ID", all.x=T)
urtdm_o2o=urtdm_o2o[urtdm_o2o$Homology.type == "ortholog_one2one" & is.na(urtdm_o2o$Homology.type) == FALSE,]
urtdm_o2o=urtdm_o2o[!duplicated(urtdm_o2o$Gene_ID),]
rownames(urtdm_o2o)=urtdm_o2o$t
urtdm_o2o=urtdm_o2o[,c(1:11,17,15)]
write.csv(urtdm_o2o, "urtdm_transcripts_witho2ocele.csv")

#Supp urtdm GOs
urtdm_fe=dlvstm_fe$result[,c(11,10,2:9,12:14)]
urtdm_fe$parents=convertGOResP(urtdm_fe)
write.csv(urtdm_fe, "urtdm_fe_terms.csv")

# Supp urtdf
urtdf=dlvstf_coexpboth_sigrel[,c(2,6,8,12,14,18,20,24,25,26)]
urtdf$t=rownames(urtdf)
urtdf$Gene_ID=substr(rownames(urtdf), 12, 26)
urtdf=merge(urtdf, hbakeri_annots_nondup, by.x="Gene_ID", by.y="GeneID", all.x=T)
rownames(urtdf)=substr(urtdf$t, 12, nchar(urtdf$t))
urtdf=urtdf[,-12:-21]
write.csv(urtdf, "urtdf_transcripts.csv")

# Supp urtdf with o2o
urtdf_o2o=urtdf
urtdf_o2o$t=rownames(urtdf_o2o)
urtdf_o2o=merge(urtdf_o2o, orthopairs, by.x="Gene_ID", by.y="Gene.stable.ID", all.x=T)
urtdf_o2o=urtdf_o2o[urtdf_o2o$Homology.type == "ortholog_one2one" & is.na(urtdf_o2o$Homology.type) == FALSE,]
urtdf_o2o=urtdf_o2o[!duplicated(urtdf_o2o$Gene_ID),]
rownames(urtdf_o2o)=urtdf_o2o$t
urtdf_o2o=urtdf_o2o[,c(1:11, 17,15)]
write.csv(urtdf_o2o, "urtdf_transcripts_witho2ocele.csv")

# Supp urtdf GOs
urtdf_fe=dlvstf_fe$result[,c(11,10,2:9,12:14)]
urtdf_fe$parents=convertGOResP(urtdf_fe)
write.csv(urtdf_fe, "urtdf_fe_terms.csv")

# Supp tissue shared
ts=tds_transcripts[,c(1,2)]
ts$t=rownames(ts)
ts$Gene_ID=substr(rownames(ts), 12, 26)
ts=merge(ts, hbakeri_annots_nondup, by.x="Gene_ID", by.y="GeneID", all.x=T)
rownames(ts)=substr(ts$t, 12, nchar(ts$t))
ts=ts[,-2:-13]
write.csv(ts, "tissue_shared_transcripts.csv")

# Supp urldm
urldm=ilvstm_coexpboth_sigrel[,c(2,6,8,12,14,18,20,24,25,26)]
urldm$t=rownames(urldm)
urldm$Gene_ID=substr(rownames(urldm), 12, 26)
urldm=merge(urldm, hbakeri_annots_nondup, by.x="Gene_ID", by.y="GeneID", all.x=T)
rownames(urldm)=substr(urldm$t, 12, nchar(urldm$t))
urldm=urldm[,-12:-21]
write.csv(urldm, "urldm_transcripts.csv")

# Supp urldm with o2o
urldm_o2o=urldm
urldm_o2o$t=rownames(urldm_o2o)
urldm_o2o=merge(urldm_o2o, orthopairs, by.x="Gene_ID", by.y="Gene.stable.ID", all.x=T)
urldm_o2o=urldm_o2o[urldm_o2o$Homology.type == "ortholog_one2one" & is.na(urldm_o2o$Homology.type) == FALSE,]
urldm_o2o=urldm_o2o[!duplicated(urldm_o2o$Gene_ID),]
rownames(urldm_o2o)=urldm_o2o$t
urldm_o2o=urldm_o2o[,c(1:11, 17, 15)]
write.csv(urldm_o2o, "urldm_transcripts_witho2ocele.csv")

# Supp urldm GOs
urldm_fe=ilvstm_fe$result[,c(11,10,2:9,12:14)]
urldm_fe$parents=convertGOResP(urldm_fe)
write.csv(urldm_fe, "urldm_fe_terms.csv")

# Supp urldf
urldf=ilvstf_coexpboth_sigrel[,c(2,6,8,12,14,18,20,24,25,26)]
urldf$t=rownames(urldf)
urldf$Gene_ID=substr(rownames(urldf), 12, 26)
urldf=merge(urldf, hbakeri_annots_nondup, by.x="Gene_ID", by.y="GeneID", all.x=T)
rownames(urldf)=substr(urldf$t, 12, nchar(urldf$t))
urldf=urldf[,-12:-21]
write.csv(urldf, "urldf_transcripts.csv")

# Supp urldf with o2o
urldf_o2o=urldf
urldf_o2o$t=rownames(urldf_o2o)
urldf_o2o=merge(urldf_o2o, orthopairs, by.x="Gene_ID", by.y="Gene.stable.ID", all.x=T)
urldf_o2o=urldf_o2o[urldf_o2o$Homology.type == "ortholog_one2one" & is.na(urldf_o2o$Homology.type) == FALSE,]
urldf_o2o=urldf_o2o[!duplicated(urldf_o2o$Gene_ID),]
rownames(urldf_o2o)=urldf_o2o$t
urldf_o2o=urldf_o2o[,c(1:11, 17, 15)]
write.csv(urldf_o2o, "urldf_transcripts_witho2ocele.csv")

# Supp urldf GOs
urldf_fe=ilvstf_fe$result[,c(11,10,2:9,12:14)]
urldf_fe$parents=convertGOResP(urldf_fe)
urldf_fe$parents[match("harater0", urldf_fe$parents)] = "" # There's an element that should be blank but became harater0 based on the manipulations in convertGOResP
write.csv(urldf_fe, "urldf_fe_terms.csv")

# Supp lumen shared
ls=lds_transcripts[,c(1,2)]
ls$t=rownames(ls)
ls$Gene_ID=substr(rownames(ls), 12, 26)
ls=merge(ls, hbakeri_annots_nondup, by.x="Gene_ID", by.y="GeneID", all.x=T)
rownames(ls)=substr(ls$t, 12, nchar(ls$t))
ls=ls[,-2:-13]
write.csv(ls, "lumen_shared_transcripts.csv")

# Supp tdshared GOs
tds_fe=tdshare_fe$result[,c(11,10,2:9,12:14)]
tds_fe$parents=convertGOResP(tds_fe)
write.csv(tds_fe, "tdshared_fe_terms.csv")

# Supp ldshared GOs
lds_fe=ldshare_fe$result[,c(11,10,2:9,12:14)]
lds_fe$parents=convertGOResP(lds_fe)
write.csv(lds_fe, "ldshared_fe_terms.csv")

# Supp DESeq2 results table
tmp=as.data.frame(res_fc_secondstranded[1])
tmp$rownames1=rownames(tmp)
for (i in 2:length(res_fc_secondstranded)) {
  tmp2=as.data.frame(res_fc_secondstranded[i])
  tmp2$rownames=rownames(tmp2)
  tmp=merge(tmp, tmp2, by.x="rownames1", by.y="rownames", all.x=T, all.y=T)
}
rownames(tmp)=substr(tmp$rownames1, 12, 100)
tmp=tmp[,-1]
write.csv(tmp, "deseq2results_allpccomps.csv")

alllists=list()
alllists[[1]]=ms_table
alllists[[2]]=fstoadd
alllists[[3]]=urm
alllists[[4]]=urf
alllists[[5]]=urtdm
alllists[[6]]=urtdf
alllists[[7]]=ts
alllists[[8]]=urldm
alllists[[9]]=urldf
alllists[[10]]=ls
names(alllists)=c("MS", "FS", "URM", "URF", "URTDM", "URTDF", "TS", "URLDM", "URLDF", "LS")
saveRDS(alllists, "hbakeri_all_genesets.rds")

```
````
