## Supplementary material for "Transcriptional patterns of sexual dimorphism and in host developmental programs in the model parasitic nematode *Heligmosomoides bakeri*": S2 File: Hbakeri_volcanoplots.R

```
library(EnhancedVolcano)

res_fc_secondstranded = readRDS("Hbakeri_deseq2_3-21-22_res_fc_secondstranded.rds")
genesets = readRDS("hbakeri_all_genesets.rds")

gens=c('transcript:HPOL_0000308001-mRNA-1', 'transcript:HPOL_0000740701-mRNA-1', 'transcript:HPOL_0001902701-mRNA-1', 'transcript:HPOL_0001264601-mRNA-1', 'transcript:HPOL_0001830501-mRNA-1', 'transcript:HPOL_0000100601-mRNA-1')

dvp <- function(g, comp, tit) {
  hc='#DE1E09'
  urmc='#52D6F7'
  urfc='#00A843'
  sc='lightgrey'
  nsc='black'
  
  keyvals <- ifelse(rownames(comp) %in% g, hc, ifelse(rownames(comp) %in% paste("transcript:", rownames(genesets$URM), sep=""), urmc, ifelse(rownames(comp) %in% paste("transcript:", rownames(genesets$URF), sep=""), urfc, ifelse(comp$padj < 0.05, sc, nsc))))
  
  keyvals[is.na(keyvals)] <- nsc
  names(keyvals)[keyvals == nsc] <- 'Not Significant'
  names(keyvals)[keyvals == sc] <- 'padj < 0.05'
  names(keyvals)[keyvals == urmc] <- 'URM'
  names(keyvals)[keyvals == urfc] <- 'URF'
  names(keyvals)[keyvals == hc] <- 'Marker Genes'
  
  dat=comp[which(names(keyvals) %in% c("Marker Genes")),c(2,6)]
  dat=as.data.frame(dat)
  dat$y=-log10(dat$padj)
  
  urm=comp[which(names(keyvals) %in% c("URM")),c(2,6)]
  urm=as.data.frame(urm)
  urm$y=-log10(urm$padj)
  
  urf=comp[which(names(keyvals) %in% c("URF")),c(2,6)]
  urf=as.data.frame(urf)
  urf$y=-log10(urf$padj)
  
  p=EnhancedVolcano(comp, x='log2FoldChange', y='padj', lab=rownames(comp), FCcutoff=0, pCutoff=0.05, colCustom = keyvals, colAlpha = 1, selectLab="", title=tit, xlim=c(-10,10), ylim=c(0,300))
  
  p = p + geom_point(data=urm, aes(x=log2FoldChange, y=y), alpha=1, colour=urmc)
  p = p + geom_point(data=urf, aes(x=log2FoldChange, y=y), alpha=1, colour=urfc)
  p = p + geom_point(data=dat, aes(x=log2FoldChange, y=y), alpha=1, colour=hc, size=4)
  
  return(p)
}

# Make 4 panel volcano plots for M / F comps
d5=dvp(gens, res_fc_secondstranded$D5_Female_vs_D5_Male, "D5 M/F")
d7=dvp(gens, res_fc_secondstranded$D7_Female_vs_D7_Male, "D7 M/F")
d10=dvp(gens, res_fc_secondstranded$D10_Female_vs_D10_Male, "D10 M/F")
d21=dvp(gens, res_fc_secondstranded$D21_Female_vs_D21_Male, "D21 M/F")

png(file="hbakeri_d5volcano.png", width=5000, height=2500, res=300)
d5
dev.off()

png(file="hbakeri_d7volcano.png", width=5000, height=2500, res=300)
d7
dev.off()

png(file="hbakeri_d10volcano.png", width=5000, height=2500, res=300)
d10
dev.off()

png(file="hbakeri_d21volcano.png", width=5000, height=2500, res=300)
d21
dev.off()
```
