## Supplementary material for "Transcriptional patterns of sexual dimorphism and in host developmental programs in the model parasitic nematode *Heligmosomoides bakeri*": S2 File: starbwa_scatter.Rmd

````
---
title: "starbwa_scatter"
author: "Stephen Pollo"
date: "19/04/2022"
output: html_document
---

```{r}
dat=readRDS("star_vs_bwa_mappingperecents.rds") # Generated from S1 Table and Table 1 with random x coordinate added

png(file="star_vs_bwa.png", width=2000, height=1800, res=300)
plot(dat$rand.x, dat$STAR, xlim=c(0,40), ylim=c(80,100), ylab="Percent of reads mapping (%)", xlab="Sample", pch=15, xaxt='n')
points(dat$rand.x, dat$bwa, pch=1)
legend("bottomright", legend = c("STAR", "bwa"), box.lty=0, pch=c(15, 1))
dev.off()
```
````
