## Supplementary material for "Transcriptional patterns of sexual dimorphism and in host developmental programs in the model parasitic nematode *Heligmosomoides bakeri*": S1 Text

**S1 Text. Supplementary Results and Discussion**

Bulk RNA-seq dataset QC

After mapping the RNA-seq datasets to the *H. bakeri* genome using STAR, reads overlapping each transcript were counted (See main Methods). The read counts per transcript were used to perform principal component analysis and hierarchical clustering (S1 Fig and S2 Fig). Biplots of the first two principal components are shown for either the raw counts (S1 Fig A, accounting for 86.3% of the variation) or the VST-transformed counts from DESeq2 (S1 Fig B, accounting for 83.7% of the variation). In both cases, all replicates of each sample condition cluster together and sample conditions cluster relative to each other in an expected way (ie. Male and female samples cluster separately, lumen-dwelling and tissue-dwelling samples cluster separately). These patterns indicate our samples are suitable for further comparison.

Hierarchical clustering using either Euclidean (S2 Fig A), Manhattan (S2 Fig B), or Canberra (S2 Fig C) distance shows sample replicates clustering together with sample groups tending to cluster by sex rather than by age (ex. lumen females cluster together rather than D21 worms clustering together). This pattern matches that seen in the principal component biplots and further indicates our samples are what we intended and are suitable for further comparison.

*H. bakeri* PRJEB15396 Annotations

To evaluate the completeness and accuracy of the genome annotations, the RNA-seq reads were aligned to the transcript sequences predicted from the genome annotation that accompanies the PRJEB15396 assembly using the non-splice-aware aligner bwa (S1 Table). Among all the datasets, 83.70 – 89.41% of the reads mapped to the predicted transcript sequences, suggesting that the annotations have captured the bulk of the polyadenylated transcript content, but there is still room for improvement. In comparison to the mapping rates to the genome, the lower mapping rates to the predicted transcripts point to the annotations missing regions that are actually transcribed. Moreover, the rates of mapped singletons, mismatched pairs, and misoriented pairs point to occasional missing exons and/or untranslated regions among the predicted transcripts.

Samples included in pairwise comparisons

Of the original 32 samples sequenced (four for each timepoint/sex combination examined in the study), two stood out in preliminary quality control analyses as problematic. D10 F6 and D10 F7 both clustered with D10 M samples in PCA biplots and hierarchical clustering dendrograms (data not shown). Moreover, the most abundant transcript in these samples is a major sperm protein (HPOL_0001609601), as is the case for the adult male samples. Furthermore, expression of orthologs of transcriptionally regulated male patterning genes in *C. elegans* mab-3 (HPOL_0001902701) and mab-23 (HPOL_0000964001) in these two samples match the levels seen in the other male samples. We therefore sequenced an additional five D10 F samples and sequenced two technical replicates (D10 M3 and D10 M6) to ensure no batch effects were seen between sequencing runs. No batch effects were observed in PCA analyses (data not shown) so the seven D10 F samples that clustered together (and near the D21 F samples) were used in all analyses. The D10 F6 and D10 F7 samples were excluded from all analyses except the assessment of the genome assembly and annotation.

Intestinal oxygen in mice

While air at sea level has a partial pressure of oxygen (pO_2_) of ~145 mmHg (~21% O_2_), pO_2_ varies considerably throughout the different tissues in mammalian bodies [1]. Similarly to humans, the mouse gastrointestinal (GI) tract has a pO_2_ gradient along the proximal-distal axis and along the radial axis [1]. Much research focuses on the distal GI tract and the anaerobic colon, however, *H. bakeri* larvae encyst in the wall of the duodenum in the small intestine and stay as adults in the lumen of the duodenum. Initial measurements in mice using spatial and spectral electron paramagnetic resonance (EPR) imaging placed the pO_2_ of the lumen of the duodenum at 32 ± 8 mmHg [2]. A second set of measurements in mice using phosphorescence quenching placed the pO_2_ of the lumen of the stomach at ~22 mmHg, the stomach tissue at ~27 mmHg, the lumen of the duodenum at ~61 mmHg, and the tissue of the duodenum at ~45 mmHg [3]. In this study it was noted that the measurement for the lumen of the duodenum required further investigation due to the possibility of small molecule quenchers in the pancreatic/bile secretions that could have given an artificially high measurement.

Sources of oxygen in the lumen of the duodenum include:1) oxygen still in the material entering from the stomach, 2) oxygen diffusing from the duodenum tissue, and 3) oxygen in the pancreatic and bile secretions entering the duodenum [3]. The measures of the first two sources were well below the 61 mmHg measurement in question. In the periportal area of the human liver (where fresh oxygenated blood enters), the pO_2_ is 65 mmHg [4]. Assuming a similar level in the mouse liver, bile in secretions would not be sufficient to boost oxygen concentration to higher than the surrounding tissue unless those secretions made up the vast majority of the content in the lumen. The alternative measurement of 32 mmHg is therefore a more reasonable estimate of the pO_2_ of the duodenum lumen. This level is generally considered ‘physiologically hypoxic’ because in addition to being a low pO_2_, the epithelial cells lining the GI tract lumen display typical hypoxia responses that are regulated by the transcription factor hypoxia-inducible factor [1].

Aging-related gene expression

To identify transcripts in *H. bakeri* with potential roles in aging we sought to find all the transcripts with expression patterns matching the orthologs of genes in *C. elegans* that have already been implicated in aging. We identified an initial 18 candidate orthologs (See table below), whose expression is plotted in S3 Fig. In addition to many sex-specific expression patterns, the 18 candidate orthologs show a varied assortment of expression patterns including: 1) increasing with age (ex. dcap-2 males), 2) decreasing with age (ex. eat-2 males), 3) changing upon final molt (ex. aak-2 females), 4) spiking at a certain time point (ex. daf-7 males), and 5) not changing (ex. skn-1 males). Identifying transcripts whose expression mirrors a candidate may identify potential interacting partners of the candidate gene product. However, in the case of the process of aging, all potential expression patterns were found among genes already implicated in the process so new candidates could not be identified based on expression pattern alone. Additional time points may refine expression patterns of interest, but without the ability to follow up on new candidate genes with the use of mutants or RNAi it is unlikely true involvement in aging can be confirmed in this worm.

| ***C. elegans* Gene Implicated in Aging** | **Reference** |
| --- | --- |
| daf-2 | [5] |
| age-1 | [5] |
| daf-16 | [5] |
| skn-1 | [5] |
| aak-2 | [5] |
| hsf-1 | [5] |
| eat-2 | [5] |
| dcr-1 | [6] |
| alg-1 | [6] |
| dcap-1 | [6] |
| dcap-2 | [6] |
| xrn-1 | [6] |
| jnk-1 | [5] |
| cst-1 | [5] |
| daf-7 | [5] |
| pqm-1 | [5] |
| sir-2.1 | [5] |
| sod-2 | [5] |

Immunomodulatory gene expression

Expression of immunomodulatory genes that could be identified is described in the main text. HbTGM1-3 were excluded from this analysis because the locus in the PRJEB15396 that was previously associated with these proteins was incorrectly assigned. The locus does not match the length of the reported protein(s), nor does the sequence from that locus match the sequences associated with these proteins. Analysis of additional unidentified immunomodulatory proteins was attempted based on gene annotations but cannot be linked to any of the peptides detected in analyses of secreted products because the information needed to do so was never made available.
